# Combinatorial protein barcodes enable self-correcting neuron tracing with nanoscale molecular context

**DOI:** 10.1101/2025.09.26.678648

**Authors:** Sung Yun Park, Arlo Sheridan, Bobae An, Erin Jarvis, Julia Lyudchik, William Patton, Jun Y. Axup, Stephanie W. Chan, Hugo G.J. Damstra, Daniel Leible, Kylie S. Leung, Clarence A. Magno, Aashir Meeran, Julia M. Michalska, Franz Rieger, Claire Wang, Michelle Wu, George M. Church, Jan Funke, Todd Huffman, Kathleen G.C. Leeper, Sven Truckenbrodt, Johan Winnubst, Joergen M.R. Kornfeld, Edward S. Boyden, Samuel G. Rodriques, Andrew C. Payne

## Abstract

Mapping nanoscale neuronal morphology with molecular annotations is critical for understanding healthy and dysfunctional brain circuits. Current methods are constrained by image segmentation errors and by sample defects (e.g., signal gaps, section loss). Genetic strategies promise to overcome these challenges by using easily distinguishable cell identity labels. However, multicolor approaches are spectrally limited in diversity, whereas nucleic acid barcoding lacks a cell-filling morphology signal for segmentation. Here, we introduce PRISM (Protein-barcode Reconstruction via Iterative Staining with Molecular annotations), a platform that integrates combinatorial delivery of antigenically distinct, cell-filling proteins with tissue expansion, multi-cycle imaging, barcode-augmented reconstruction, and molecular annotation. Protein barcodes increase label diversity by *>*750-fold over multicolor labeling and enable morphology reconstruction with intrinsic error correction. We acquired a ∼10 million µm^3^ volume of mouse hippocampal area CA2/3, multiplexed across 23 barcode antigen and synaptic marker channels. By combining barcodes with shape information, we achieve an 8x increase in automatic tracing accuracy of genetically labelled neurons. We demonstrate PRISM supports automatic proofreading across micron-scale spatial gaps and reconnects neurites across discontinuities spanning hundreds of microns. Using PRISM’s molecular annotation capability, we map the distribution of synapses onto traced neural morphology, characterizing challenging synaptic structures such as thorny excrescences (TEs), and discovering a size correlation among spatially proximal TEs on the same dendrite. PRISM thus supports self-correcting neuron reconstruction with molecular context.

## 1 Introduction

Information flow in the mammalian brain is governed by neuronal morphology, with function shaped both by where synapses are distributed on the cell and by the molecular composition of synapses. Disruptions in circuit organization and synaptic machinery have been implicated in diverse disorders such as autism, Parkinson’s disease, and epilepsy (Lepeta et al., 2016). A robust, self-correcting method for neural tracing with molecular annotation would therefore benefit diverse areas of neuroscience.

Electron microscopy (EM) has long been the primary method for mapping neuron morphology, supplying dense ultrastructural detail for tracing. Despite its power for mapping morphology at scale in mammals (Bae et al., 2025; Schneider-Mizell et al., 2025; Shapson-Coe et al., 2024) and other organisms, scalable EM reconstruction faces two central limitations. First, it is sensitive to both within-slice signal discontinuities and between-slice serial section loss, where even small gaps can fragment a dataset into disjointed subvolumes (“Scaling up connectomics — Reports,” 2023). Second, scaling EM reconstruction remains prohibitively expensive because it requires extensive human proofreading of automatic segmentation outputs (Dorkenwald et al., 2024). For example, section loss in a recent cubic-millimeter EM dataset forced subdivisions of the volume to be reconstructed separately. Within the largest subvolume, a data release of approximately 1,500 proofread cells required more than one million human proofreading edits (Bae et al., 2025). Furthermore, EM cannot intrinsically visualize specific molecules, requiring correlative methods to capture this information (Begemann & Galic, 2016; Lam et al., 2015).

Light microscopy provides a complementary path to morphology reconstruction. Recent advances demonstrate how genetic (Gao et al., 2019) or protein density labeling (Damstra et al., 2023; M’Saad et al., 2022; Tavakoli et al., 2025), combined with tissue clearing and expansion microscopy (F. Chen et al., 2015), can provide neuronal reconstructions from light microscopy data with nanoscale detail. Importantly, combining these methods with the labeling of endogenous markers and proteins allows for the addition of key biological information to the reconstructed circuits (Shen et al., 2020). However, like EM, these approaches fundamentally rely on tracing the continuity of neurons through sections, making them sensitive to section loss. Tissue clearing approaches help address this by allowing for thicker sectioning (Winnubst et al., 2019) and even wholebrain imaging (Glaser et al., 2025). Still, serial sectioning and extensive proofreading are key bottlenecks to scaling up these approaches when high resolution is required (e.g. for synaptic analysis), or when applying such techniques to the diverse questions of everyday neuroscience. For example, imaging an entire mouse brain for light microscopy connectomics would involve tomography of thousands of sections, and proofreading the resulting volume would be cost-prohibitive (Kornfeld, 2025). Despite several connectomic triumphs (Bae et al., 2025; Dorkenwald et al., 2024; Shapson-Coe et al., 2024), that have mapped neural circuits with unprecedented detail, comprehensive studies of neuronal morphology - essential for linking structure to function within specific cellular and circuit contexts (Hardingham & Bading, 2010) - remain challenging and are still uncommon in routine neuroscience research.

Cellular barcoding approaches have the potential to overcome these scaling bottlenecks by providing each neuron with a molecular signature (i.e., a barcode) that can be used to correct tracing errors and bridge spatial gaps, thereby mitigating issues of serial loss and proofreading cost. Combinatorial labeling with spectrally-distinct fluorescent proteins simplifies tracing by increasing the number of distinguishable neurons (Cai et al., 2013; Livet et al., 2007; Sakaguchi et al., 2018), and can even reconnect neurons across discontinuities (Leiwe et al., 2024). However, this approach is constrained by a limited spectral budget of fluorophores and laser lines. RNA barcodes can distinguish many more neurons but, because they do not fill the cell, cannot be used for morphological reconstruction (X. Chen et al., 2019; Goodwin et al., 2022; Kebschull et al., 2016; Yuan et al., 2024).

We present PRISM, a technology platform enabling scalable, self-correcting morphology reconstruction with molecular annotation, designed to address key challenges of accurate neuron tracing and sample fragility. We developed a toolbox of highly diverse cell-filling protein labels with cognate antibodies, achieving barcode diversity 750-fold greater than conventional multicolor labeling. We co-optimized this barcoding system with expansion microscopy and rounds of sequential staining for nanoscale and multiplexed detection. We then built machine learning (ML) pipelines that incorporate barcode information into automatic image segmentation and automatic proofreading (Figure 1.C). We show the expected error-free tracing distance is 8x longer with PRISM than with conventional tracing based on a single-channel fluorescent protein (eGFP). We demonstrate that, by using protein barcodes, we can error-correct micron-scale discontinuities and cross spatial gaps of up to hundreds of microns. We used PRISM to map thousands of synaptic contacts across reconstructed neurons. Using the enhanced resolution of expansion microscopy, we resolve nanoscale features such as thorny excrescences (TEs) and identify a size correlation between spatially proximal TEs on the same dendrite.

**Figure 1.**
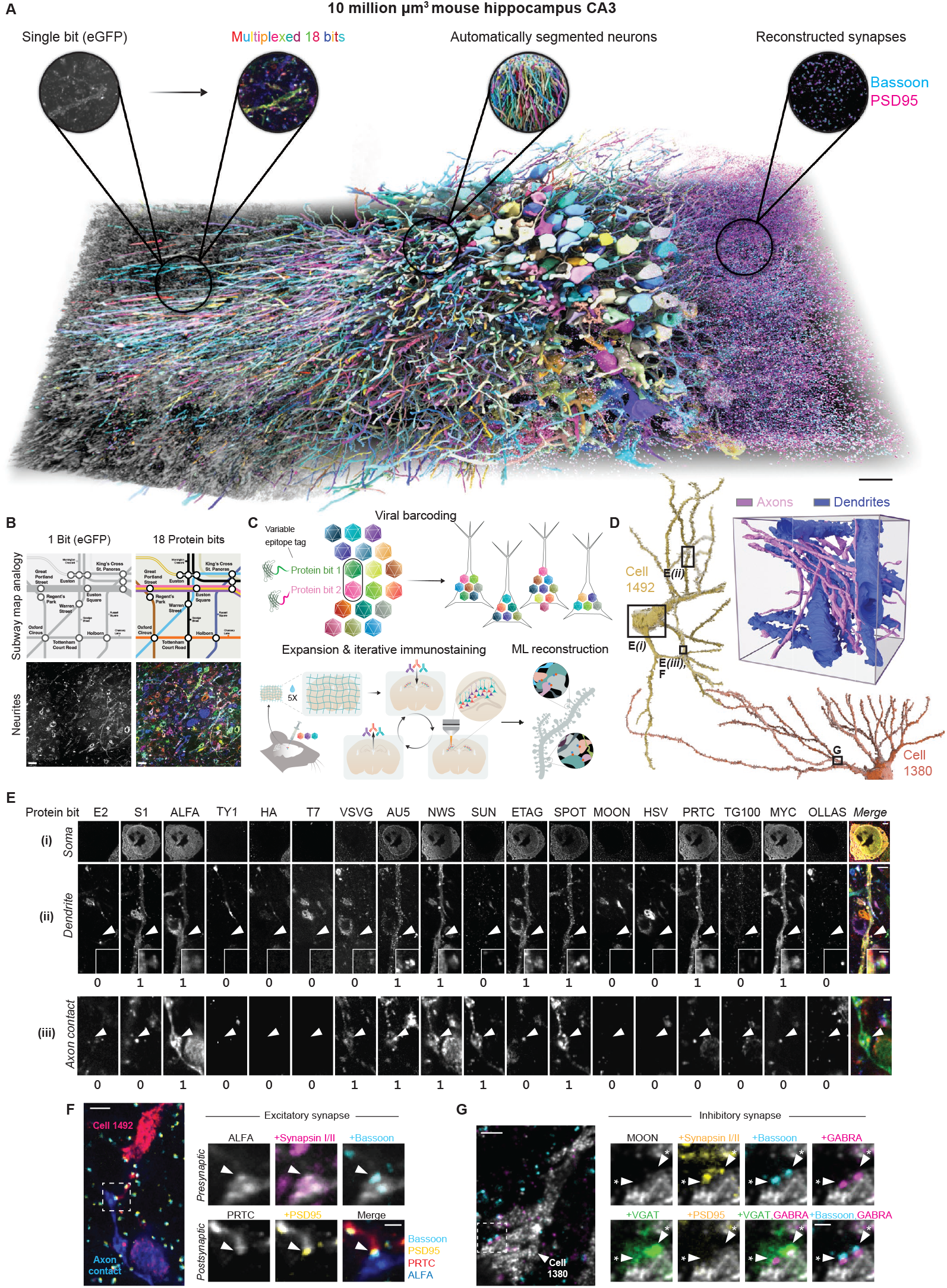
: PRISM enables multiplexing and reconstruction of sparsely labeled mouse hippocampal CA3 neurons. A. 3D rendering of a ∼10 million µm^3^ volume of mouse hippocampal CA3 with 18 protein bits, 5 synaptic markers, and a 35 × 35 × 80 nm voxel size. Panel highlights are four different views of the dataset (left to right): single-plane images containing single bit (eGFP) in grayscale, single-plane multi-colored images containing 18 protein bits, 3D segmentations using 18-bit protein barcodes, and synapse reconstructions of presynaptic active zone marker Bassoon and postsynaptic excitatory scaffold protein PSD95. Insets are demonstrative images for each of the four views and do not correspond to the circled region of interest (ROI). The full volume for morphology and synapses can be further explored in Movie S1. Scalebar: 10 µm. B. Subway map analogy of leveraging protein barcodes for automated segmentation. Top, view of the London Underground map showing multiple different tube lines and bottom, single-plane images of a neurite ROI in the hippocampal CA3 dataset. The left panels are both presented in single color while the right panels are presented with multiple channels or barcodes. Color enables distinguishing of overlapping objects. The single-color grayscale image of neurites is an anti-GFP stain. Scalebar: 20 µm. C. Overview of PRISM. (Top) Combinatorial protein barcoding via stochastic AAV infection. Stochastic infection leads to random expression of protein bits; representative structure shown on right. (Bottom left) Multiplexed antibody staining in 5X expanded barcoded brain slices. (Bottom right) ML-based reconstruction of neurons utilizing barcodes for automated proofreading and segmentation and molecular annotations for synapse reconstruction. Assortment of high-resolution 3D images of axons and dendrites as well as two human-proofread cells with ID #s 1492 and 1380. Boxes on cell 1492 and 1380 correspond to the approximate ROI for neuronal structures shown in the breakout panels in E, F, and G. PRISM data can be used to reconstruct dendrites and axons in the same volume, distinguishable by morphology. E. Example of barcode multiplexing in a soma and dendrite of the same cell, and axon contact from an adjacent cell. Multiplexing data shown for the soma and dendrite of cell ID 1492 (top panel) and a crossing barcoded axon (bottom panel). Single-plane raw images for each ROI are displayed for each protein bit channel, with binary protein bit assignments shown at the bottom of the corresponding panels. Insets of a dendritic spine (arrowhead) are displayed. Scale bars: 2 µm. F. Identification of an excitatory synapse between two barcoded neurites via synapse multiplexing. Left: Overview image of dendrite of cell 1492 contacting the axon shown in E, III (bottom row). Arrowhead marks the putative synapse. All images are single-plane. Scale bar: 5 µm. Right, top row, left to right: presynaptic neuron in ALFA; overlaid with Synapsin I/II; overlaid with Bassoon. Right, bottom row: postsynaptic neuron in PRTC channel; overlaid with PSD95; merged view showing both barcodes with Bassoon and PSD95. Scale bar: 2 µm. G. Identification of an inhibitory synapse on cell 1380. Left: Zoom out of the dendritic shaft receiving inhibitory input with GABRA, Bassoon, and MOON overlay. Scale bar: 5 µm. Right, 1.2 µm thick maximum intensity projection (MaxIP) images of synapse channels overlaid on the MOON protein bit channel. Arrow indicates putative inhibitory synapses. Top row, left to right: 1380 dendrite in the MOON channel, Synapsin I/II overlay, Bassoon overlay; GABRA overlay. Bottom row, left to right: VGAT overlay; PSD95 overlay; VGAT and GABRA overlay; Bassoon and GABRA overlay. Scale bar: 2 µm. All figure scale bars represent pre-expansion biological distances.

## 2 Results

### 2.1 Protein barcode design and delivery

We set out to design an ideal protein barcoding system, which should have high diversity approaching that of RNA barcodes (Kebschull et al., 2016) while labeling morphology such that it can be traced in 3D image volumes and coregistered with synaptic labeling. Informed by work leveraging combinatorial adeno-associated virus (AAV) infection for protein diversity (Cai et al., 2013; Sakaguchi et al., 2018), we chose to use an AAV delivery system, which offered both the high multiplicity of infection required for protein bit diversity and the delivery of cell-filling protein payloads. Drawing from work on multiplex imaging of epitope tags (Kudo et al., 2022; Rovira-Clavè et al., 2021; Wroblewska et al., 2018), we chose to focus on antigenic diversity over spectral diversity, reasoning that expanding the palette of peptide tags and corresponding antibodies would provide a far larger and more flexible design space than conventional fluorescent approaches.

To build this system, we chose enhanced green fluorescent protein (eGFP) as a scaffold protein well-characterized for safety and trafficking in neuronal tracing (Viswanathan et al., 2015). Based on the simple structure used in epitope tagging of native proteins, we fused tags to the C-terminal with a flexible GGSGGS linker to minimally disrupt the native folding and trafficking properties of eGFP. We expressed these protein bits *in vivo* using AAV and screened libraries of candidate epitope fusions and antibodies in expanded tissue. We worked with a MAGNIFY gel chemistry (Klimas et al., 2023) because of its robust gel chemistry and generalized protein retention, likely to be compatible with a wide range of protein bits. Epitope-binder pairs were qualitatively assessed for signal specificity and cross reactivity, compatibility with expansion chemistry, and expression in cell bodies and neurites. From this, we developed a set of top-performing antibodies for these protein bits, spanning multiple species with a set of 18 validated pairs (Supplementary Table 1), providing a foundation for a robust epitope palette for barcode generation.

Finally, to make the approach economically feasible, we produced AAV barcoding constructs as pooled libraries of plasmids rather than generating each virus individually. However, as with our initial antibody screens, evaluating how these pooled libraries functioned as complete barcodes required extensive multiplexed analyses. Having established a library of discrete protein bits and a delivery mechanism, we carried out a pilot study to examine their performance *in vivo*. This provided an opportunity to assess barcode diversity and representation across cells, and test the utility of protein barcodes for reconstruction.

### 2.2 Detection of virally-expressed protein barcodes and synaptic markers

We injected a PHP.eB-pseudotyped viral pool of 18 protein bits into the mouse hippocampal CA3 region, whose welldefined circuitry (Cherubini & Miles, 2015; Rebola et al., 2017; Sammons et al., 2024) makes it particularly suitable for circuit tracing (Figure 1.C). To achieve high-resolution morphology and identify putative connections, we used a modified MAGNIFY protocol with robust gel chemistry and protein retention to expand brain slices 5-fold, yielding a biological voxel size of ∼35 × 35 × 80 nm (x, y, z) (Klimas et al., 2023) (Figure 1.C; Supplementary Section 6.1).

To simultaneously detect protein barcode-labelled morphology and molecular annotations, we developed an iterative immunostaining protocol to overcome the limited spectral diversity of conventional fluorescence microscopy (Figure 1.C). This protocol incorporated two key elements: (1) preservation of antigenicity across multiple staining rounds using heated homogenization buffer for antibody stripping, and (2) improved imaging performance via a photoprotective glucose oxidase–catalase buffer under anoxic conditions, which maintained pH stability and minimized photoinduced crosslinking and epitope damage (Aitken et al., 2008; Cordes et al., 2009; Gut et al., 2018; Herdly et al., 2023; Kingsley et al., 2001). With this strategy, we achieved highly efficient signal extinction across protein bit channels, with median stripping efficiencies consistently above 93% (Supplementary Figure 2.A,B, Supplementary Section 6.1.3). Restaining stripped samples with secondary antibodies from the previous round showed minimal antibody crosstalk, with median change in signal-to-background ratio ranging from −0.97% to 8.06% across channels (Supplementary Figure 2.D, Supplementary Section 6.1.4), demonstrating the method supports extensive round by round multiplexing.

With this multiplexing strategy, we acquired a preexpansion volume of ∼10 million µm^3^ of mouse hippocampal CA3, detecting 18 protein barcodes and 5 synaptic markers across 8 stripping cycles and 12 rounds of imaging (Figure 1.A,E,F,G; Supplementary Figure 1; Supplementary Table 2; Movie S1; Movie S2). Despite repeated heat denaturation and exposure to high-powered lasers during imaging, all 18 protein bits were detectable in somas, dendrites, axons, and dendritic spines alongside 5 other synaptic markers (Figure 1.E,F,G, Supplementary Figure 1) demonstrating PRISM preserves sample integrity throughout multiplexing. Antigenicity was further supported by registering 11 imaging rounds using the ALFA protein bit, stripped and re-stained over seven consecutive cycles (Supplementary Table 2), with visual matching confirming consistent signal (Supplementary Figure 2.E). Regions where protein bit channels were stained with same-species primary and secondary antibodies across different rounds showed minimal signal overlap, suggesting low crosstalk optimal for reliable barcode assignment (Supplementary Figure 2.C).

Our strategy also supported multiplexed detection of synaptic markers. Due to fine spatial localizations and differing marker-specific needs for optimal immunodetection, detecting a range of endogenous pre-, post-, excitatory, and inhibitory synaptic markers has proven difficult to achieve within the same sample (Konno & Watanabe, 2021; Konno et al., 2023; Lorincz & Nusser, 2008; Melone et al., 2005; Schneider Gasser et al., 2006). Using our workflow, we detected the expected spatial correlations among the active zone protein Bassoon, pre-synaptic vesicle marker Synapsin I/II, inhibitory vesicular protein VGAT, and postsynaptic excitatory and inhibitory proteins PSD95 and GABRA1, even after 4–6 rounds of heat-stripping and imaging (Figure 1.F,G). This performance was likely aided by strong protein retention in the MAGNIFY gel, epitope decrowding from expansion, and antigen retrieval during homogenization and stripping before sample degradation became limiting (Klimas et al., 2023).

Together, these results establish a robust workflow for multiplexed detection of 23 protein targets (18 protein bits and 5 synaptic markers) across repeated stripping and imaging cycles in expanded tissue. This approach expands the scale and molecular detail achievable in a single brain volume at nanoscale resolution, as illustrated by our ∼ 10 million µm^3^ hippocampal dataset, providing a foundation for barcode analysis, morphological reconstruction, and synaptic profiling.

#### AAV-delivered protein barcodes

With a fully multiplexed dataset in hand, we next evaluated the efficiency and diversity of our barcoding system. We annotated the presence or absence of each protein bit in the somas of this dataset (Figure 2.A,B), focusing on somas to avoid potential confounds from branching neurites. Within the volume, we identified a total of 298 somas with 146 containing at least one protein bit (49%). Across these 146 labeled somas, each protein bit was present on average in 22% ± 5% of cells (range: 15%–36%; Figure 2.C) with a mean barcode length of 4 ± 2.5 bits per soma (Figure 2.D). Within this set, 80% of somas had a unique barcode, defined as a Hamming distance of ≥ 1 from their nearest neighbor (Figure 2.E). Somas averaged a Hamming distance of 6 ± 2.5 from other somas in the volume (Supplementary Figure 3.B). Barcode collisions were primarily associated with single-bit barcodes (70% of collisions; Supplementary Figure 3.A), and repeated barcodes were observed in fewer than four somas per repeat (Figure 2.E). These results suggest that in a modestly-sized barcoding dataset, the scheme produces a diverse set of barcodes with limited collisions, motivating further analysis of scalability through simulation of the barcodes.

**Figure 2.**
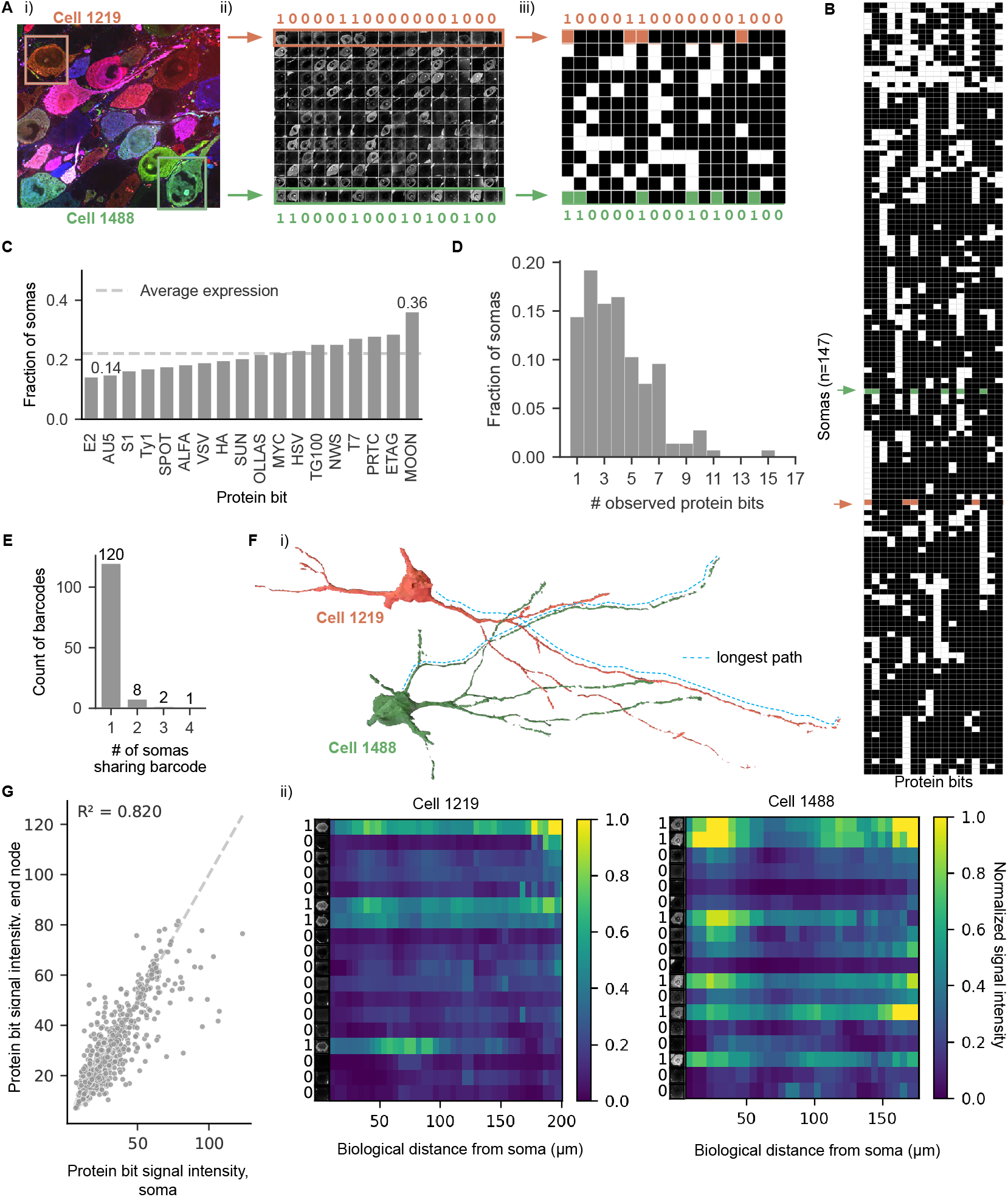
: Combinatorial protein barcode expression. A. Barcode bit assignment. Example of workflow for protein bit barcode assignment in a given FOV. i) Cell body selection ii) Channel breakout per soma with selected barcode assignments and iii) binarized soma calls after assignments. B. Binarized barcode calls for all 146 somas, clustered using Ward’s method. The largest silhouette score (Hamming) across k=2–10 was S=0.19 at k=2, consistent with the diversity and overlap expected from combinatorial label generation. Cells are used in C, D, E. C. Fraction of somas expressing each epitope. Fraction of each protein bit detected across 146 somas (mean = 0.22±0.05). D. Distribution of protein bit counts per soma. 85% of somas contain *>*1 protein bits (mean 4.2±2.4). E. Per barcode collision rate. For each barcode observed in the dataset, we calculated the number of cells containing that barcode. 120 barcodes appear only once in the dataset, with a total of 131 codes observed. F. Barcodes can be detected down the longest path of neurites. (i) Representative longest path intensities of example cells 1219 and 1488. (ii) Signal intensity from the soma down the longest path of cells 1219 and 1488. Distance is pre-expansion biological distance. Minimum and maximum values were set per channel for low background and high signal and correspond to the inset of somas shown on heatmap left. Average intensity was computed per skeleton node, and plotted as a fraction of the maximum value for that channel. G. Intensity correlation between cell body and end point of skeletonized somas (n= 74). Each point represents the values for a single channel measured at the start and end of a single cell. Across cells and channels, intensities are correlated with an *R*^2^ of 0.82.

To estimate how this diversity would scale across larger populations, we performed Monte Carlo simulations based on the observed barcode distributions (Supplementary Figure 3.D). All simulations were conducted under a strict binary model, considering only the presence or absence of each unique protein bit. For a typical stereotaxic injection labeling ∼ 10^6^ cells, the simulations predict ∼ 100,000 ± 200 unique binary barcodes (Supplementary Figure 3.D,F). To characterize the labeling efficacy of these barcodes, we defined a singlet rate (SR) metric, representing the estimated fraction of cells with unique labels. Applying this metric to both our observed distribution and several Poisson-ideal models, we found that our data provides a 64-fold improvement over current methods (SR of 50%: ∼ 80 cells 7 bits vs. ∼ 5,000, 18 bits) (Supplementary Figure 3.E,G), and up to a 2,000-fold improvement under ideal conditions of AAV infection and protein bit balance.

#### Barcode trafficking into neurites and fine structures

To establish the extent to which barcodes traffic into fine structures and down neurites, we performed expert semantic segmentation on a test crop to assess labeling of dendrites, axons, and spines. In this training crop, 26.81% of pixels were classified as dendrites and 4.96% as axons, confirming barcodes fill neurites and can be used to reconstruct fine structures.

We then evaluated barcode trafficking across neurites. We mapped signal intensity along the longest path of each neuron to qualitatively assess trafficking and signal stability along the neurite (Figure 2.F). Comparing signal intensity at the soma with the farthest point along each neuron, we observed a strong correlation across channels and cells (*R*^2^ = 0.82; n = 76; Figure 2.G), indicating that barcodes can support tracing neurites over long distances, including across gaps.

Together, these data demonstrate that AAV-delivered protein barcodes provide a scalable and high-diversity system with reliable, morphologically-filling neurite labeling and trafficking, achieving levels of practical uniqueness comparable to RNA- or DNA-based methods. The consistent labeling across neurites and diversity make these barcodes well-suited for computational morphological reconstruction.

### 2.3 Incorporating barcode information for automated neuronal segmentation

To automatically reconstruct neuronal morphologies in our dataset we employed an affinity graph partitioning approach, an established solution for efficient segmentation of large volumes. In an affinity graph, edges between pairs of pixels are assigned a weight (which can be learned by a network), denoting whether those pixels are connected or not (Turaga et al., 2010). The affinities can be subsequently agglomerated into unique segments using graph partitioning algorithms (Beier et al., 2017; Funke et al., 2019; Wolf et al., 2018). However, these methods have so far been designed only for grayscale images (as in EM). Due to the favorable properties of affinity graphs for large scale neuron segmentation, we extended these approaches to incorporate color information.

To investigate whether protein bit color information could be used to improve segmentation accuracy, we extended a previous multitask network that learns an affinity graph together with local shape descriptors (Sheridan et al., 2022). To evaluate each model we used a blockwise mutex watershed (Wolf et al., 2018) post-processing pipeline to generate segmentations masked to barcoded regions (Figure 3, Supplementary Figure 7, Supplementary Section 6.2.4). As a baseline, we used a model with a single channel input, specifically the GFP-scaffold channel that is common across all protein bits.

**Figure 3.**
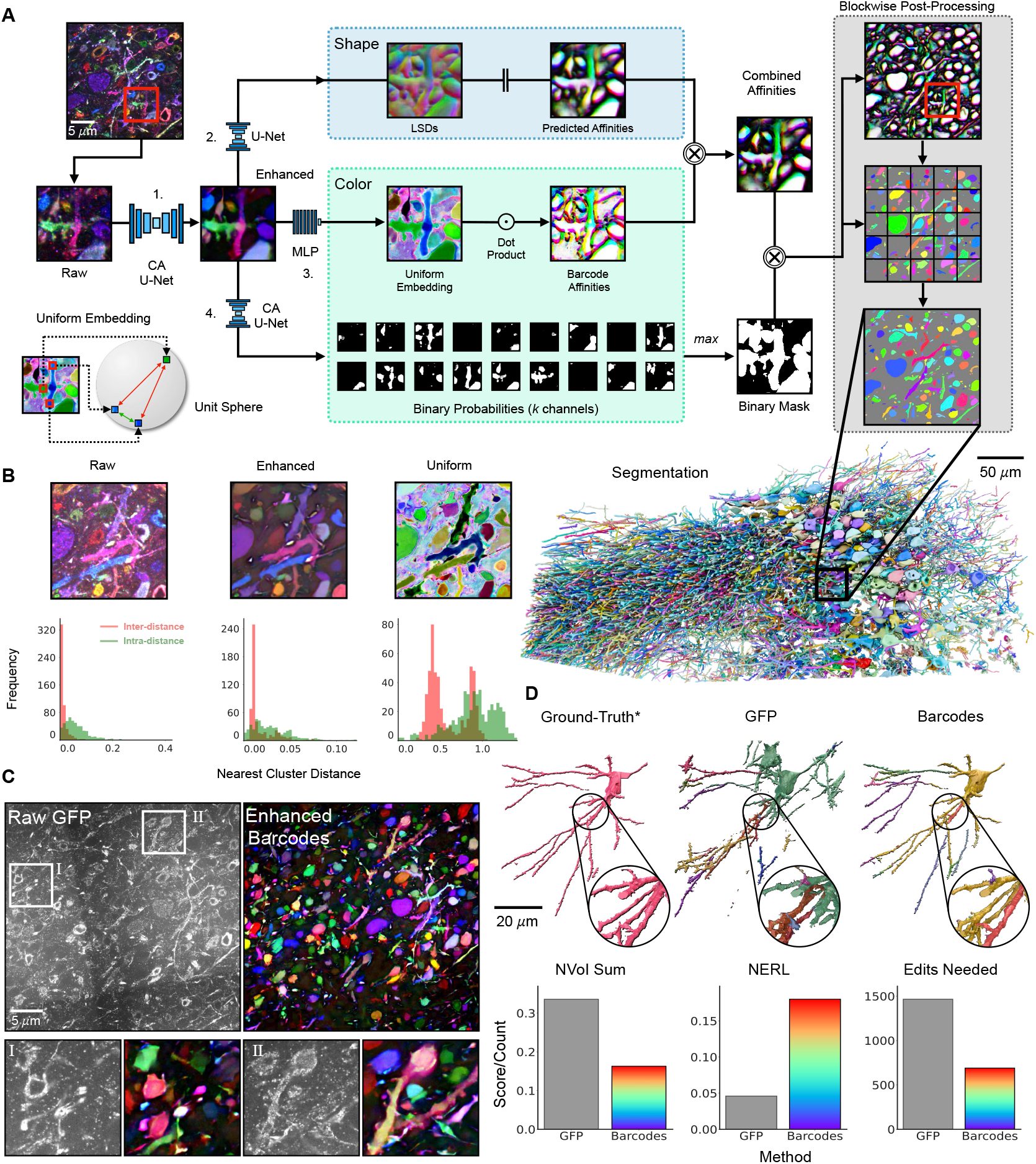
: Incorporating barcodes for automated neuron segmentation. A. Barcode assisted segmentation pipeline. Raw multi-channel barcode data (RGB-color encoded for visualization) is first enhanced by network 1. The resulting enhanced data is then used to learn affinities and LSDs (network 2), a uniform embedding (network 3), and binary expression probabilities (network 4). Barcode affinities are computed on the uniform embedding and then combined with the predicted affinities. Using a binary mask generated on the expression probabilities, a segmentation is then obtained in barcoded neurons in a blockwise fashion. Largest ∼ 5k segments visualized. B. Uniform embedding increases barcode separability. Enhancing the raw barcodes smooths the signal inside objects but does not provide an easy way to separate objects. The distributions of nearest cluster distances of each component (red: nearest true negative, green: furthest true positive) demonstrate that while none of the intensities are distributed such that they could be directly clustered to segments (minimum inter: high, maximum intra: low), the uniform embedding does provide increased separability (larger distribution of minimum inter-distances) which, in turn, allows for the direct computation of edge affinities via the dot product. C. Example image data. Single channel raw GFP data and multi-channel enhanced barcode data. D. Example meshes. Ground-truth mesh (skeleton constrained) and corresponding GFP and barcoded segmentation meshes (with ≥ 50% volumetric overlap). E. Barcodes improve segmentation accuracy. Across all validated metrics, the accuracy of the segmentations improved when using enhanced barcodes as input compared to raw GFP.

Here each “method” refers to the respective image input to the affinities network. For each method, we evaluated standard neuron segmentation metrics against manually annotated skeletons. These metrics included Variation of Information (VoI and normalized VoI, NVoI; (Meilă, 2007); lower scores are better), Expected Run Length (ERL and normalized ERL, NERL; (Januszewski et al., 2018); higher scores are better), and the Min-Cut Metric (MCM; (Sheridan et al., 2022); lower scores are better). Here we focus on NVoI results. For detailed metric evaluations, see Supplementary Section 6.2.6.2, Supplementary Figure 9, Supplementary Figure 10, and Supplementary Table 18.

We found providing all raw barcode color channels as inputs to this network significantly improved segmentation quality compared to providing only the GFP channel (R-S vs R-GFP; NVoI Sum 0.251 vs 0.336, Supplementary Figure 9, Supplementary Figure 10, Supplementary Table 18). We tested this with two approaches. The first approach simply averaged the barcode intensities across the channel dimension such that the affinities network took a single averaged channel as input. In the second approach we gave the affinities network access to all barcode channels (Supplementary Figure 9.A). Surprisingly, we found that simply averaging the intensities across the channels showed similar performance (R-M vs R-S; NVoI Sum 0.246 vs 0.251, Supplementary Figure 9, Supplementary Figure 10, Supplementary Table 18).

We hypothesized this was due to discontinuities and noise in the barcode signals. Therefore, in order to maximize the usability of the barcode information, we performed a supervised signal enhancement step with the aim of preserving the barcode intensity within the neurons. Using sparsely annotated neurons as ground-truth, we taught a UNet to predict the average protein bit intensities within a neuron using the raw data as input to create an enhanced signal for all protein bit channels (Figure 3.A, network 1, Supplementary Figure 5, Supplementary Section 6.2.2). Providing these enhanced color channels as input to the network significantly improved performance compared to both the previous multi-color (E-S vs R-S; NVoI Sum 0.176 vs 0.251) and averaged channel networks (E-M vs R-M; NVoI Sum 0.192 vs 0.246) (Supplementary Figure 9, Supplementary Figure 10, Supplementary Table 18).

To further extend the diversity of the barcode space, we set out to leverage information from the variability in relative protein bit intensities between cells. We used a simple multi-layer perceptron (MLP) (Figure 3.A, network 3) to project the enhanced barcodes onto a higher dimensional unit sphere to learn a uniform embedding space (Wang & Isola, 2020). The aim of this approach is to preserve barcode uniformity within neurons, and increase separability across different neurons. Specifically, we minimized a contrastive loss with two terms, one to pull embeddings within an object towards that object’s mean, and the other to push the mean embedding of different objects apart.

With this learned embedding, nearest cosine distances between the intensities of distinct neurites were more widely distributed (spanning [0,2]) than in the raw data, demonstrating increased separability (Figure 3.B, Supplementary Figure 6). However, the embedding did not preserve uniformity within neurites, preventing direct clustering into unique segments. Since the uniform embedding enforces strong boundaries between cells, we were able to efficiently generate “barcode” affinities by simply computing affinity weights via cosine similarity (Figure 3.A, barcode affinities). We combined these barcode affinities with the predicted affinities from the Local Shape Descriptor (LSD) network (“shape” affinities; Figure 3.A, combined affinities) thereby making effective use of both barcode and morphological information.

By combining these affinities, we observed a further increase in accuracy and specifically were able to decrease the false merge rates in the segmentation. Compared to the GFP baseline model, we observed a fourfold increase in NERL (E-S+U vs R-GFP; 0.18 vs 0.046, Figure 3.D, Supplementary Figure 9, Supplementary Figure 10, Supplementary Table 18), and a twofold decrease in the NVoI Sum (E-S+U vs R-GFP; 0.163 vs 0.336, Figure 3.D, Supplementary Figure 9, Supplementary Figure 10, Supplementary Table 18) and edits needed (E-S+U vs R-GFP; 690 vs 1466, Figure 3.D). Taken together, we show that combining shape and barcode information into an affinity graph partitioning approach increases segmentation accuracy of neuronal morphologies at minor computational costs (Supplementary Figure 11).

### 2.4 Barcode information can be used for automated proofreading

We next asked whether, in addition to directly assisting the segmentation, the barcodes could be used for automated proofreading i.e., using the unique barcode identities to automatically correct errors that remain after segmentation. Proofreading errors can be separated into two types: merge and split errors. We reasoned that split errors would be easier to correct with the barcodes than merge errors, as merge errors directly affect the barcode readout of the resulting segments (Supplementary Section 6.2.5). We therefore aimed to limit the number of false merges in the initial segmentation, via the combination (product) of the shape and barcodes affinities (Supplementary Section 6.2.5). With fewer merges we were able to then re-merge these disconnected segments using the barcodes.

To efficiently re-merge false splits in the segmentation, we first skeletonized (Sato et al., 2000; Silversmith et al., 2021) all segments, and created a spatial-partitioning KDTree on end node positions of the skeletons. We then created an “average barcode” for each segment by combining the underlying raw barcode intensities masked to the supervoxels. Finally, we defined edges between pairs of nodes with weights equal to the cosine distance between their respective “average barcode”.

Considering each pairwise edge would be too expensive, so for each candidate node we instead considered all pairs of skeletons within a certain distance threshold for merging and splitting and added a splitting signal for skeletons above that threshold. For practical reasons we also have a separate distance threshold for the splitting signal to limit the number of pairwise candidates but this, in theory, could be pushed infinitely (Supplementary Section 6.2.5). Similar to regular affinity processing, including long range split biasing edges here helps mitigate the cascading effects of small merge errors.

By formulating the task as such, we were able to cluster the edges with mutex watershed (Wolf et al., 2018), similar to the initial affinity segmentation. The resulting lookup table was then used to relabel the original segments (Figure 4.A). While sensitive to local barcode collisions, this optimization limits the occurrence of spurious merges based on similarities in barcode space and allows for easy tuning of the merge rate.

**Figure 4.**
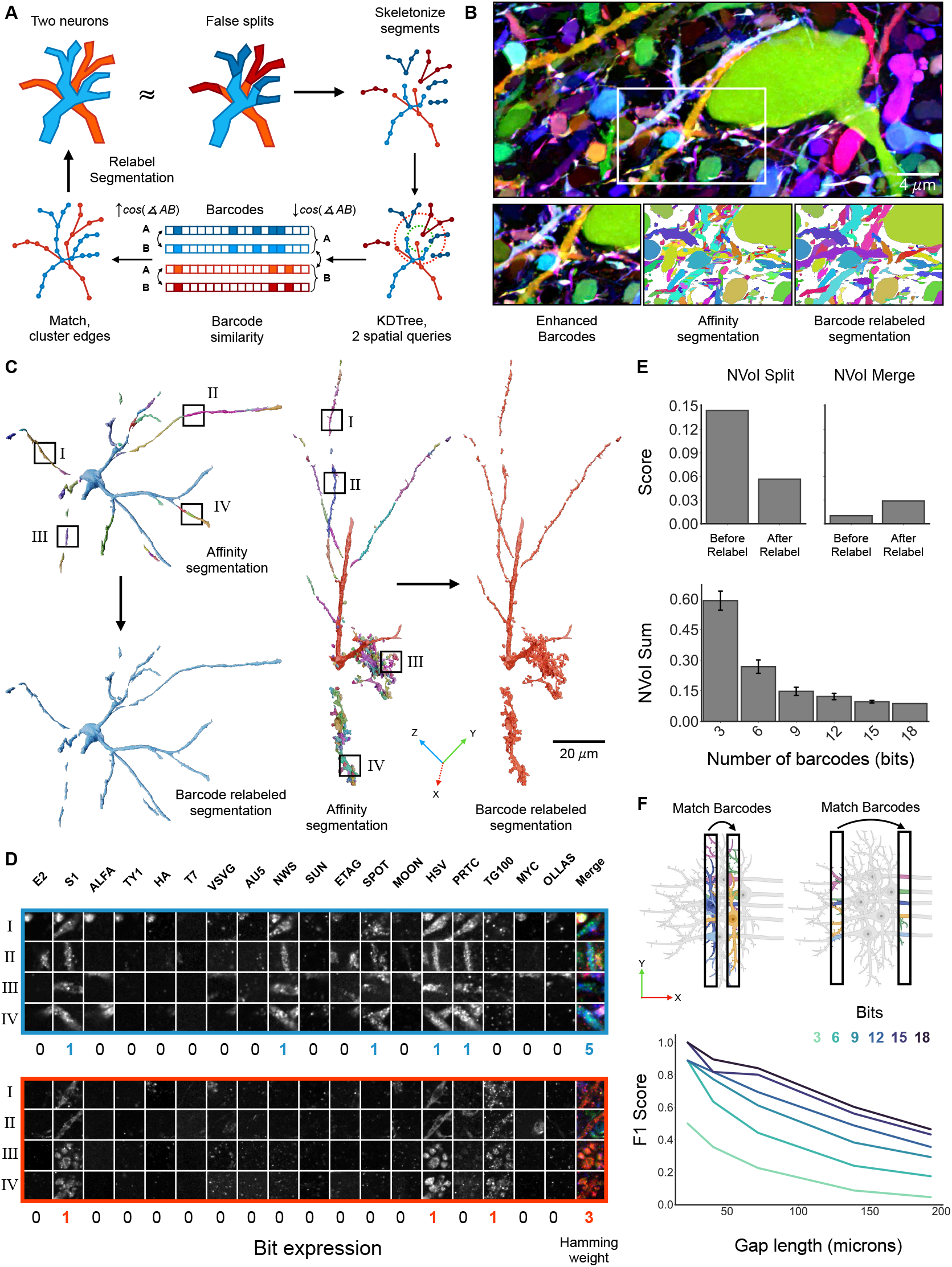
: Using barcodes for automated proofreading. A. Barcode relabeling overview. Given a split-preferring affinity Figure 4: (*continued*) graph segmentation, we skeletonize the segments and create match candidates using a KDTree with two spatial queries (∼ 10 µm, ∼ 30 µm). Using the distance between the barcodes as edge weights, we perform a matching and use the resulting clustering to relabel the initial segmentation. B. Example images. A maximum intensity projection across 40 sections of enhanced barcode data, visualized with a multi-channel shader. Insets show the initial affinity segmentation and the resulting barcode-relabeled segmentation. C. Example Meshes. Two example neurons with their resulting meshes from the affinity segmentation and corresponding barcode-relabeled meshes (≥ 50% volumetric overlap). Boxes correspond to panels below. D. Barcode expression. Corresponding raw data for boxes on above neurons for each of the 18 channels. Expression is consistent across each neuron and different between neurons, allowing for gap crossing. E. Barcode relabeling improves segmentation accuracy. By relabeling segmentations with the barcodes we are able to decrease the split rate while not drastically increasing the merge rate. The accuracy is increased with the number of barcodes used for matching. F. Crossing larger gaps. While barcodes can be used to match segments across gaps, the chance of barcode collisions preventing successful matching increases with gap size and decreases as more barcodes are used.

We found that by relabeling segments using barcodes, we were able to bridge small gaps in the tissue due to signal loss and masking (Figure 4.B,C). This is possible since the barcode expression level across channels is relatively consistent across neurons but differs between neurons (Figure 2.F). This approach is still sensitive to collisions in barcode space (Supplementary Figure 14), but these collisions are mostly absent inside small local windows. Increasing the spatial context for matching will lead to more merges (Supplementary Section 6.2.5), but opens the door for integration with globally aware approaches (Januszewski et al., 2025; Troidl et al., 2025).

Nevertheless, on an evaluated sub-ROI, we found barcode matching significantly improved segmentation accuracy (NVoI Sum 0.154 vs 0.086, NERL 0.446 vs 0.610) specifically by accurately re-merging previously split segments (NVoI Split 0.144 vs 0.057), while limiting the amount of increased false merges (NVoI Merge 0.010 vs 0.029), (Figure 4.E, Supplementary Section 6.2.5, Supplementary Section 6.2.6.3,). Importantly, by sub-selecting the number of considered channels, we found that the accuracy of automated proofreading increased with the number of barcodes (Figure 4.E). Since this automated proofreading approach is computed locally on skeletons it could be extended to run block-wise on larger volumes. Finally, by simulating larger spatial gaps we found that barcode information could potentially be used to merge segments across larger distances (Figure 4.F, Supplementary Section 6.2.6.4, Supplementary Figure 15, Supplementary Table 20, Supplementary Table 21), a promising direction for future research to reinforce the strength of barcodes in overcoming sample fragility.

### 2.5 PRISM enables synaptic analysis with molecular annotations

Previous studies have shown that resolving synapses and assigning them to neurons is possible using expansion microscopy (Gao et al., 2019; Shen et al., 2020). We therefore hypothesized that the unique capabilities of PRISM would allow us to not only reconstruct neuronal morphology but also map synaptic connections by characterizing their molecular machinery. Specifically, we observed that with 5x expansion, the excitatory postsynaptic marker PSD95 and the presynaptic protein Bassoon formed spatially well defined and partially overlapping pairs (centroid distance: 99±53 nm; n: 6784 pairs), indicating the location of putative synapses (Figure 1.E,F, Figure 5.A). This allowed us to combine a ML segmentation pipeline with an overlap-based heuristic algorithm to reliably segment each marker in the volume (F1 scores Bassoon: 0.97, PSD95: 0.95) and group them into individual synapses (F1=0.95; Figure 5.B). In this way we identified a total of ∼ 9 million synapses in the imaged volume with an average density of 1.02 ± 0.62 excitatory synapses per µm^3^, which aligns with previous estimates from EM (Santuy et al., 2020). These detected synapses could then be assigned to segmented neurites with high accuracy based on their spatial overlap (F1=0.85; compared to human annotators; Section 4.5). To evaluate this approach, we examined the well-studied synaptic connections on the dendrites of CA2/3 pyramidal neurons using a set of proofread neurons that could be traced back to their somas (n = 6; Figure 5.C-E) and identified a total of 12,168 synapses (average: 2,028 ± 1,092 synapses per cell). Looking at the distribution of excitatory synapses on both apical and basal dendrites, we observed the number of synapses on the dendritic tree increased with distance from the soma (fold change at 85 µm, apical = 2.54, basal = 2.04, Figure 5.D,E). This reflects the well known layered organization of the hippocampus, where apical and basal dendrites originating in the pyramidal layer extend into the stratum radiatum and oriens respectively, and form connections with layer-segregated excitatory afferents (Megías et al., 2001). These findings show that PRISM can be used to investigate circuit level synaptic innervation patterns.

**Figure 5.**
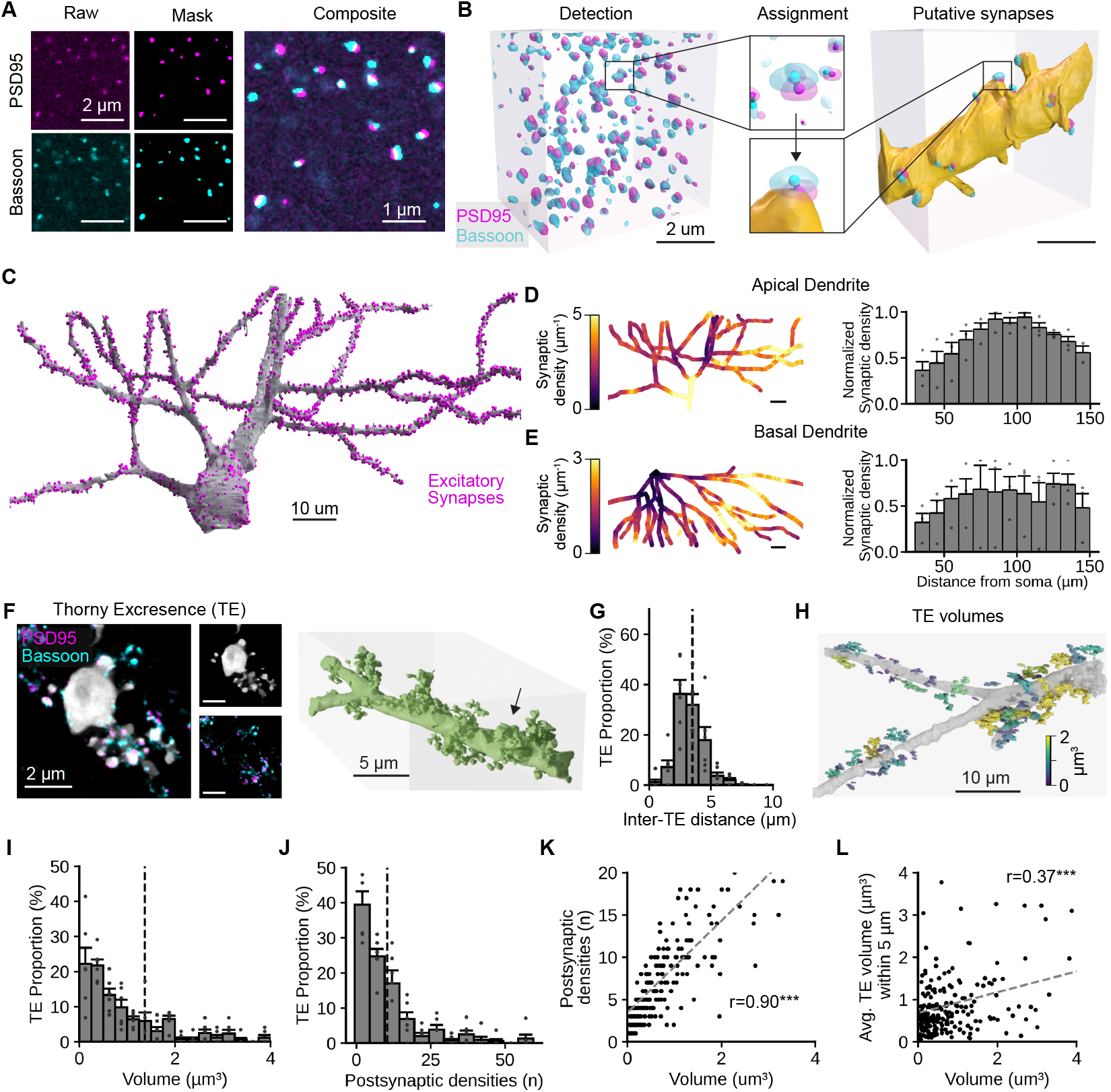
: Analysis of synaptic connections and their distribution. A. Example synaptic marker detection of PSD95 (top) and Bassoon (bottom). Left, raw intensity images of each antibody label. Center, predicted label masks. Right, composite showing overlap between pre- and postsynaptic markers used to determine true synapses. B. Schematic of ML synapse detection and assignment. Left, 3d volume of detected overlapping PSD95 (magenta) and Bassoon (cyan) labels. The center of each detected label is shown as a solid sphere. Square highlights the zoomed-in region. Center, example assignment of a pair of PSD95 and Bassoon labels to segmented neuronal morphology. Right, same area shown on the left but only showing label markers that could be assigned to the shown dendrite. C. Example of proofread pyramidal neuron with assigned excitatory synapses (magenta). D. Distribution of excitatory synapses on apical dendrites. Left, heatmap of excitatory synapses per µm on a reconstructed dendrite. Right, density of excitatory synapses normalized to the maximum value per cell as a function of soma distance showing that excitatory synaptic density increases with distance. E. Distribution of excitatory synapses on basal dendrites. Same representation as in D but for basal dendrites. F. Example of a thorny excrescence on a CA3 apical dendrite. Composite image showing single protein bit channel (gray) with PSD95 (magenta) and Bassoon (cyan) antibody label. Right, 3d reconstruction of the same stretch of dendrite. The arrow highlights the location of the thorny excrescence (TE) shown on the left. G. Histogram showing the distribution of distance between TEs and their nearest neighbor on the dendrite. The dashed line shows the average value. H. Mapped TEs of a single dendrite with their volume encoded by color. I. Histogram of detected TE volumes. The dashed line shows the average value. Individual points mark the values of individual cells. J. Similar to I but shows the histogram of the number of postsynaptic densities per TE. K. Correlation between TE volume and number of detected postsynaptic densities. The gray dashed line indicates the best-fit linear regression and its statistical significance. L. Similar to K but showing the correlation between a TE’s volume and the volume of those within 5 µm, indicating that the volumes of neighboring TEs on the same dendrite are significantly correlated. r=Pearson correlation coefficient, ^***^: p*<*0.001.

Due to the optical resolution of PRISM we were also able to investigate the fine-detail nanostructural organization of synaptic connections. Specifically, we observed the elaborate synaptic complexes known as thorny excrescences (TEs; Figure 5.F), formed by CA3 pyramidal neurons at contacts with mossy fiber axons from granule cells in the dentate gyrus. Since TEs contain multiple large synaptic active zones they can greatly affect synaptic integration and dendritic membrane potential. However, due to their complex morphology, conventional light microscopy approaches have struggled to characterize them (McAuliffe et al., 2011) and they are most often studied using EM approaches, which lack molecular information (Zheng et al., 2025). We therefore used the molecular information in our PSD95 and Bassoon stains to investigate a total of 247 proofread TEs across 6 dendrites in CA3. We observed multiple TEs were often grouped close together (inter-TE distance: 3.5 ± 1.3 µm; Figure 5.G) on shorter sections of the dendritic tree, matching previous descriptions of their cluster-like distribution in ∼ 10 µm stretches (Figure 5.H; Gonzales et al., 2001; McAuliffe et al., 2011). TE volumes were highly diverse (Figure 5.I) with an average volume of 1.37 ± 0.51 µm^3^. Similarly, the number of postsynaptic densities present on each TE also strongly varied (average: 10.38 ± 2.42 synapses per TE; Figure 5.J) and was strongly correlated to volume size (r: 0.90; p*<*0.001; Figure 5.K). Due to the observed clustered distribution of TEs (Gonzales et al., 2001) we investigated whether nearby TEs had similar properties compared to more distant TEs. Interestingly, we discovered a strong correlation between the volume of each TE and the average volume of nearby TEs (*<*5 µm) on the same dendrite (r: 0.37; p*<*0.001; n = 228 TEs across 6 cells; Figure 5.L). One model that could account for this observation is that clusters of TEs perform similar sampling of the surrounding mossy-fiber input pool, an effect potentially related to their relative position in the mossy fiber pathway (Zheng et al., 2025).

Taken together, these examples illustrate how PRISM can be applied to span studies of circuit-level innervation patterns and nanoscale synaptic architecture that standard light microscopy cannot resolve. Its iterative multiplexing profiles the molecular machinery of these contacts and could further be expanded with cell-type markers of interest for targeted circuit mapping.

## 3 Discussion

Recent advances in microscopy and sample preparation now enable high-resolution brain imaging across large spatial volumes, a critical step toward understanding the cellular basis of brain function and informing therapeutic development. However, these advances exceed our ability to reconstruct neuronal morphologies at scale. Scalable neuron reconstruction faces two primary challenges: first, the limitations in the accuracy of automatic neuron tracing (“Scaling up connectomics — Reports,” 2023); and second, signal discontinuities (e.g. staining artifacts or serial section loss) which impair or prevent continuous neuron tracing. Incorporating molecular annotations into morphological reconstructions is also challenging, yet essential for understanding the complex interactions between drug targets and the cells and circuitry which they target. To overcome these challenges, we developed PRISM, a method for robust, scalable, and molecularly-annotated neuron reconstruction, designed to allow individual labs to incorporate morphological analysis into everyday neuroscience.

PRISM integrates complementary advances, including protein barcode labeling with diverse antigens; expansion microscopy with extensive multiplexing capacity; barcode-augmented segmentation and proofreading; and molecular-annotation of reconstructed neurons. We demonstrate a system capable of delivering more than 100,000 unique protein codes (theoretical maximum of 2^18^ − 1 = 262,143). Being protein based, these barcodes ensure morphological filling in contrast to sparse RNA-based approaches (X. Chen et al., 2019; Goodwin et al., 2022; Kebschull et al., 2016; Yuan et al., 2024). This represents a 750-fold improvement in the number of labels over previous multispectral methods (Leiwe et al., 2024; Livet et al., 2007). Essential to PRISM is a co-optimized method for iterative antibody staining and destaining in expansion microscopy. By refining conditions along multiple dimensions of gel chemistry and imaging set-up, we achieved excellent preservation of both barcode antigens and endogenous proteins. We applied PRISM to generate a dataset in mouse CA3 hippocampus covering over 10 million cubic microns and 23 molecular targets, providing a uniquely large and highly-multiplexed resource.

Using this dataset, we demonstrated that protein-based neuronal barcoding has the potential to address the cost and robustness challenges of scalable morphology reconstruction. Incorporating barcode information in automatic segmentation and proofreading rather than relying on shape information alone (i.e., eGFP), improved multiple measures of traceability, most prominently neurite run length (∼ 8x increase). Crucially, we showed automatic proofreading can bridge spatial gaps to reconnect neurite segments both locally and even across hundreds of microns, a major step towards addressing signal discontinuity challenges. Finally, we demonstrated the power of combining molecular annotations with morphological reconstructions by revealing a size correlation between TEs on the same dendrite.

PRISM could be applied in a range of existing neuroscience work. This platform is ready for use, supporting applications such as improved accuracy and throughput in brain-wide projection mapping (Winnubst et al., 2019); barcoding of viral transgenes (e.g. enhancers Hunker et al., 2025); lineage tracing (Li et al., 2021; Loulier et al., 2014); barcoded screening and analysis of gene and cell therapies (Borch Jensen & Marblestone, 2021); and multiplexed synapse profiling (O’Rourke et al., 2012). Future developments could readily expand the capabilities of PRISM with a minimum of risk, building from the firm foundation here set. Integrating PRISM with ultrastructure labeling and high expansion factors could extend self-correcting neuron tracing to dense connectomics. As barcoding capacity scales exponentially with detectable antigens, PRISM could scale to millions or even billions of unique combinations with the addition of a few new antigens and antibodies. For example, expanding the antigen-antibody palette from 18 to 30 – newly feasible in the era of generative AI protein design (Cao et al., 2022; Vosbein et al., 2024) – could enable PRISM to uniquely barcode every cell in the mouse brain. To image such large volumes, enhanced labels (Viswanathan et al., 2015) and molecular signal amplification (Saka et al., 2019; Schwarzkopf et al., 2021) could allow modern scientific CMOS cameras to be run at maximum speed. To fully segment such volumes, self-correcting barcodes could be further augmented by advances in global shape-based segmentation and proofreading (Januszewski et al., 2025; Troidl et al., 2025). In conclusion, PRISM provides a scalable foundation for next-generation morphological reconstruction techniques, addressing the disparity between the rapid generation of imaging data and the slower pace of accurate reconstruction and annotation.

## 4 Method

### 4.1 Barcodes

#### 4.1.1 Barcode vector design, cloning, and production

Barcode vectors were assembled using pAAV-CAG-GFP as a backbone vector. pAAV-CAG-GFP was a gift from Edward Boyden (Addgene plasmid #37825; http://n2t.net/addgene: 37825 RRID:Addgene 37825). Briefly, the backbone vector was digested with BsrGI-HF and EcoRI-HF. Gblocks with complementary overhangs were designed for Gibson assembly and ordered from Integrated DNA Technologies. Assemblies were transformed into NEB Stable cells and, after Sanger confirmation in individual colonies of the correct insertion, DNA was individually maxiprepped. The final assembly and ITR stability was verified with whole-plasmid sequencing performed by Plasmidsaurus. All plasmids used are deposited with Addgene (Addgene #242764-242781).

AAV transfer vectors were pooled at equimass ratios and packaged as a pool by Sanford-Burnham’s Viral Vector Core. Virus was purified via iodixanol gradient ultracentrifugation and quantified with qPCR of ITRs and measured at 9.1E13 vg/mL.

#### 4.1.2 Barcode delivery

All animal work and injections were performed by the Crick Biological Research Facility and Surgical Services. 51-52 week old C57BL/6Jax mice were subcutaneously injected with meloxicam (10 mg/kg) and buprenorphine (0.1 mg/kg), then unilaterally injected with 250 nl 9.1E13 vg/mL virus (A/P: −1.70 mm, ML: ±100 mm, DV: −1.5 mm, Paxinos). At least 3 weeks after injections, mice were perfused with ice-cold 1X PBS for 1 minute and then ice-cold 4% PFA in 1X PBS for 10 minutes under terminal anaesthesia (intraperitoneal injection of 600 mg/kg pentobarbital). Brains were harvested for downstream sample preparation (Section 4.2).

### 4.2 Sample prep and imaging

Harvested brains were post-fixed in 4% (w/v) PFA in 1X PBS for 28 hours at 4°C, then quenched in 0.1 M glycine in 1M Tris, pH 7.5 for 20 hours at 4C, before sucrose impregnation in gradient steps of 15% (w/v) sucrose in 1X PBS, 0.05% (w/v) NaN3, overnight at 4°C followed by 30% (w/v) sucrose in 1X PBS, 0.05% NaN3, for 2 days at 4C. Brains were equilibrated in a 1:1 mixture of OCT and 30% (w/v) sucrose for 4-6 hours at 4°C, incubated in 100% (v/v) OCT for 2 hours at 4°C, then incubated in fresh 100% (v/v) OCT overnight at 4°C. Brains were mounted in 100% OCT in cryomolds and frozen on dry ice, before cryostat sectioning 50 µm coronal sections on a Leica CM1860 UV cryostat at −20°C.

Due to inconsistencies in sodium acrylate quality across commercial vendors, sodium acrylate for MAGNIFY gels were synthesised by the Crick Chemical Biology STP by dissolving acrylic acid in methanol and sodium hydroxide in methanol in a 1:1 molar ratio in an ice-bath. Upon precipitation of sodium acrylate, the slurry was filtered through a Bü chner funnel and then vacuum desiccated for at least 2-3 days to yield sodium acrylate powder. Batches were quality controlled via NMR and LC-MS/MS analysis.

50 µm coronal sections positive for GFP barcode expression were preincubated in MAGNIFY monomer solution (34% (w/v) sodium acrylate, 10(w/v) acrylamide, 4% (v/v) DMAA, 1% (w/v) NaCl, 0.01% (w/v) bis-acrylamide, in 1X PBS) with 0.001% (w/v) 4-hydroxy-TEMPO, 0.2% (w/v) TEMED, 0.2% (w/v) APS, on ice shaking in a 12-well plastic plate, before gelation at 37C overnight in glass slide chambers. Chambers consisted of untreated glass slide, where brains were mounted, one #0 coverslip spacer on each side, and a hydrophobic glass slide as the cap. After gelation, GFP-positive regions of the sample were trimmed and homogenized with pre-heated MAGNIFY homogenisation solution (10% (w/v) SDS, 8M Urea, 25 mM EDTA, 2X PBS) at 80°C in an oven for 7 hours, and then washed with 1X PBS, 0.05% NaN3 for 20 min × 3 at room temperature (RT), shaking, then 1% (v/v) DGME in 1X PBS for 1 hour at 60°C in an oven, then washed with 1X PBS, 0.05% (w/v) NaN3 for 20 min × 3, at RT. Samples were then expanded in ddH2O for at least 10 min × 3.

Expanded MAGNIFY gels were trimmed and then preincubated in re-embedding solution (4% (w/v) acrylamide, 0.15% (w/v) bis-acrylamide in 5 mM Tris with 0.1% (w/v) TEMED and 0.1%(w/v) APS) on ice, shaking, in a 6-well plate, before gelation overnight at 37°C. Gelation chambers consisted of silanized glass-bottom 6-well plates (Cellvis) with each spacer as 3x #2 glass coverslips and tops as 25-30 mm Sigmacote-treated round #1 coverslips. Plates were silanized by incubating 1 mL per well of bind silane solution (1%(v/v) 3-(Trimethoxysilyl)propyl methacrylate (Sigma-Aldrich M6514), 80% (v/v) ethanol, 2% (v/v) acetic acid) for 1-3 minutes, followed by 3 × 30 sec washes with 1 mL of 100% ethanol per well and air drying. Gels were trimmed to the MAGNIFY sample containing region, washed with 1X PBS, and then processed for iterative immunostaining.

In each round of staining, imaging, and stripping, reembedded MAGNIFY samples were first blocked with maleimide in 1X PBS for 30 minutes at RT in order to block photo-crosslinking via oxidation of free sulfhydryl groups during imaging (Gut et al., 2018). Samples were then washed with 1X PBS for 10 min × 3, before blocking with blocking buffer (1% (w/v) BSA in 0.2% (v/v) Triton X-100 in 1X PBS), 0.05% NaN3 for 1 hour at RT. Samples were then stained with primary antibodies in blocking buffer overnight at RT, shaking, then washed with blocking buffer, 20 min × 3, RT, before incubation with secondary antibodies in blocking buffer overnight at RT, shaking. Secondary antibodies were washed off with blocking buffer, 20 min × 3, RT. Samples were then mounted in glass-bottom 6-well plate (Cellvis) with 2-stacked Bio-Rad gaskets as spacers and a 25mm round coverslip and M15 washer on top. To prevent photobleaching and photo-crosslinking, the imaging chamber was filled to the brim with glucose oxidase-catalase + Trolox buffer (1.4 mg/mL ≈≥140 U/mL glucose oxidase (Sigma-Aldrich G2133), 4 µl/mL ≈≥1600 U/mL catalase (Sigma-Aldrich C100) in 100 mM Tris, 25 mM NaCl, 10% (w/v) glucose, 2 mM Trolox (Sigma-Aldrich 238813), pH 8.0) and sealed with a PCR film to maintain anoxic conditions. Samples were imaged with a Nikon-CSU Ti-2 W1 system with a Nikon Apo LWD 40x WI *λ*S DIC N2 (NA = 1.15) water immersion lens. After imaging, samples were stripped with pre-heated MAGNIFY homogenisation solution at 80°C for 5-6 hours, and then washed with 1% DGME at 60°C in an oven, shaking, for 1 hour, then washed with 1X PBS 20 min × 3 at RT, shaking, before the next round of immunostaining.

### 4.3 Stitching and Registration

The reported image volume was captured by a total of 62 partially-overlapping (15% in each dimension) microscopy image stacks each ∼ 345×7345×380 µm in size. These volumes needed to be computationally stitched together in order to create a continuous volume. To achieve this, we made use of the bigstitcher (Hö rl et al., 2019), and accompanying bigstitcher-spark, software packages. In short, bright interest points were identified in each volume using a difference-of-gaussian spot detector approach (sigma=1.6). Fiducial descriptors were then generated based on their spatial configuration with their 3 nearest neighbors and matched with fiducials in overlapping volumes. True matches were identified using a translation-invariant, model-constrained Random Sample Consensus (RANSAC) algorithm (Preibisch et al., 2009). Identified pairwise links between all volumes were then used to calculate a globally optimized affine for each individual volume.

In order to register the imaged volumes from iterative cycles of immunostaining together into a continuous volume, we imaged the abundant ALFA protein bit in every consecutive round to act as a registration channel. ALFA was chosen as a reliable channel validated across many rounds of immunostaining and stripping. For alignment, manual registration to the first imaging round was performed on downsampled images (800×672×672 nm voxel size) using the BigWarp software package in Fiji by selecting matching fiducials in the registration channel of both rounds (Bogovic et al., 2016; Schindelin et al., 2012). The resulting landmarks were then used to compute a displacement field with a thin plate spline function and applied to all high resolution channels using an adapted version of the BigStream rendering algorithm^*^. Staging and execution of all related stitching and registration functions was performed using custom nextflow pipelines^†^ (Di Tommaso et al., 2017).

### 4.4 Segmentation

Following stitching and registration of barcodes across imaging rounds, we developed a neuron segmentation pipeline that incorporates the barcode color information. Assuming a perfect barcoding space, the barcodes could theoretically be directly clustered into segments. However, in practice, this is challenging due to several factors, including variability across datasets in the number of available barcodes, noise, artifacts, and collisions in barcode space. We therefore instead opted to incorporate the barcodes directly into an affinity graph agglomeration pipeline, as this is robust to variations in data, is scalable, and trivially parallelizable across chunks. Our segmentation pipeline processes the registered multi-channel barcode volume through a series of four specialized neural networks. Each network is trained for a specific objective, and their outputs are combined to produce the final neuron segmentation.

#### 4.4.1 Training objectives

##### 4.4.1.1 Barcode Signal Enhancement

###### Overview

Due to the noise and variable intensity ranges across the volume, we first perform a supervised signal enhancement and denoising. Traditional denoising methods can be useful for removing noise prior to segmenting a volume (Weigert et al., 2018). However, since we don’t have access to a clean target volume, we can’t utilize supervised denoising approaches such as CARE (Weigert et al., 2018). Furthermore, we found that unsupervised denoising approaches, such as Noise2Void (Krull et al., 2019), were insufficient for removing specific noise present in the data. Since the main goal was to diffuse the barcode signal throughout the objects, unsupervised denoising is not sufficient as the model also needs to learn to fill in gaps inside objects which is not achievable without target ground-truth. Instead, using ground-truth labeled cells, we derived ground-truth barcodes by computing the average barcode intensities (across each channel) masked to each label. We then trained a U-Net (Ronneberger et al., 2015) to learn the average barcode from the raw intensities, in a channel-agnostic fashion. While the main goal was to diffuse barcode intensities throughout the volume, it also resulted in the removal of noise as a side effect.

###### Formal Definition

Let *x*_*C*_: Ω ↦ ℝ^C^;and *x*_*c*_: Ω ↦ ℝ; | *c* ∈ *C* where Ω ⊂ ℕ ^3^represents the voxel spatial locations in a barcoded 3D plus channels (*C*) image volume. Let *f* (*x*_*c*_): Ω ↦ ℝ; be the enhancement network learning the *x*_*c*_ function. Given a sparse ground-truth instance segmentation, let *L* = {0, 1, …, *l* }where 0 denotes background or unlabeled regions and *>* 0 denotes unique foreground objects and *y*: Ω ↦ *L*.

We define a binary weight mask *w*(*v*):

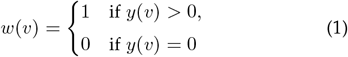

For each label *k* ∈ {1, …, *l* } we define a set of voxels belonging to that label:

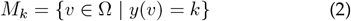

And compute the average intensity *µ*_*c*_(*k*) ∈ ℝ; over all voxels in *M*_*k*_:

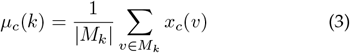

We then define the target function *d*_*c*_(*v*) as the difference between the average *µ*_*c*_(*y*(*v*)) and input *x*_*c*_(*v*) image (i.e the residual barcode):

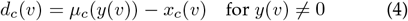

Where the average can then be recovered (i.e., during inference):

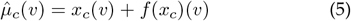

Our primary learning objective is to then densely infer a residual barcode from the raw barcode data for each labeled object in the volume:

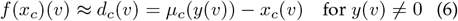

We then minimize a MSE loss in mini-batches, weighted by the labeled regions:

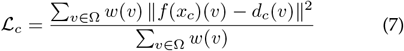

Every operation *x*_*c*_, *f* (*x*_*c*_), *µ*_*c*_, *d*_*c*_, ℒ_*c*_ can be trivially vectorized into *x*_*C*_, *f* (*x*_*C*_), *µ*_*C*_, *d*_*C*_, ℒ_*C*_ to operate on the full set of channels *C*, including the network function *f*. We do this in PyTorch^‡^ by simply treating channels as a batch dimension in order to work with arbitrary numbers of barcodes.

##### 4.4.1.2 Affinities and Local Shape Descriptors (LSDs)

###### Overview

We next extracted a boundary representation (affinity graph) from the enhanced barcodes. We learned long range affinities (Lee et al., 2017) in addition to nearest neighbor affinities as an auxiliary learning task to force the network to use more context in its receptive field, and to use as attractive/repulsive weights during downstream processing. Additionally, we jointly learned Local Shape Descriptors (LSDs) (Sheridan et al., 2022), to further improve the added context for the network. This is learned sparsely through a multi-channel 3D U-NET using a weighted combined MSELoss minimizing the sum of the affinities and LSDs losses (Sheridan et al., 2022).

###### Formal Definition

Following inference of the enhancement network, using the enhanced barcodes *x*: Ω ↦ ℝ; ^*C*^we then train (sparsely) a multitask (MTLSD) network to learn an affinity graph with Local Shape Descriptors as an auxiliary learning task (Sheridan et al., 2022).

##### 4.4.1.3 Contrastive Uniform Embedding

###### Overview

We found a subset of merge errors on the learned affinities occurred between neurons with distinct barcodes due to a smooth change in intensities. We therefore aimed to represent the distance between barcodes directly as affinities to enforce the constraint that different barcodes do not get merged. However, since the raw barcode channels do not have consistent intensity ranges, we used a simple MLP to first project the barcodes into a higher-dimensional space where distances are more reliably computed. Given a barcode of length *n* we project it into a *d*-dimensional space and enforce unit length while maximizing inter-barcode distance. This pushes voxels inside a neuron closer in embedding space while voxels across neurons are pushed farther away. Since this is sensitive to noise and variance in the image data, we train with the enhanced barcodes as input rather than the raw barcodes. Inspired by other contrastive learning approaches (Lee et al., 2021; Wang & Isola, 2020), we compute cluster means and formulate the learning objective as:

1. Minimize distances of samples to their cluster mean (uniform objective).
2. Maximize distances between cluster means (align objective).

This approach is similar to learning dense voxel embeddings via deep metric learning (Lee et al., 2021), with three important distinctions:

1. Our regularizing term encourages unit length.
2. We encourage the neuron embeddings to be uniformly distributed on the unit hypersphere, rather than using a margin term.
3. Rather than learning this as a shape embedding (i.e via a U-Net with spatial context), we simply use an MLP to learn this as a color embedding (since this information is directly encoded in the data).

###### Formal Definition

We first define our learned embedding function *f* (*x*_*C*_): Ω ↦ ℝ; ^D^where *C* is our set of barcode channels, and *D* is our set of embedding channels. To define the learning objective, we need to compute the mean barcode in our learned embedding space given some cluster *M*_*k*_:

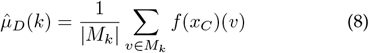

and normalize to unit length:

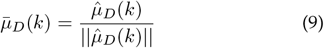

For the first objective, to maximize the distance between cluster means, we use the empirical approximation of the logarithm of the expected pairwise Gaussian potential (Wang & Isola, 2020):

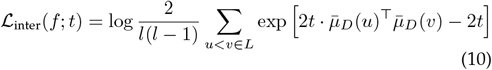

To minimize the variance within clusters we directly minimize the angle between the cluster and the mean:

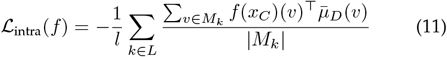

The final loss is then the sum of both losses with *α, t* as tunable hyper-parameters:

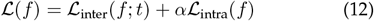

##### 4.4.1.4 Barcode Expression Probabilities

###### Overview

Due to the sparsity of the barcodes, a large portion of the image is background and should therefore be masked out during post-processing. We found that training traditional 3D binary foreground/background and semantic networks did not produce an optimal mask of “barcode-able” regions, and therefore introduced significant errors during segmentation. This was likely due to the large variance in image intensities across the volume, and across channels. Instead, we found a channel-agnostic approach to this task yielded superior results. A traditional foreground / background network would be trained using a single image with a single mask training signal. For *k* channels, this would require *k* highly redundant and expensive-to-generate training masks. Instead, we simply used a single channel mask of all labeled objects, and performed a max projection across channels directly in the loss.

###### Formal Definition

Our learning objective is to densely infer the probability of a barcode expressing in a given channel from the raw barcode data, for each labeled object in the volume. For this task, we reuse *x*_*c*_ and *x*_*C*_, and define a new mask:

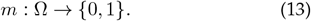

The channel-agnostic binary network that we train now operates in a map reduce framework, by taking the max across channels of the predicted probabiltiies:

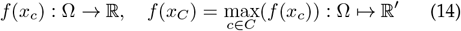

Due to the lack of differentiating signals between certain objects such as dim cell bodies and bright background, we also introduce a boundary-weighted mask where we only consider the edges of objects within some distance *τ* for training. This mask is computed on the thresholded euclidean distance transform (EDT) of the labels:

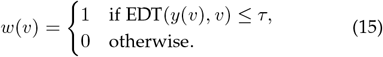

We then optimize a weighted binary cross-entropy (BCE) loss:

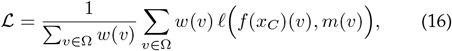

#### 4.4.2 Network Architectures

The affinities and LSDs were implemented in a multitask U-Net architecture (Sheridan et al., 2022). Both the enhancement and expression networks both used a channel-agnostic U-Net in which the channel dimension was simply treated as a batch dimension, in order to process datasets with differing numbers of barcodes. The uniform embedding network was a simple MLP^§^. Networks were trained with Gunpowder^¶^ and Pytorch^‡^. Training scripts are available in a public repository^||^. For more details on training, see Supplementary Section 6.2.2.

#### 4.4.3 Post Processing

Prediction and post-processing were done in a block-wise fashion. Block distribution was handled with Volara^**^ (Patton & Sheridan, 2025) and Daisy^††^ (Nguyen et al., 2022). For more details see Supplementary Section 6.2.4.

### 4.5 Synaptic Reconstruction

A putative excitatory synapse was defined as colocalization of the presynaptic marker Bassoon and the postsynaptic marker PSD95 (Michalska et al., 2024; Tavakoli et al., 2025). First, we performed background removal on the Bassoon and PSD95 channels using a difference-of-gaussian approach (*σ* = 0.1 (signal) and *σ* = 8 (background)) and normalized the data to chosen global min/max values per channel (min = 0 for both channels, max = 60 for PSD95 and 70 for Bassoon). The main challenges in reconstructing the excitatory synaptic connections were the variability in intensity, density, and shape of the synaptic markers as well as the need to separate closely located structures. We therefore developed a U-Net based segmentation approach with a weighted binary cross-entropy (BCE) loss function that emphasizes object boundaries. Separate Bassoon and PSD95 networks were trained on manually-generated segmentation and contour masks. The masks served as two-channel objectives for network training, and more weight was given to the contour mask channel in the BCE loss to enforce clear separation of neighboring markers. Additionally, the semantic mask loss was weighted using an inverted distance transform, further emphasizing the boundaries of the structures. Networks were trained using Gunpowder^¶^ and Pytorch^‡^. During training, we sampled the batches containing closely-located instances more frequently, to make sure the model efficiently learned to avoid merge errors. Simple and intensity augmentation was applied to the data.

Instance segmentations were generated by postprocessing predicted semantic masks with a seeded watershed algorithm. To generate seeds for the algorithm, we first thresholded the predicted semantic segmentation (threshold = 0.5) and then computed the distance transform on the resulting thresholded network output. Next, we smoothed the resulting distance transform and computed local maxima, which were used as input for the seeded watershed algorithm. Processing of the full volume was done in a block-wise fashion and information on the detected segments (centroid and size) was stored in a graph database using Volara^**^ (Patton & Sheridan, 2025).

To detect putative excitatory synapses, we performed overlap-based matching of the detected instances after dilating each segmentation instance by one voxel. Bassoon and PSD95 instances with the largest overlap were matched together and identified as putative synapses. Here, we enforced that PSD95 could only be assigned to one Bassoon instance while Bassoon could be matched to multiple PSD95 instances. Finally, we assigned these detected putative synapses to segmented neuronal masks based on their spatial overlap (*>*20%).

Validation was performed on an additional annotated test crop from a different area of the full dataset by matching ground-truth and predicted instances based on spatial overlap. Since we focused here on mapping synapses on dendritic morphologies, only one-to-one connections were allowed for PSD95, whereas Bassoon instances could be matched to multiple ground-truth objects. For the neuron-centered validation, we first manually identified putative excitatory synapses along a segmented neurite by using Bassoon and PSD95 imaging channels. Next, these ground-truth point annotations were matched to automatically-generated putative excitatory synapses from the same region using a nearest neighbor approach^||^.

Due to the complex structure of the reconstructed thorny excrescences and their unique synaptic appearance (Figure 5.F), manual annotation of synaptic densities were performed for those sets of experiments.

## 5 Acknowledgements

We thank members of E11 Bio and the Boyden, Kornfeld, and Rodriques labs for helpful discussions; S. Collins, A. Marblestone, D. Markowitz, H. Parthasarathy, S. Smith, J. Troidl, and P. Vemuri for useful suggestions; S. Wood and A. Resasco from Crick Biological Research Facility for animal rearing and surgeries for the hippocampal dataset; H. Nagaraj and G. Papageorgiou from Crick Chemical Biology STP for acrylate synthesis; Nikon UK for imaging support throughout dataset collection; V. Holmes for project management support; Ariadne.ai for image annotation services; E. Nitishinskaya and J. Dapello for computational support. Graphics used from BioRender: https://BioRender.com/d800v2x.

## Author Contributions

B.A., E.S.B., G.M.C., T.H., J.F., J.M.R.K., K.G.C.L, A.C.P., S.Y.P., S.G.R., A.S., S.T., and J.W. conceptualized the project; B.A., J.M.R.K., D.L., K.S.L., S.Y.P., A.C.P, and S.G.R. pioneered early methods and protocols; J.Y.A., S.W.C., K.G.C.L., and M.W. designed, cloned, and tested the C-terminal eGFP barcoding system; S.Y.P. and S.T., developed the iterative MAGNIFY protocol; E.J., J.M.M., S.Y.P., and S.T. screened the MAGNIFY-compatible antibodies, S.Y.P. imaged the hippocampal dataset; K.G.C.L., A.M., and M.W. characterized the barcodes; J.W. developed the stitching and registration pipeline; H.G.J.D., K.G.C.L., C.M., J.M.M., S.Y.P, S.T., and J.W. registered the volume; W.P. and A.S. developed the barcode-augmented segmentation; W.P. and A.S. developed the barcode-augmented automatic proofreading; A.S., C.W., F.R., J.L., and J.W. developed the synapse detection pipeline and performed synaptic analysis; E.S.B., K.G.C.L, A.C.P., S.Y.P., S.G.R., A.S., S.T., and J.W. wrote the manuscript with input from all co-authors; J.Y.A. and C.M. provided operational support; E.S.B., J.M.R.K., A.C.P., and S.G.R supervised the study.

## 6.1 PRISM Multiplexing Details

### 6.1.1 Experimental details of CA3 dataset collection

Summary of imaging order, registration channels, hardware, and laser power and exposure time settings for each color channel detailed in Supplementary Table 2, Supplementary Table 4, Supplementary Table 5. Ordering of targets was optimized based on antigen sensitivity to photo-induced epitope damage and heat-strip cycles. Imaging rounds, including order of channels per round, were optimized for imaging time and fluorophore bleaching. Note that in between some strip cycles, staining and imaging of targets were split into two consecutive parts to reduce imaging time and thus risk of GLOX-catalase buffer acidification, as well as photobleaching of sensitive fluorophores (e.g. with low intrinsic fluorescence or tolerance to acidic pH) (Supplementary Table 2)

### 6.1.2 Custom antibody production and conjugations

Custom antibodies were produced by ProSci against MOON, SUN, TAG-100 (Supplementary Table 1), as there were no commercially available antibodies suitable for PRISM multiplexing. Peptide sequences KNEQELLELD-KWASL, EELLSKNYHLENEVARLKK, and EETARFQP-GYRS were generated for MOON, SUN, TAG-100, respectively, and conjugated to a cysteine residue at the N-termini or C-termini of each peptide for column purification. For each antigen, N and C-terminal conjugates were mixed 1:1 and injected into each rabbit. Collected test bleeds were validated with ELISA and affinity purified for downstream PRISM multiplexing.

To expand the number of channels imaged per stain-strip cycle, long Stokes-shift dye ATTO490LS was custom-conjugated to polyclonal secondary antibodies. Antibodies were purchased from Jackson ImmunoResearch (Supplementary Table 3) in solution in 0.01 M Sodium Phosphate, 0.25 M NaCl, pH 7.6 Buffer. Dimethyl Sulfoxide (DMSO) was added at 10% of total antibody volume and mixed by pipetting. ATTO490LS linked with NHS ester PEG was then incubated with the antibody-DMSO mixture at a 10-fold molar excess (dye:antibody-DMSO), for 30 minutes at room temperature in the dark. Afterwards, conjugates were purified via size exclusion chromatography, using a Cytiva Akta Avant FPLC instrument on a Sepax size exclusion column (SRT-C SEC-150, 5 µm, 150 A, 7.8 × 300 mm, Sepax Technologies) with 0.01 M Sodium Phosphate, 0.25 M NaCl, pH 7.6 buffer as the mobile phase.

### 6.1.3 Quantification of stripping efficiency in CA3 dataset

After each cycle of stripping, stripping was evaluated by incubating and imaging samples in GLOX buffer as outlined in Methods 2.2. In each instance, samples were first imaged with a Nikon 10X (NA = 0.45) dry lens to capture the full volume of the imaged areas alongside surrounding tissue to identify any photoinduced crosslinking. Afterwards, a full length Z-stack with Z-step size of 4 µm was acquired with Nikon Apo LWD 40x WI *λ*S DIC N2 (NA = 1.15) water immersion lens, applying the same imaging settings (i.e., laser power, imaging exposure times, channels) of those in the previous imaging(s) of the previous stain-strip cycle. The same single tile was imaged for 40X lens stripping efficiency validations throughout the dataset collection.

To quantify for stripping efficiencies between stain cycle 7 and 8, 10X downsampling was applied to imaging round tiles to match the Z-step sizes of stripping validation data. The stripping validation image was then registered with BigWarp (Fiji) with 4×4 binning for landmark placement and transformed with thin plate spline transformations at the original resolution (Bogovic et al., 2016; Schindelin et al., 2012). The matching processed imaging round and stripped tiles were then aligned and cropped prior to segmentation. Automated ROI segmentation was performed for each channel in the processed imaging tile by implementing a custom built Python pipeline using scikit-image and scipy libraries (van der Walt et al., 2014; Virtanen et al., 2020). Imaging channels were pre-processed with Gaussian smoothing (*σ* = 0.8), and binary masks were generated with Otsu thresholding (and Li’s minimum cross-entropy thresholding as a fallback). Masks were morphologically processed via opening followed by skeletonization and dilation of skeletons (structural element radius = 2 pixels). Connected component analysis was performed, before ROI filtering, in which objects with areas of 200-50,000 pixels, aspect ratio ≤50, and solidity ≥0.05, were retained.

ROIs were then applied to registered stripped images to generate paired ROIs per channel, and the centroid coordinates, area, and mean, maximum, minimum, and total intensity values were obtained per ROI per condition. Background values per channel per condition were obtained by obtaining the 5th percentile intensity value of empty regions (those outside of ROIs). Signal-to-background ratios (SBR) were calculated for each ROI in each condition, in which perfect stripping would equate to SBR of 1.

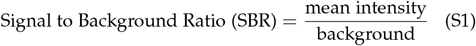

Relative stripping efficiency per ROI was then measured from SBR values, correcting for the original signal strength in the pre-stripped image (SBR_before_):

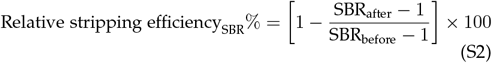

### 6.1.4 Experimental details and quantification of antibody crosstalk

Hippocampal injection samples were prepared and stained with mouse anti-PRTC, rabbit anti-MOON, and goat anti-HSV and donkey anti-mouse Cy3, donkey anti-rabbit Alexa Fluor 647, and donkey anti-goat Alexa Fluor Plus 405, as outlined in Methods 2.2, alongside ATTO488-conjugated nanobody anti-ALFA as a registration marker. Samples were incubated and imaged in GLOX buffer as detailed in Methods 2.2. To obtain pre-strip images (condition “stained,” Supplementary Figure 2.D), a single tile, full-resolution Z-stack (0.4 µm step size) was acquired in the CA3 region on the contralateral side of the stereotaxic injection, matching the spread and barcode density of that of the centerpiece dataset, with a Nikon Plan Apo LWD 40x WI *λ*S DIC N2 (NA = 1.15) water immersion lens on a Nikon-CSU Ti-2 W1 spinning disk confocal and Hamamatsu ORCA-Fusion Digital sCMOS camera (standard scan mode). Samples were then stripped as detailed in Methods 2.2, and then stained overnight with ATTO488-conjugated nanobody anti-ALFA. Afterwards, the same tile was imaged for stripping validation (condition “stripped,” Supplementary Figure 2.D) with a full-resolution Z-stack (0.4 µm step size). To test for antibody crosstalk, samples were then stained overnight with the same secondary antibodies (condition “stripped+2°,” Supplementary Figure 2.D) as in the first staining round and imaged with a full-resolution Z-stack (0.4 µm step size).

16-bit images for each condition were then 10X downsampled and trimmed in Z-axis to ensure matched Z-length and step sizes before transformation. Images of the stripped and stripped with the secondary re-stain conditions were each registered to the ALFA-ATTO488 channel of the reference image in 4×4 binning mode for landmark placement with BigWarp software in Fiji (Bogovic et al., 2016; Schindelin et al., 2012). Thin plate spline transformation was applied to full resolution and 2×2 x/y binned images of the 10X Z-downsampled stripped and stripped+2°conditions, with xy-bin corrected landmarks. Images across all conditions and xy-bin modes were then aligned and cropped.

XY-binned pre-stripped data were then used for automated ROI detection using a custom built Python pipeline using scikit-image and scipy libraries (van der Walt et al., 2014; Virtanen et al., 2020). Imaging channels were preprocessed with Gaussian smoothing (*σ* = 1.0), followed by adaptive thresholding (Li method) and morphological cleanup of small object removal (*<*100 voxels), hole filling, and binary closing (single-pixel structuring element). Watershed segmentation was then performed by applying Euclidean distance transform to binary masks, generating seed points (≥7 pixel separation, 20% maximum distance volume, ≤10,000 points), and implementing watershed algorithm with inverted distance transform. Post-segmentation processing was applied by removing gross artefacts (stripes and aggregates), ROI filtering, and merging over-segmented regions.

The final ROI masks were then applied to the stripped and stripped+2°conditions, and the centroid coordinates, area, and mean, maximum, minimum, and total intensity values were obtained per ROI per condition. Background values were obtained per condition per channel by obtaining the raw intensity value of the 5th percentile of pixels in empty regions (area outside of ROIs).

Due to fluorophore-specific differences in background accumulation and photobleaching upon imaging, stripping, or re-staining, signal to background ratio (SBR) was used as a metric robust to such variables to evaluate stripping efficiency (Supplementary Section 6.1.3). SBR values were obtained for each ROI in the stripped and stripped+2°conditions, and % crosstalk per ROI was measured by the following:

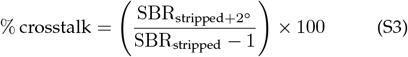

## 6.2 Volume Segmentation Details

### 6.2.1 Training Data

The fully registered target volume contained 40 channels (30 barcodes and 10 markers). We deemed 18 barcode channels to be of high enough quality to use for the segmentation pipeline. We additionally had access to two earlier datasets, which had fewer available barcodes (12 and 11, respectively). Annotations were performed by Ariadne^‡‡^. We selected crops from those volumes that contained varying structures such as cell bodies, axons, dendrites, densities, and noise. Individual neurons were sparsely reconstructed inside these crops. The five crops from the 18-channel volume are visualized in Supplementary Figure 4. Additionally, we selected crops from the 12-channel volume, which contained varying semantic structures that were then densely annotated. During training, these semantic classes were combined into a simple foreground/background such that neurons which were expressing in at least one channel were labeled as foreground. Further crop details can be seen in Supplementary Table 6, Supplementary Table 7.

### 6.2.2 Training Pipeline

All networks were trained using Gunpowder^¶^ and Py-Torch^‡^. Batches were randomly sampled from the training crops, normalized, and zero-padded to ensure valid label density (≥ 50%) to account for elastic rotations. The data was then augmented with spatial and intensity transforms (Supplementary Table 10, Supplementary Table 12, Supplementary Table 14, Supplementary Table 16). A hold-out crop was used to validate networks.

#### Enhancement

The enhancement network was trained in a channel-agnostic fashion and used all training samples as input. A probability weighting was used to select samples from the 18-channel volume more frequently. Only spatial augmentations were applied, as intensity augmentations would skew the input intensities from the target average intensities. Additionally, the difference between the average barcodes and the raw barcodes (i.e, the residual barcodes) was used as the training target (in contrast to the direct average barcodes). Validation was done using PSNR and SSIM at various checkpoints during training. These metrics both converged relatively quickly during training, and it was decided to use a later checkpoint for inference after empirically observing better diffusion of the barcode signal throughout the neurons. For network details see Supplementary Table 8.

#### Affinities and LSDs

Since this network was trained in a multi-channel fashion, we first randomly duplicated (with replacement) the sample channels (not including GFP) to the target number of channels (18) and then randomly shuffled the channels. Spatial and intensity augmentations were applied. Since the training data was sparsely annotated, a label mask (true where labels *>* 0) was used to create a scale array (via per batch inverse frequency weighting) to balance the loss between class labels. This was done independently per offset in the affinity neighborhood. We used an affinity neighborhood with two long-range offsets (6 channels), in addition to the short-range offsets (3 channels) and trained with a weighted MSE loss. The final loss is a sum of the affinities and LSDs losses. Validation was done on segmentations generated at various iterations using a variation of information (VoI), and the optimal checkpoint was used for inference. For network details see Supplementary Table 9, Supplementary Table 11.

#### Contrastive Uniform Embedding

The uniform embedding model was trained only on the 18-channel data. For the network to learn to compare specific channel intensities, we did not expand and shuffle the other datasets which contained fewer channels. Additionally, as this network is a lightweight MLP, it can be trained quickly and can therefore be dataset-specific. Thus, we did not apply spatial augmentations. Small intensity augmentations were applied to help the model generalize to slight distribution shifts in the enhanced intensities. The model was trained with a contrastive loss, constrained to labeled objects. Since small changes in the loss could lead to significant qualitative differences in the uniform embedding, but we did not have a method for turning the predictions directly into a segmentation for evaluation, the final model was chosen via manual inspection of the predictions on a held-out dataset. For network details see Supplementary Table 13.

#### Barcode Expression Probabilities

The goal of this model was to learn per channel barcode expression probabilities, which could ultimately be thresholded and combined to generate a foreground mask for postprocessing. We also trained this network in a channel-agnostic fashion and therefore trained on all samples. We trained with a binary cross-entropy loss on the max projection across channels. The large variation in intensity expression between and within channels led to excessive false positives in the background. To mitigate this, we trained with a distance mask to the nearest boundary, ignoring the loss deep within objects. Networks were validated via binary cross entropy (BCE) on a held-out dataset of semantic labels, and the final model was chosen after manual inspection of the validation results. For network details see Supplementary Table 15.

### 6.2.3 Inference

Prediction was done in a blockwise fashion using Volara^**^ (Patton & Sheridan, 2025) (and under the hood Daisy^††^ (Nguyen et al., 2022)) for each of the four described networks. Each block was processed using a Gunpowder^¶^ predict node and distributed using Volara. Blocks were grown to the maximum size that could fit in GPU memory (24 GB RAM). Inference of the full volume was then run on 175 AWS EC2 A10 GPUs. Enhancement took ∼ 3.5 hours, while other networks each took ∼ 30 minutes. All networks wrote predictions to Zarr, which were then used for downstream processing.

### 6.2.4 Post-Processing

Following block-wise inference of the above networks, we first combined the predicted affinities with barcode affinities, which were generated by taking the dot product on the predicted uniform embedding. The affinity between pixels inside the same object should be high (close to 1) while the affinity between pixels across objects should be low (close to − 1). Clipping these barcode affinities between 0 and 1 produces results that are similar to what is trained in the affinities network (Figure 3). Combining these two approaches improves cases in which one approach correctly predicts a split edge while the other one incorrectly predicts a merge edge. Here, we combined the affinities by simply multiplying the two across all neighborhoods. We also tried computing a weighted sum of the two affinities, in which we highly weight the predicted affinities in the short range (for merging) and highly weight the barcode-affinities in the long range (for splitting). The former approach (direct product) resulted in fewer false merges, so we chose this strategy as it is preferable for downstream barcode relabeling.

Since the networks were trained with sparse labels, there was no explicit training signal to supervise the background (i.e to predict zero in these regions). Masking is commonly used during segmentation post-processing pipelines to prevent errors introduced by structures not contained in neuropil (i.e. background, cell bodies, blood vessels, etc.) (Januszewski et al., 2018; Macrina et al., 2021; Sheridan et al., 2022). We similarly needed to employ a masking strategy to prevent merges against the background (or non “barcode-able” regions). We therefore created a mask of foreground regions by computing the max across channels on the predicted binary probabilities. To handle single-channel outlier values, we only considered where ≥ 2 channels were expressed rather than 1 channel. This mask was then combined with a “valid” mask denoting the boundaries of the fully registered volume. We then used the combined affinities constrained to this mask to compute a segmentation using a block-wise post-processing pipeline, similar to what was proposed by Sheridan et al., 2022.

However, rather than using a seeded watershed and hierarchical agglomeration approach, we first computed supervoxels with blocks via mutex watershed (Wolf et al., 2018), which makes use of the short-range affinities for merging (attractive edges) and long-range affinities for splitting (repulsive edges). For each of these supervoxels, we stored its center of mass and ID as nodes in a region adjacency graph (RAG). We then added edges to the region adjacency graph (RAG) between supervoxels that share affinities and assigned them merge and split weights based on the underlying affinities. We performed a final global mutex watershed on the RAG edges to get a globally optimal segmentation across blocks. For post processing parameters, see Supplementary Table 17.

### 6.2.5 Barcode Relabeling

Following post-processing to obtain a segmentation, we then developed an approach to reconnect split segments using the barcode information. As an input for this step, it was important to minimize the amount of false merges since they would incorrectly skew the computed average barcodes. Additionally, we were able to focus on reconnecting across larger distances using the barcodes, since we already agglomerated supervoxels into reasonably sized segments using the affinities.

Given a segmentation, we first created skeletons for each segment using Kimimaro (Silversmith et al., 2021), a TEASAR algorithm (Sato et al., 2000) implementation. This was done in memory on downsampled segments, (but it could also be done in a block-wise fashion at higher resolution and would thus result in more valid nodes to consider for merging). We computed an average barcode for each segment. We used a mapping from segment to supervoxels, and then computed the average intensity inside each supervoxel. The segment barcodes were generated via a weighted mean of the supervoxel barcodes to account for variance in supervoxel size. The segment barcodes were zero-mean normalized and assigned to the skeleton nodes.

We created a KDTree on end nodes and performed two spatial queries. The first spatial query fetched all pairs of nodes less than a distance match threshold (*D*_*m*_) (∼ 10 µm) apart. Nodes were then mapped back to their skeletons, leaving us with edges between skeletons that are less than *D*_*m*_ apart. We then computed affinities via cosine similarities between normalized skeleton barcodes. The second query followed the same process to get affinities between skeletons that at their closest point were less than *D*_*m*_ + *δ* apart (*D*_*s*_) (∼ 30 µm). We then filtered out any positive edges from this list to use the larger distance query purely as a splitting bias for mutex watershed. For efficiency, we also removed redundant positive edges from the second query that were present in the first smaller query. The second larger query was important to mitigate the effects of long chains of low-distance merge errors, resulting in a merge between two skeletons with a large barcode distance. Resulting edges were then clustered with mutex watershed (Wolf et al., 2018) to obtain a final lookup table, mapping original segments to barcode relabeled segments. Using this lookup table, we relabeled the input segmentation similarly to what was done for the affinity segmentation.

### 6.2.6 Evaluation

#### 6.2.6.1 Data

We created 1542 ground-truth skeletons for testing, which consisted of both large skeletons and fragments of skeletons (due to the size of the sample). The skeletons contained 206,442 nodes (average=134), a total length of 232,296 µm (biological distance, average=150 µm), and a total longest path length of 31,017 µm (avg=20 µm). See Supplementary Figure 4 for a visualization.

Due to the expansion factor and sparsity of this dataset, rather than focusing on tracing small spines, we created ground-truth skeletons that were reconstructed along the main paths of the neurons. We therefore do not consider segmentation accuracy in the context of small spines in this evaluation, but we anticipate trends to hold at higher expansion factors for smaller objects.

#### 6.2.6.2 Segmentation evaluation

##### Evaluated baselines

We conducted a “baseline” method evaluation to gauge how adding barcodes and enhancing the data improves segmentation accuracy (Supplementary Figure 9). Evaluated “methods” are described below.

- **R-GFP:** The minimal baseline considered. Since GFP expresses in every labeled cell, it is a reasonable proxy for a sparse morphology channel, or one in which every barcode was imaged in a single channel. The affinity network then sees a single channel as input.
- **R-Mean:** Computed by averaging the raw barcode intensities across channels into a single channel. Since barcodes imaged in separate channels also provide distinguishable intensity ranges, computing the mean gives a better readout than by taking data imaged in a single channel. The affinity network sees a single channel as input.
- **R-Stack:** Computed by stacking the raw barcode intensities into a multi-channel array. The affinity network then takes all channels as input, and therefore has full access to the intensities provided by imaging multiple barcodes.
- **E-Mean:** The raw barcodes are first enhanced and then the average is computed across channels. The affinity network sees a single channel as input.
- **E-Stack:** The raw barcodes are first enhanced, and then stacked across channels. The affinity network sees multi-channel input.
- **E-Stack+U:** The raw barcodes are enhanced and stacked across channels. The affinity network and uniform embedding network both see multi-channel input.

##### Evaluation Method

We evaluated increasingly large ROIs grown outward from the center of the full ROI. The largest ROI was empirically chosen to be a large enough ROI to contain a reasonable amount of ground-truth skeletons while minimizing the amount of stitching and registration artifacts. Outside of this ROI, segmentation accuracy decreases substantially. For each ROI and method, we obtained the RAG and fragments within the ROI. We cropped the ground-truth skeletons to the ROI, masked, and then relabeled connected components. We used a fixed mutex watershed bias ([-0.4, −0.7]) to generate a segmentation. This was the same bias used for generating supervoxels, as it provides a reasonable intermediate for merging and splitting. The segmentation was then evaluated against the ground-truth using standard metrics (VoI, NVoI, ERL, NERL). We also evaluated MCM on all but the largest ROI (due to memory constraints). For details, see Supplementary Figure 9, Supplementary Figure 10, Supplementary Table 18.

##### Computational costs

For each evaluated method, we computed the TeraFLOPs per µm^3^ to gauge the computational costs. We found that simply incorporating the barcodes provides an immediate increase in accuracy with little to no increase in computation (single channel vs 18 channel input). Enhancing the barcodes is then a relatively expensive operation (which scales by the number of barcodes), but is necessary for the increased accuracy and for training the subsequent embedding network (which greatly benefits from clean input data). The final embedding then allows for further accuracy increases (and split/merge tradeoffs) with little increased computational cost (Supplementary Figure 11).

#### 6.2.6.3 Barcode relabeling evaluation

We evaluated barcode relabeled segmentation on a smaller candidate evaluation ROI, which contained a relatively dense quantity of ground-truth skeletons with obvious splits in the initial affinity segmentation. We evaluated standard metrics (VoI, NVoI, ERL, NERL) in a grid evaluation to assess the effects of the number of bits, match threshold, and spatial distance threshold. We evaluated random subsets of bits for matching and averaged across 10 runs. We found that matching accuracy increases with the number of barcodes used to match. Additionally we found the optimal matching threshold to be around 0.9, and the optimal inner spatial distance threshold to be 50000 nanometers, see Supplementary Figure 13 for details.

##### Gap crossing evaluation

###### Evaluation Method

We then did a gap crossing evaluation to see how well we could reconnect segments across increasingly large spatial gaps using the barcodes. Here, we used the fragments generated from the E-M+U network since it had minimal false merges. We evaluated the gap crossing on the same increasingly large ROIs as were used for the segmentation evaluation. For each ROI we evaluated how well we could match across regions of missing data. ROI evaluation was performed by chunking the ROI into blocks of size (full z shape, full y shape, fragment block size in x). We then iterated over all pairs of chunks going from the easy cases (two neighboring chunks) to the hardest case (two chunks on opposite sides of the evaluation ROI). In each chunk we assumed perfect segmentation according to the intersecting ground-truth skeletons and computed the mean barcode for each skeleton in each chunk via agglomerating supervoxels. We then zero-mean normalized the computed intensities. Using these normalized intensities, we performed a pairwise matching on embedding distance with a threshold, enforcing at most one edge per node in a chunk, and no edges between nodes in the same chunk. Finally we computed the F1 score of the matched segments against the ground-truth. The mean scores across all pairs of chunks was reported for each evaluation ROI.

###### Evaluation Results

We found that across the board, the matching accuracy decreases as spatial distance increases, as expected. The raw barcodes yielded the best results (over the enhanced and discrete barcodes). The small improvement over enhanced barcodes is likely due to the smoothing losing important information for matching (especially in fine processes). Similarly, the discrete barcodes remove even more information that would be useful for matching (by effectively turning the continuous signal into binary probabilities). Notably, matching accuracy increases with the number of barcodes used for matching which makes sense considering collisions become rarer as the number of barcodes increases. Additionally, matching accuracy increases when fewer skeletons are considered. This is important to understand when considering that high F1 scores (close to 1) at low distances may be due to simply not having that many skeletons to match together. See Supplementary Figure 15 for details.

---

**Supplementary Figure 1:**
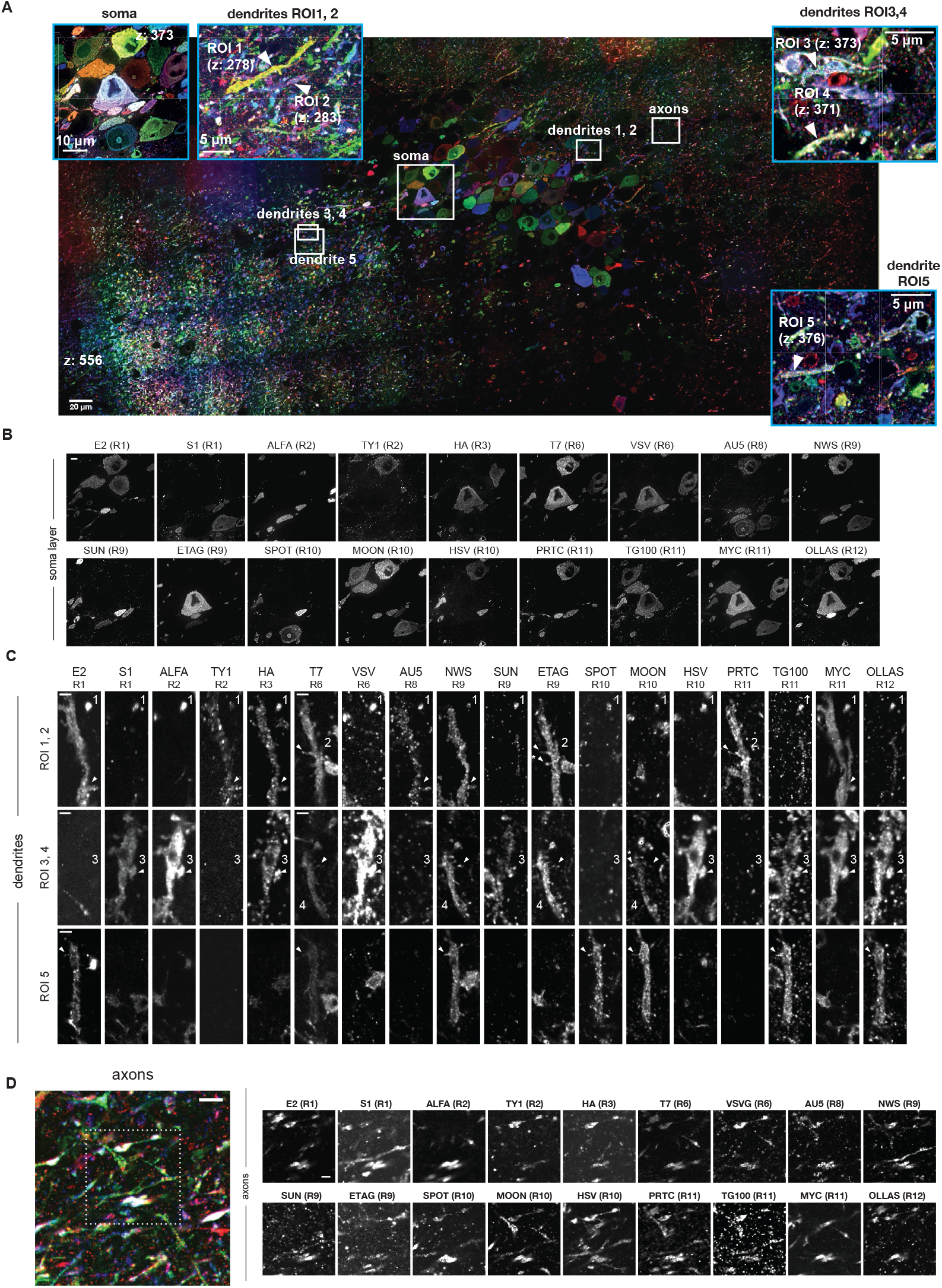
Demonstration of multiplexing and signal detection of all 18 protein bits in mouse hippocampal CA3 dataset. A. Overview of entire FOV of stitched and registered raw mouse hippocampal CA3 dataset. B. Single-plane raw images demonstrating detection of all 18 protein bits across 11 imaging rounds in the soma layer. Scale bar 20 µm. C. Single-plane raw images demonstrating detection of all 18 protein bits across five dendrites. Arrowheads point to the same dendritic spine for each dendrite. Arrowheads with asterisks are different dendritic spines containing detectable protein bit signal, due to signal in the former spine being out of focus. Scale bar 2 µm. D. Single-plane raw images demonstrating detection of all 18 protein bits in axons. (Left) A zoom-in of an axon-rich region indicated in overview image (A). Dotted box demarcates ROI of breakout panel on right. Scale bar 2 µm. (Right) Breakout panel of zoomed-in single-plane images per protein bit channel. Scale bar 1 µm. All scale bars correspond to pre-expansion distances.

**Supplementary Figure 2:**
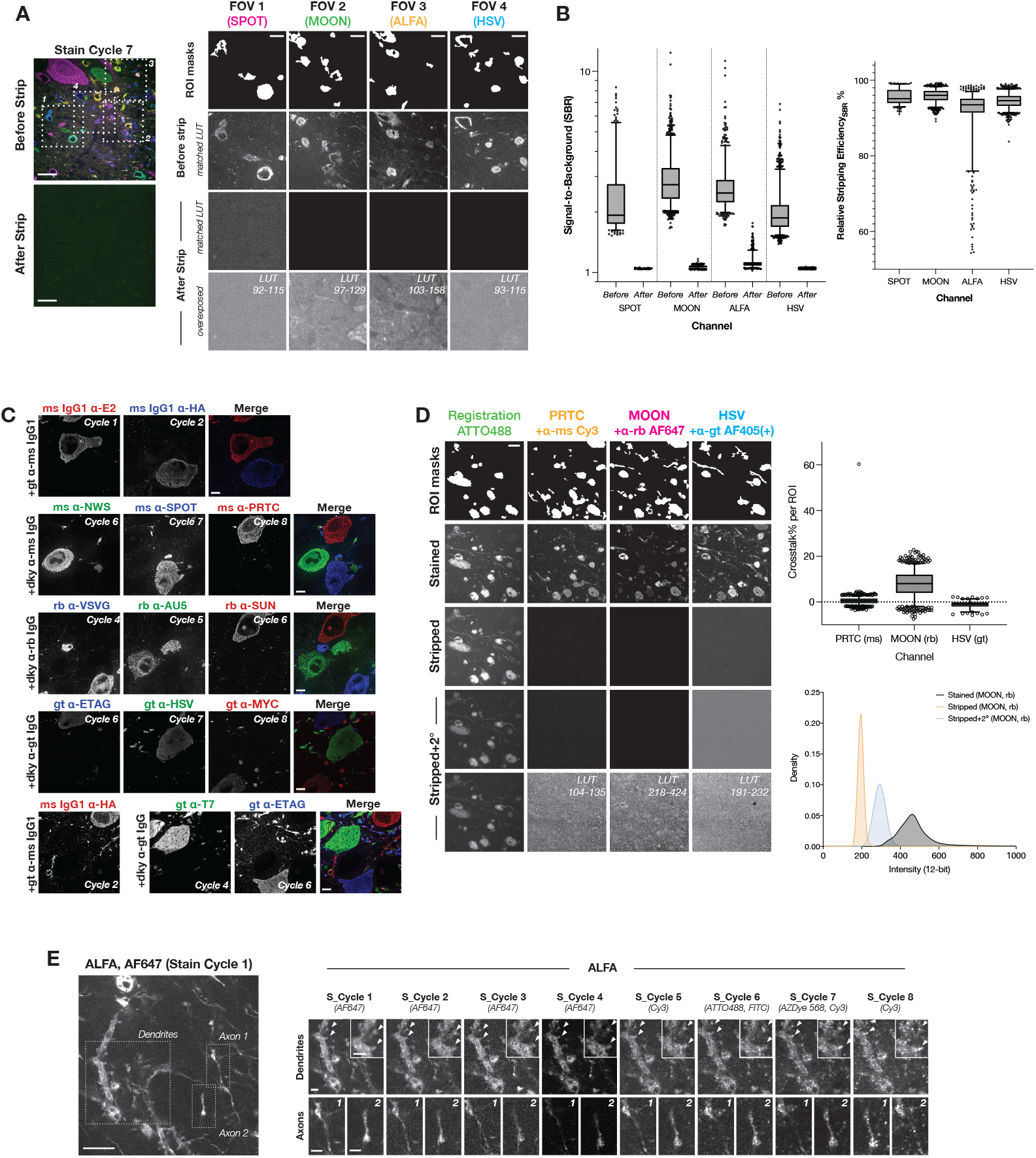
Validation of PRISM multiplexing strategy for mouse hippocampal CA3 dataset collection. A. Representative images showing high stripping efficiency after cycle 7 in a hippocampal dataset. (Left) Full-field single-plane before (top) and after (bottom) stripping, acquired under identical imaging conditions and LUTs. Post-strip images were transformed by BigWarp registration (Supplemental Methods). (Middle) Magnified regions from boxed areas in (left). Top row: ROI masks detected pre-strip for quantification in B. Middle rows: matching LUT images before and after stripping. Bottom row: same post-strip images shown overexposed. LUT min/max values (12-bit scale) indicated at right. Scale bars, 4 µm. B. Quantification of stripping efficiency after cycle 7. (Left) Per-channel boxplots of signal-to-background ratio (SBR) before and after stripping across all ROIs (Supplementary Material 5.1.3). Median [IQR], *n* ROIs: SPOT 95.1 [94.0–97.4], *n* = 309; MOON 96.0 [94.7–97.1], *n* = 1092; ALFA 93.5 [91.5–95.1], *n* = 690; HSV 94.6 [93.2–95.8], *n* = 1191. (Right) Boxplots of relative stripping efficiency (%) across ROIs (Supplementary Material 5.1.3). Outliers outside the 5–95th percentile are shown as individual points. C. Example regions in raw hippocampal dataset demonstrating minimal antibody crosstalk across rounds. Single-plane images show non-overlapping signal between (top four rows) protein bits stained with same-species primary antibodies and corresponding same-species secondary antibodies, or (bottom) a protein bit, HA, stained with secondary antibodies raised in goat while protein bits in later rounds were detected with anti-goat secondary antibodies. Scale bars: 20 µm. D. Demonstration of minimal antibody crosstalk with PRISM multiplexing. Registration channel is an ATTO488-conjugated nanobody against ALFA. (Left) Representative images of the crosstalk experiment (Supplemental Methods) in the CA3 region of a separate barcoded sample. All post-strip images were transformed using BigWarp registration. Scale bar: 4 µm. (Right, top) Boxplots of % crosstalk per matching ROI (Supplementary Methods 5.1.4). Median [IQR], *n* ROIs: PRTC 0.5 [™ 0.5–1.5], *n* = 954; MOON 8.1 [4.0–12.0], *n* = 817; HSV ™ 1.0 [™ 2.0– ™0.1], *n* = 163. (Right, bottom) ROI-independent pixel intensity histogram of MOON across conditions, indicating slightly higher apparent crosstalk due to global intensity increases from addition of AF647-conjugated anti-rabbit secondary on stripped samples. X-axis limited to 0–1000 (1171 datapoints excluded). E. Strip cycle-by-cycle comparison of protein bit ALFA registration channel staining. (Left) Overview of a neurite-rich region imaged before strip cycle 1, with boxed regions highlighting individual neurites shown at right. Scale bar: 4 µm. (Right) Zoom-ins of single-plane, non-registered images of a dendrite (top row) and two individual axons (bottom row) across seven stripping cycles. Dendrite images include insets of two dendritic spines (arrowheads). Axons are labeled “1” and “2” in the top-right corner of each image. Fluorophores on secondary antibodies are indicated for each cycle (AF647 = Alexa Fluor 647). Scale bars: 1 µm. All scale bars correspond to pre-expansion distances.

**Supplementary Figure 3:**
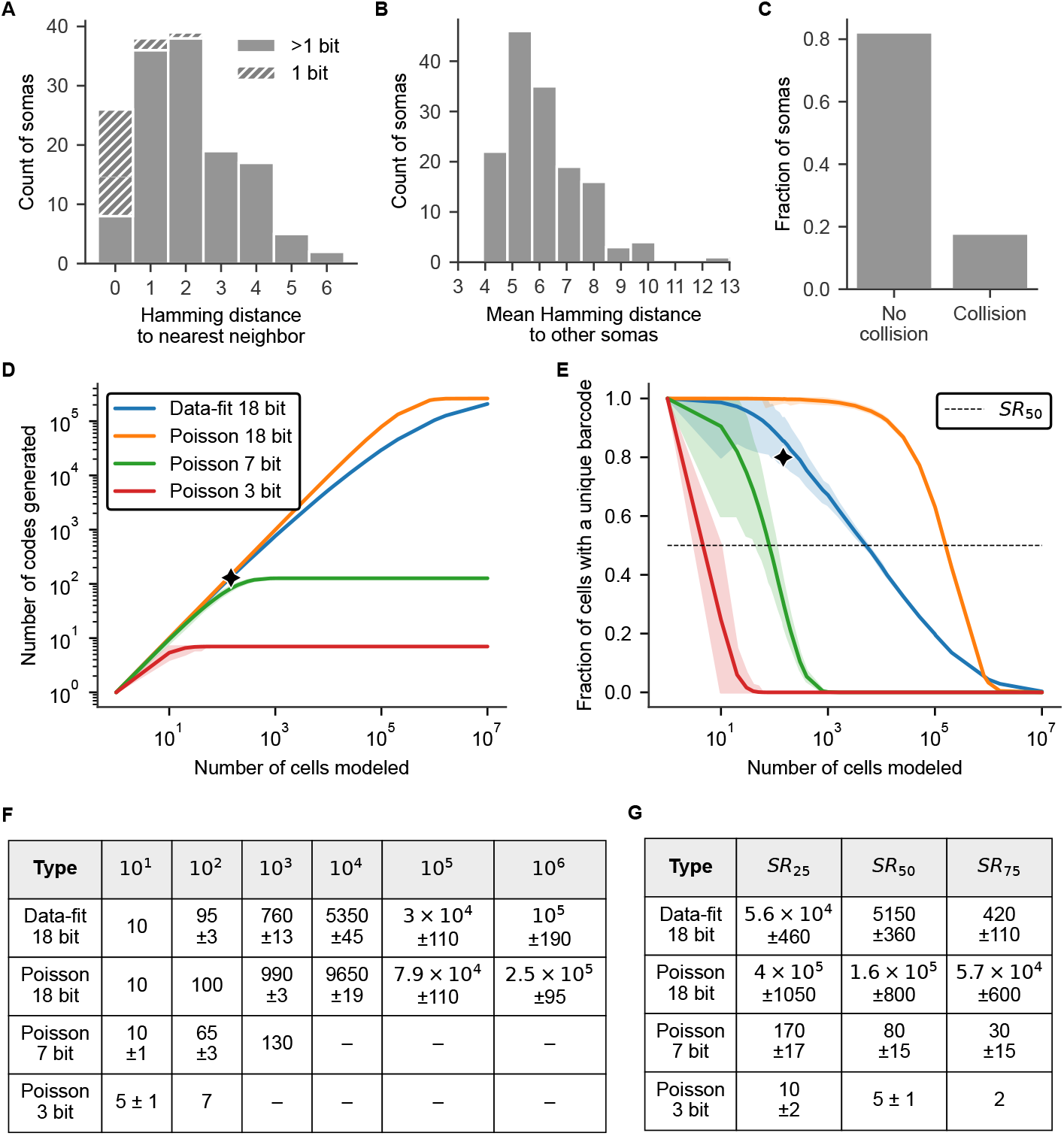
Detailed barcode statistics and simulations. A. Hamming distances per soma to nearest neighbor (mean = 3.2). The majority of cells with a collision are single-protein-bit cells (18/26, 69.2%). B. Mean pairwise Hamming distances between all somas in the dataset (mean = 6.1). C. Frequency of barcode collisions. Proportion of soma barcodes colliding with 1 or more other somas (unique =.82, non-unique =.18) D. Monte Carlo estimates of binary barcode generation. Simulations estimate the total number of barcodes generated under four models. Data-fit is parametrized by the observed epitope frequencies and barcode length distribution. Three cases represent best-case scenarios for binary combinatorial labels with proteins using 3, 7 or 18 bits. The ideal models assume equal protein bit usage with a Poisson-istributed barcode length centered at *n/*2. Simulations do not incorporate signal intensity information, as all methods benefit at the same rate. E. Monte Carlo estimates of cell uniqueness. Using the same models as in D, we calculated the fraction of unique cells at a given size up to 10^6^ total cells. 95% confidence intervals shown as shaded regions. F. Barcode generation across modeled systems. Representative number of barcodes generated across several simulated orders of magnitude. Reported values are mean ± standard deviation; data is a subset from all data shown in panel D. To improve readability, small numbers (*<*100) are shown as integers, medium numbers (100–10,000) are rounded to the nearest 10 or 50, and very large numbers (*>*10,000) are displayed in scientific notation. – indicates saturation at the previous value. Singlet rate (SR) across modeled systems. The SR limit (*SR*_*p*_) indicates the number of cells that can be labeled while retaining a p% unique fraction. Limits (*SR*_25_, *SR*_50_, *SR*_75_) were estimated from Monte Carlo simulations (5,000 iterations per condition per number of samples). Values are reported as mean ± standard deviation.

**Supplementary Figure 4:**
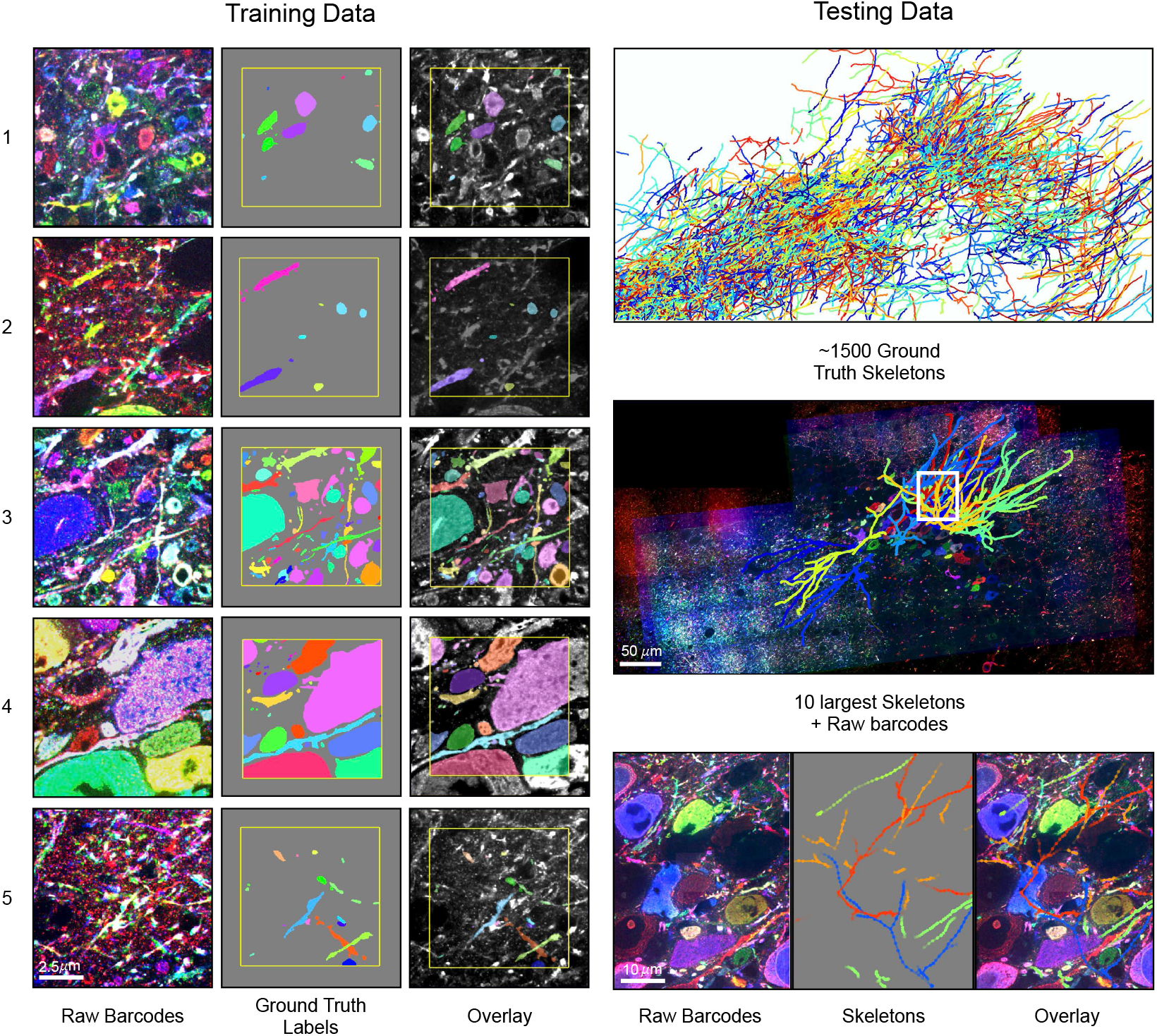
Neuron segmentation ground-truth data overview. Training data shows five crops used to train networks from the target 18 channel dataset (from left to right: raw barcodes, sparse ground-truth labels, overlay). Additional crops from other 12 channel datasets were also used but omitted for clarity. Testing data shows 1500 ground-truth skeletons used for evaluation and the largest 10 skeletons overlaid on the raw data. Zoom in shows example skeletons over raw data.

**Supplementary Figure 5:**
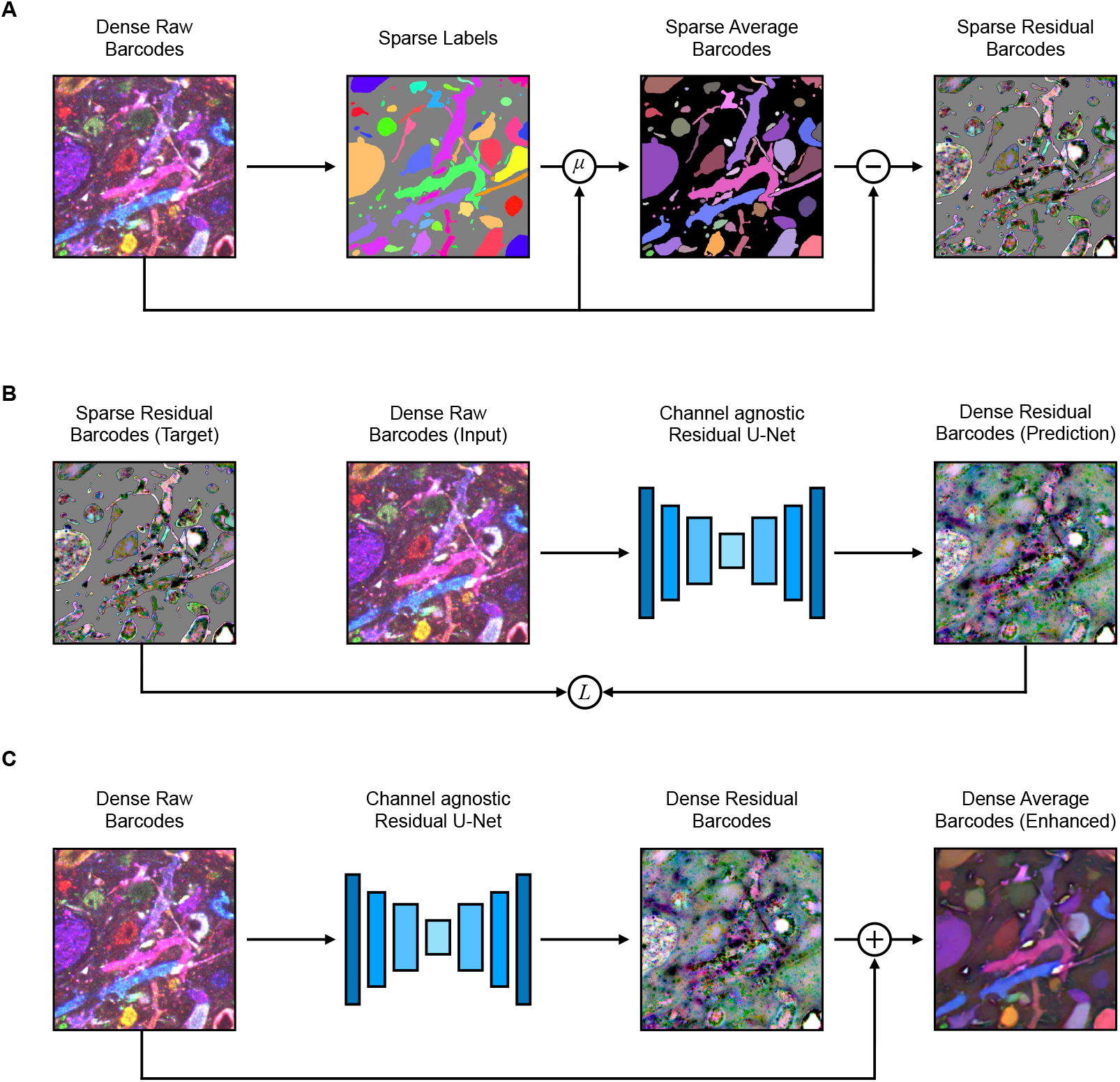
Enhancement network overview. **A**. Target data generation. Given sparse ground-truth labels and raw barcodes, sparse average barcodes are computed for each label. The raw barcodes are then subtracted from the average barcodes to generate residual barcodes. **B**. Training objective. A channel agnostic UNet is trained to learn residual barcodes from raw barcode input. **C**. Inference. Residual barcodes are predicted and then added to the raw barcodes to obtain average barcodes (enhanced barcodes).

**Supplementary Figure 6:**
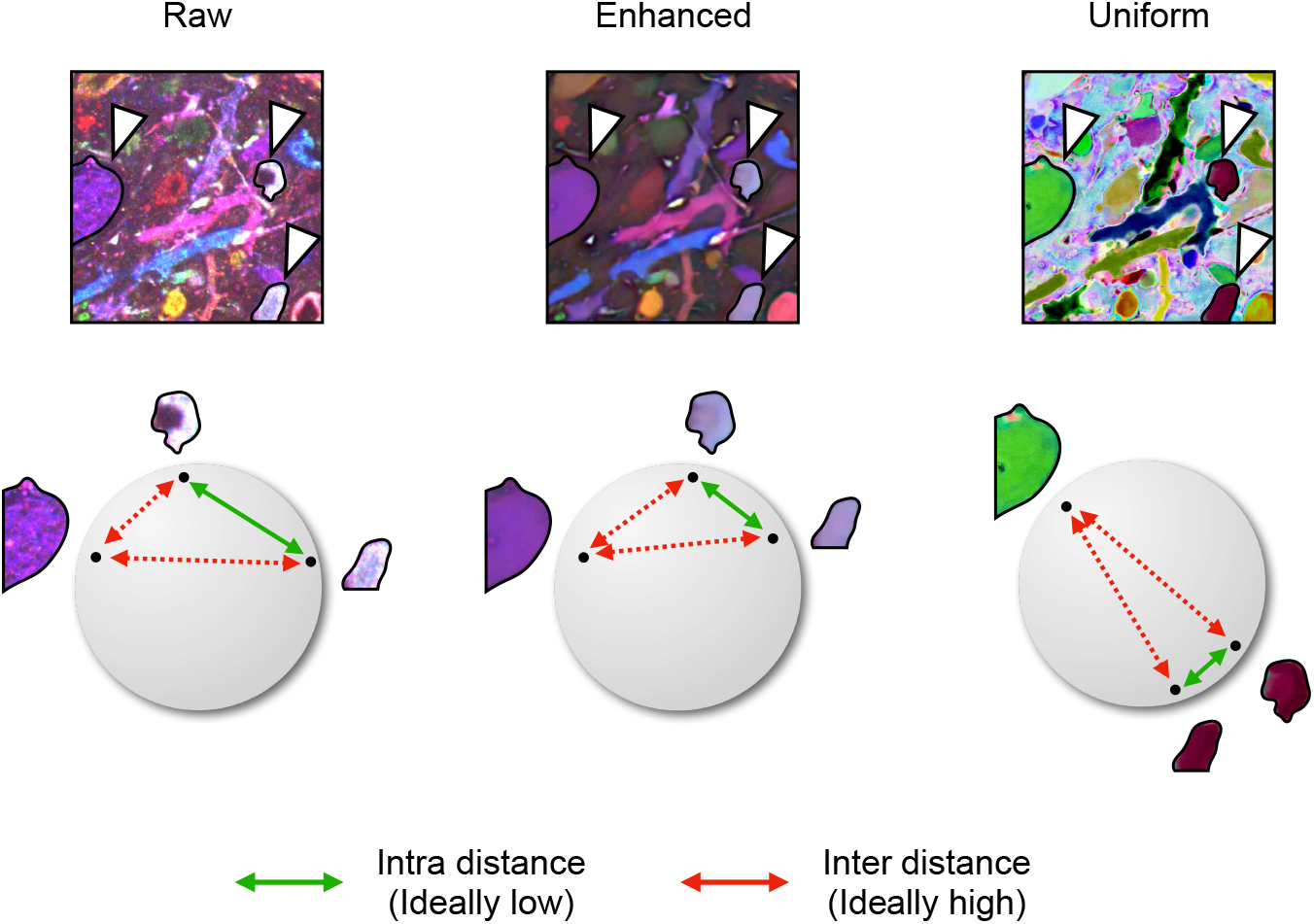
Uniform embedding objective. The uniform embedding (24 channels) aims to optimize two objectives: minimizing the intra-object distance while maximizing the inter-object distance. This is done by projecting the barcodes onto a unit sphere. Raw and enhanced data (18 channels) are visualized on unit sphere here for clarity. Image data RGB-color encoded for visualization.

**Supplementary Figure 7:**
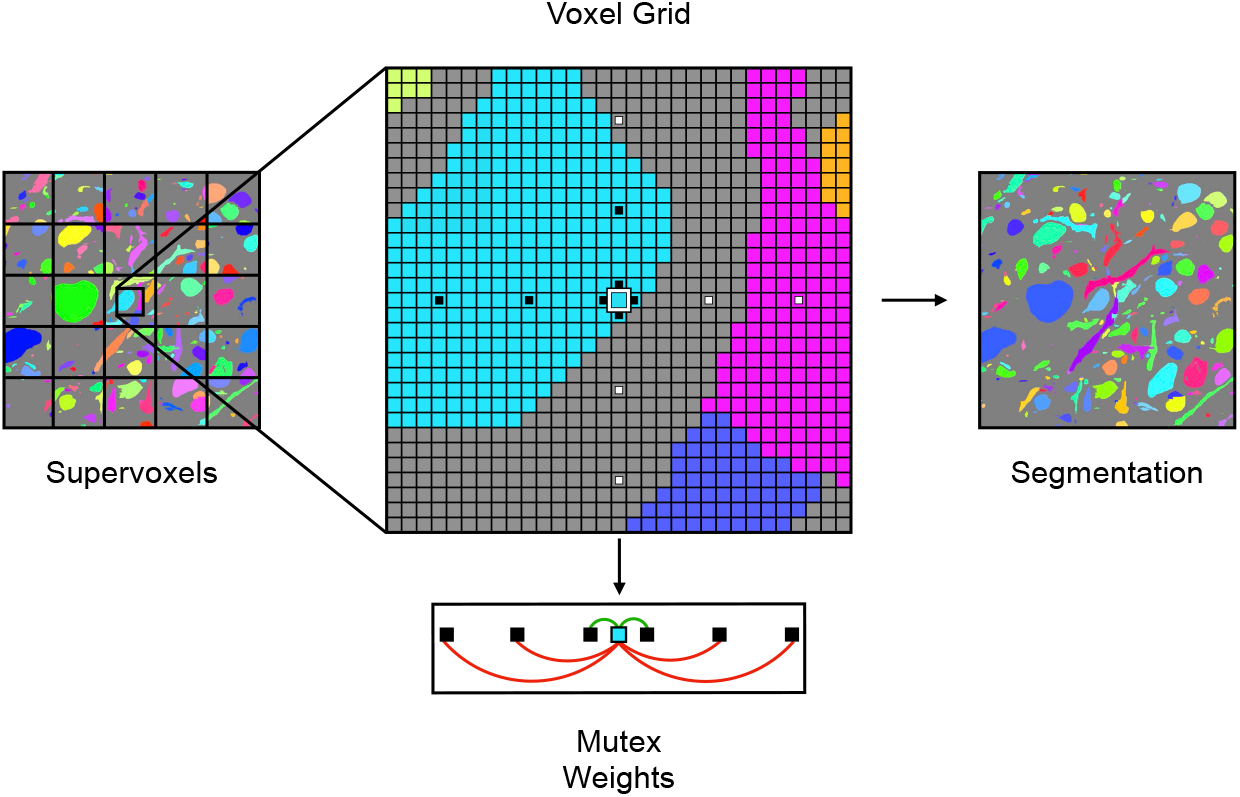
Mutex watershed overview. Weights are determined by affinity neighborhood. Short range edge affinities are used as attractive weights (green, merge signal), while long range edge affinities are used as repulsive weights (red, split signal).

**Supplementary Figure 8:**
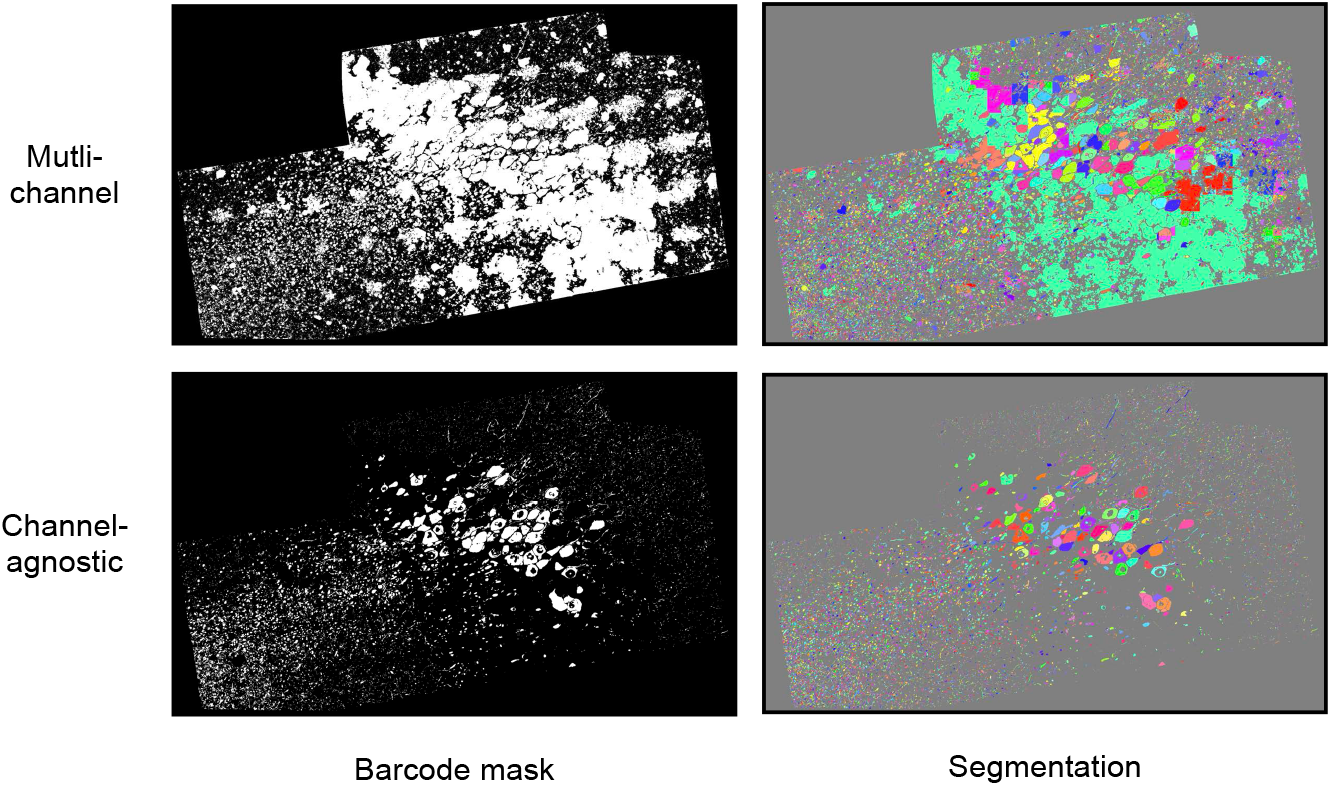
Masking effect on segmentation quality. Initial masking with a multi-channel network contained many false positives, even when randomly shuffling channels, likely due to large variability in cross channel and volume intensity ranges. This led to significant false merges to the background. A channel agnostic approach yielded better masking, and subsequent segmentation.

**Supplementary Figure 9:**
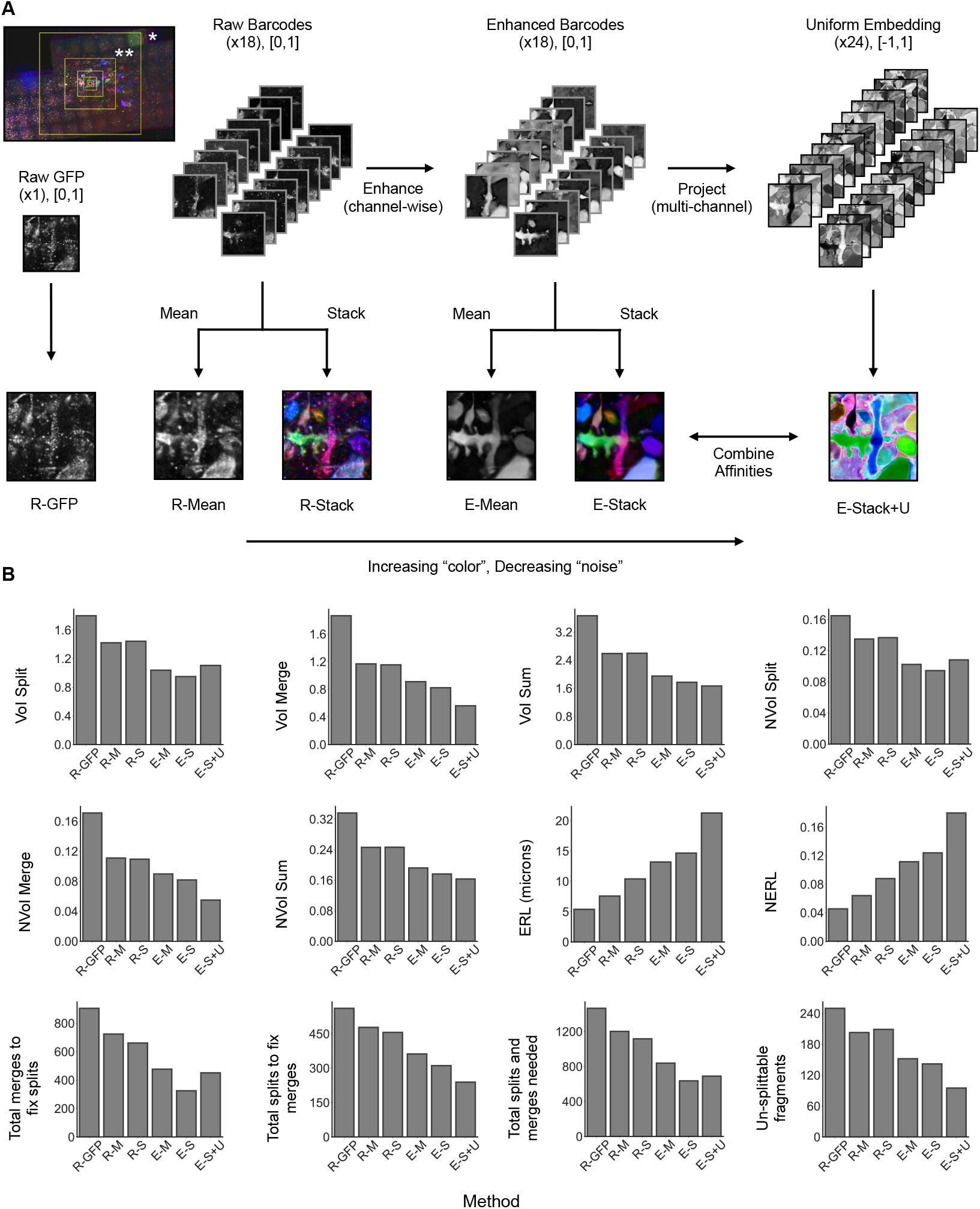
Segmentation Evaluation method and results. **A**. Overview of methods, demonstrating increasing “color” information from barcodes and decreasing “noise” from enhancement. Inset in top left shows overview of increasingly large ROIs used for evaluation. **B**. Evaluation results. Top two row metrics were evaluated up to and shown on largest ROI (asterisk in inset). Bottom row metrics (min-cut metric related) were evaluated up to and shown on second to largest ROI (double asterisk in inset) due to computational costs.

**Supplementary Figure 10:**
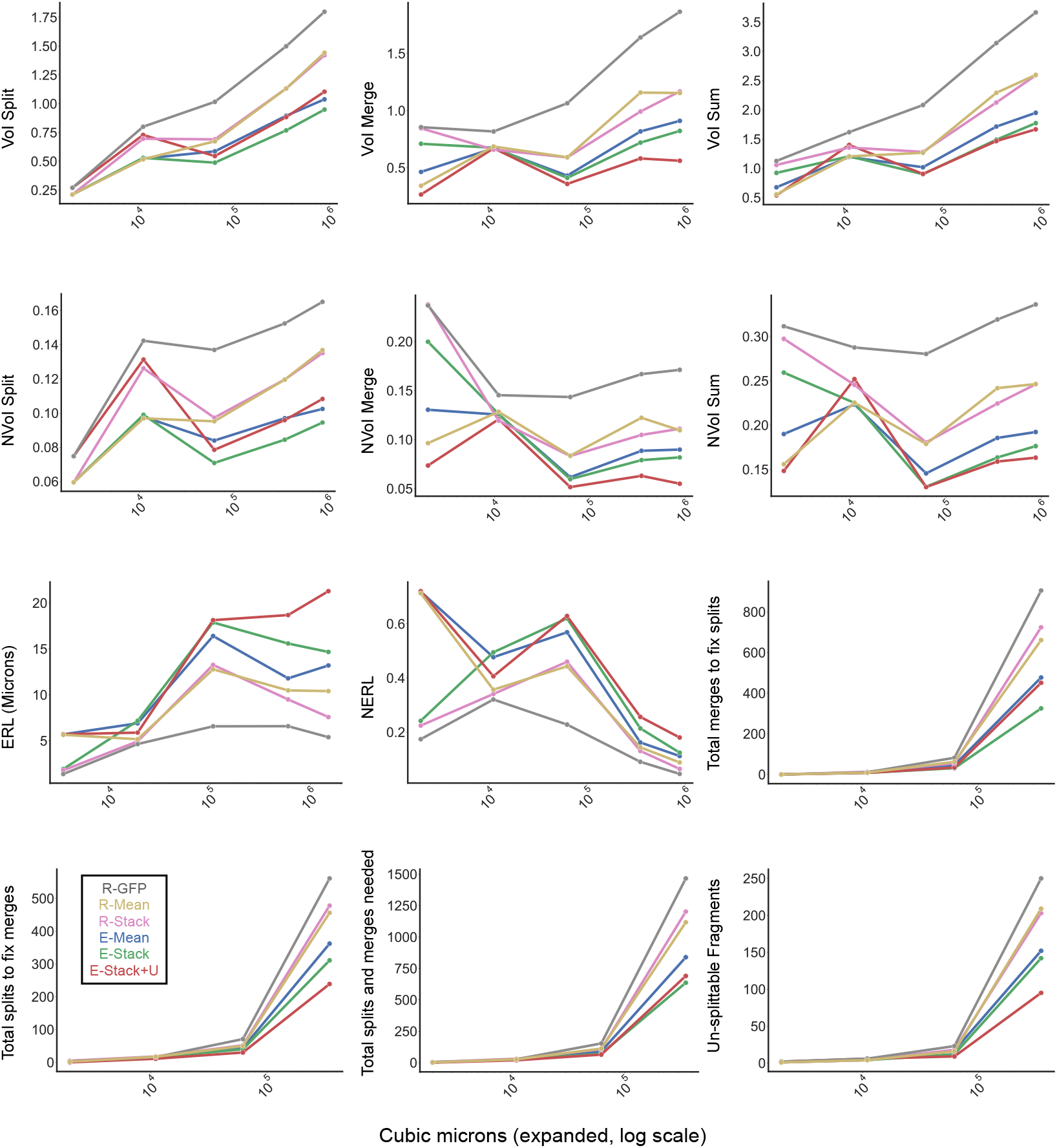
Segmentation Evaluation ROI evaluation. For each “method”, metric accuracy is shown against volume size (log scale). The first three rows were evaluated up to largest ROI (asterisk in Supplementary Figure 9.B). Bottom row metrics (min-cut metric related) were evaluated up to second to largest ROI (double asterisk in Supplementary Figure 9.B) due to computational costs.

**Supplementary Figure 11:**
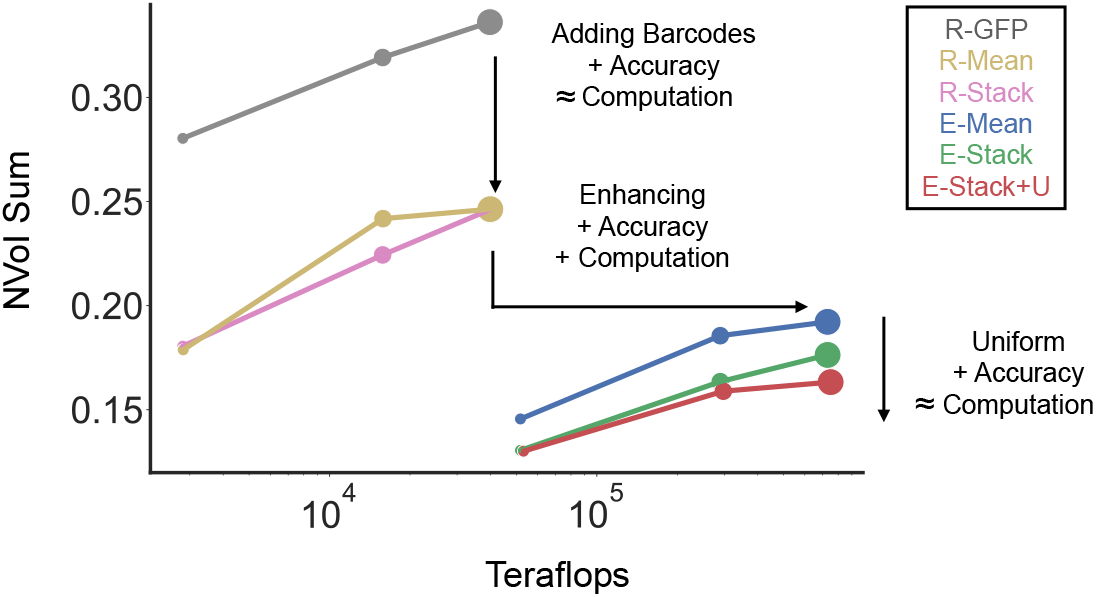
Computational costs of evaluated methods. For each “method”, accuracy (NVoI Sum is shown against Teraflops for the largest three evaluated ROIs (plotted on a log scale; dot sizes correspond to ROIs), computed using the method from (Sheridan et al., 2022). Simply adding raw barcodes provides a significant accuracy increase over raw GFP at no increased computational cost. Further accuracy increases are possible at an increased computational cost due to the enhancement (which scales with number of channels). A final accuracy increase (and further merge/split tuning can be achieved with the uniform embedding at no extra computational cost. All components are still computationally “cheap”, relatively speaking - every network was trained on a single GPU and easily distributed over modest GPUs during inference.

**Supplementary Figure 12:**
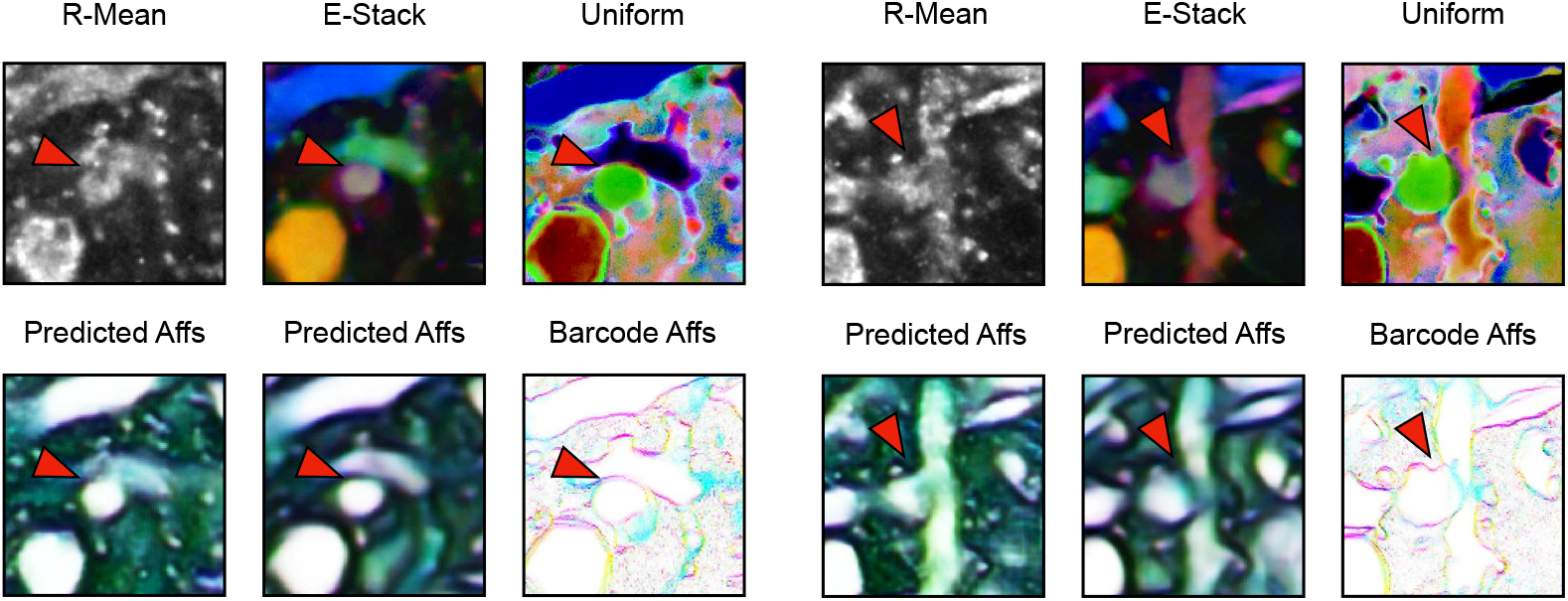
Example affinities generated from different methods. Red arrow shows two examples in which the affinities from the single channel raw barcode input (R-Mean) struggle to form a boundary between two neurons, which would likely lead to a downstream false merge. This boundary is resolved in the affinities from the enhanced and uniform networks.

**Supplementary Figure 13:**
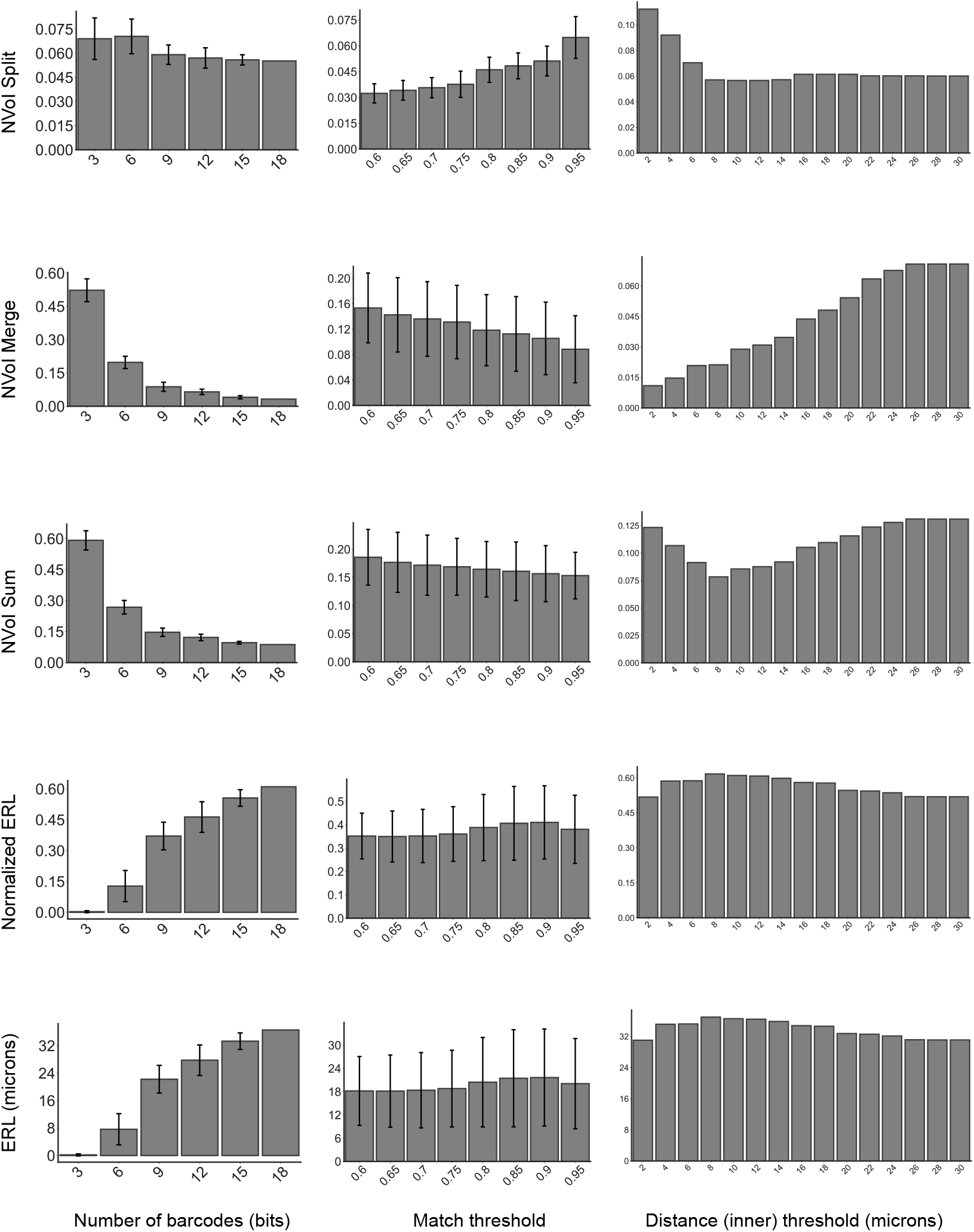
Barcode relabeling ablations. Evaluation metrics plotted against conditions from ablation grid search. From left to right: segmentation accuracy increases with number of barcodes used to relabel segments, optimal barcode distance matching threshold is around 0.9, optimal spatial matching threshold is around 40k nanometers. Except for the value they are varying, each column by default uses 18 barcodes, a match threshold of.9 and an inner distance threshold of ∼10 µm.

**Supplementary Figure 14:**
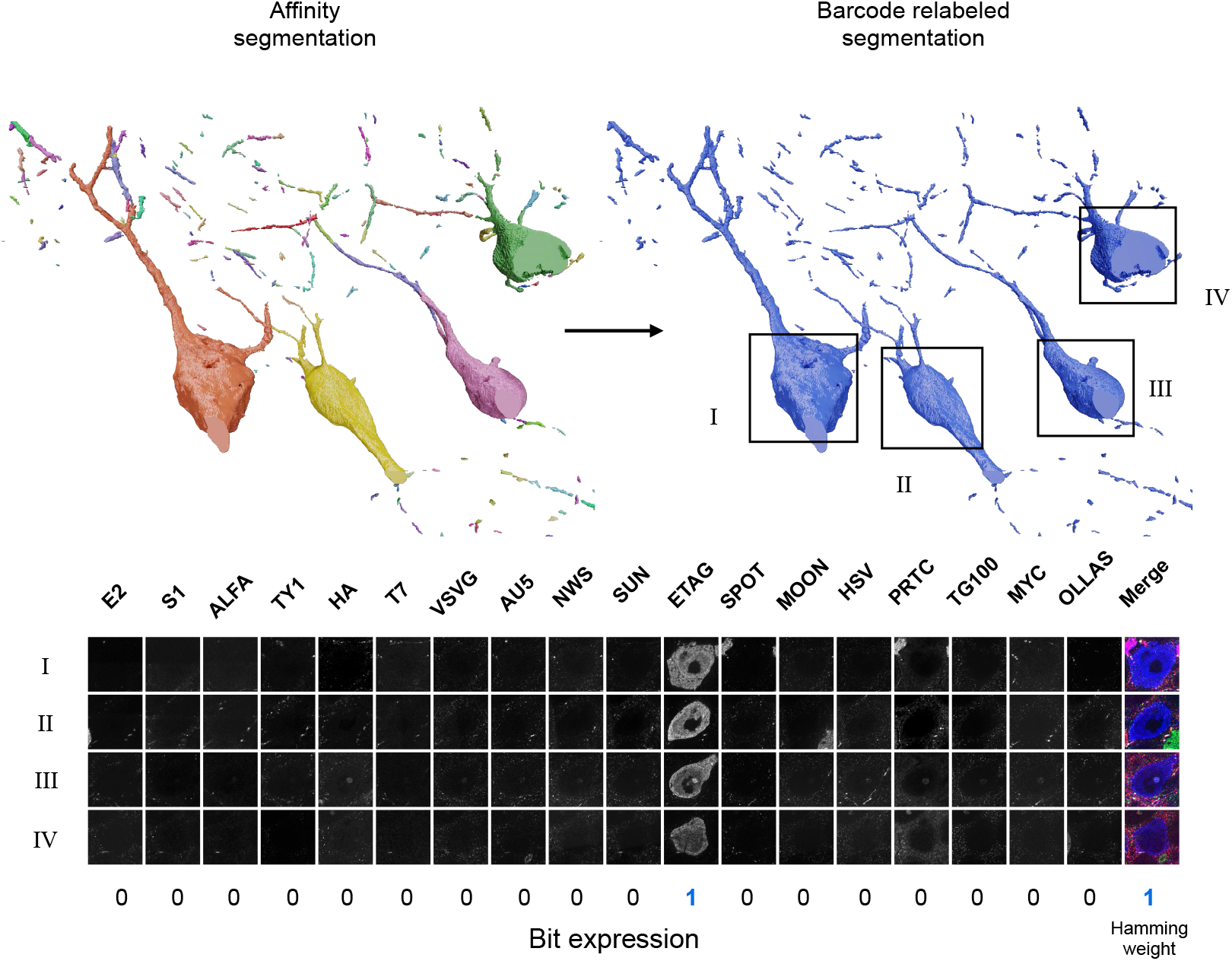
Example barcode relabeling failure case. Four neurons are incorrectly split in the affinity segmentation. The barcode relabeling falsely merges these into one segment. Looking at the underlying raw barcode channel data at the cell bodies shows a collision in barcode space, making this merge unavoidable. While it would be trivial to resolve these cases if they were restricted to cell bodies, collisions in axons/dendrites are more difficult to split without global shape priors.

**Supplementary Figure 15:**
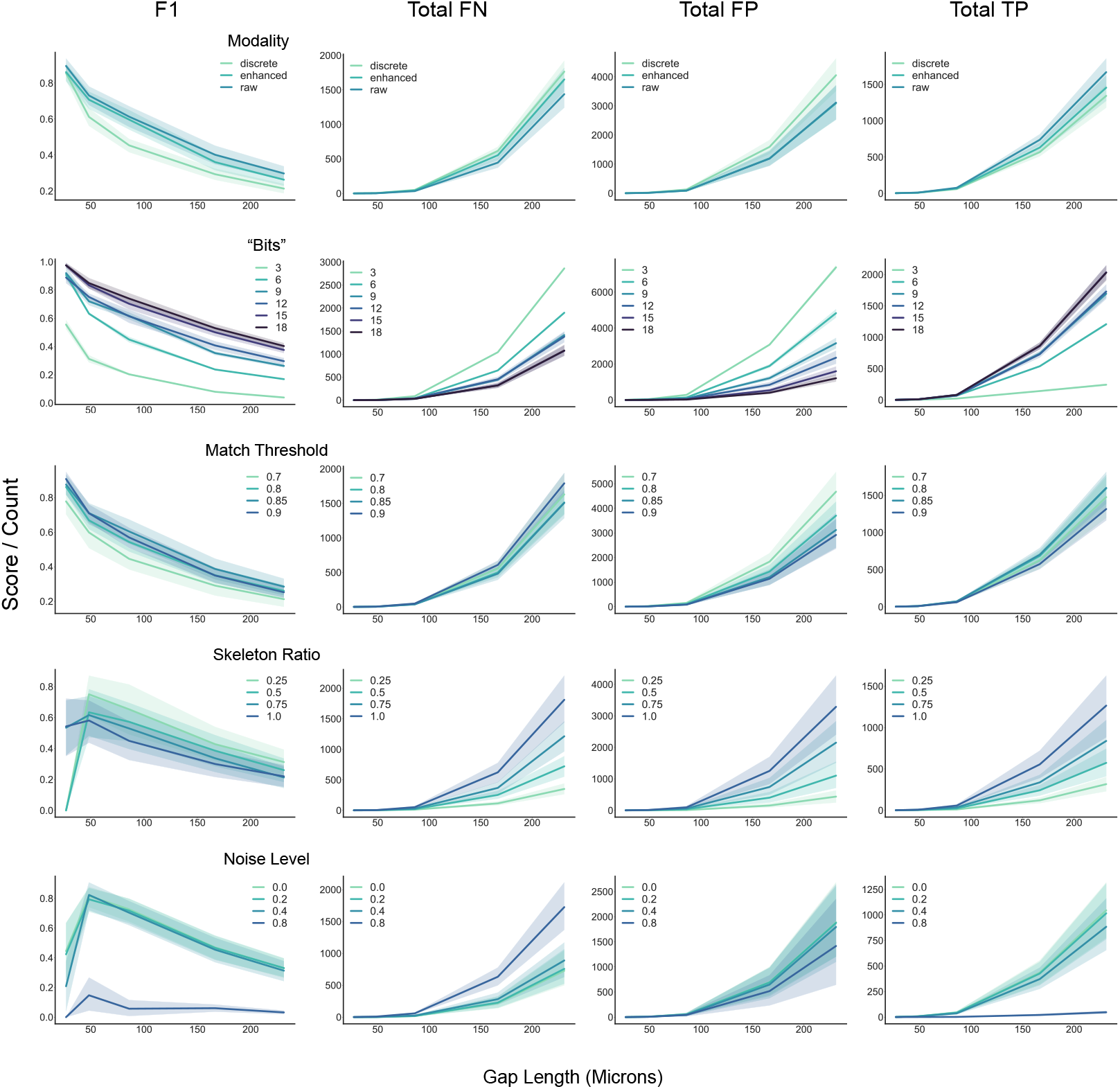
Gap crossing ablations. Each column corresponds to an evaluated metric. The score/count is shown against the gap length (in µm) for matching. Each row corresponds to an evaluated condition used for matching. From top to bottom: intensity modality, number of barcode “bits”, matching threshold, skeleton ratio (i.e the percent of skeletons used for evaluation), noise level.

**Supplementary Table 1:**
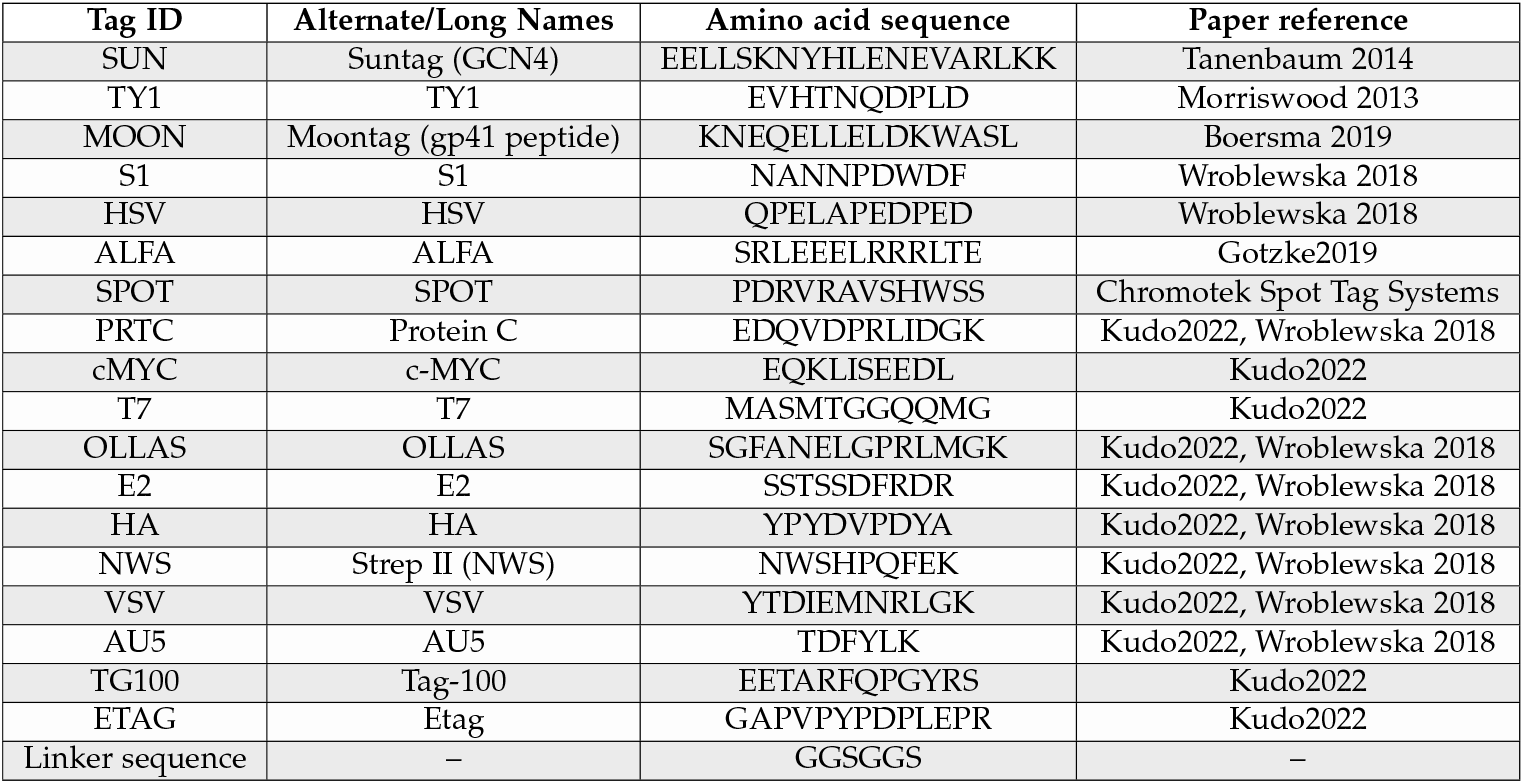
Tag names and amino acid sequences for the tags used in the study to create protein bits.

**Supplementary Table 2:**
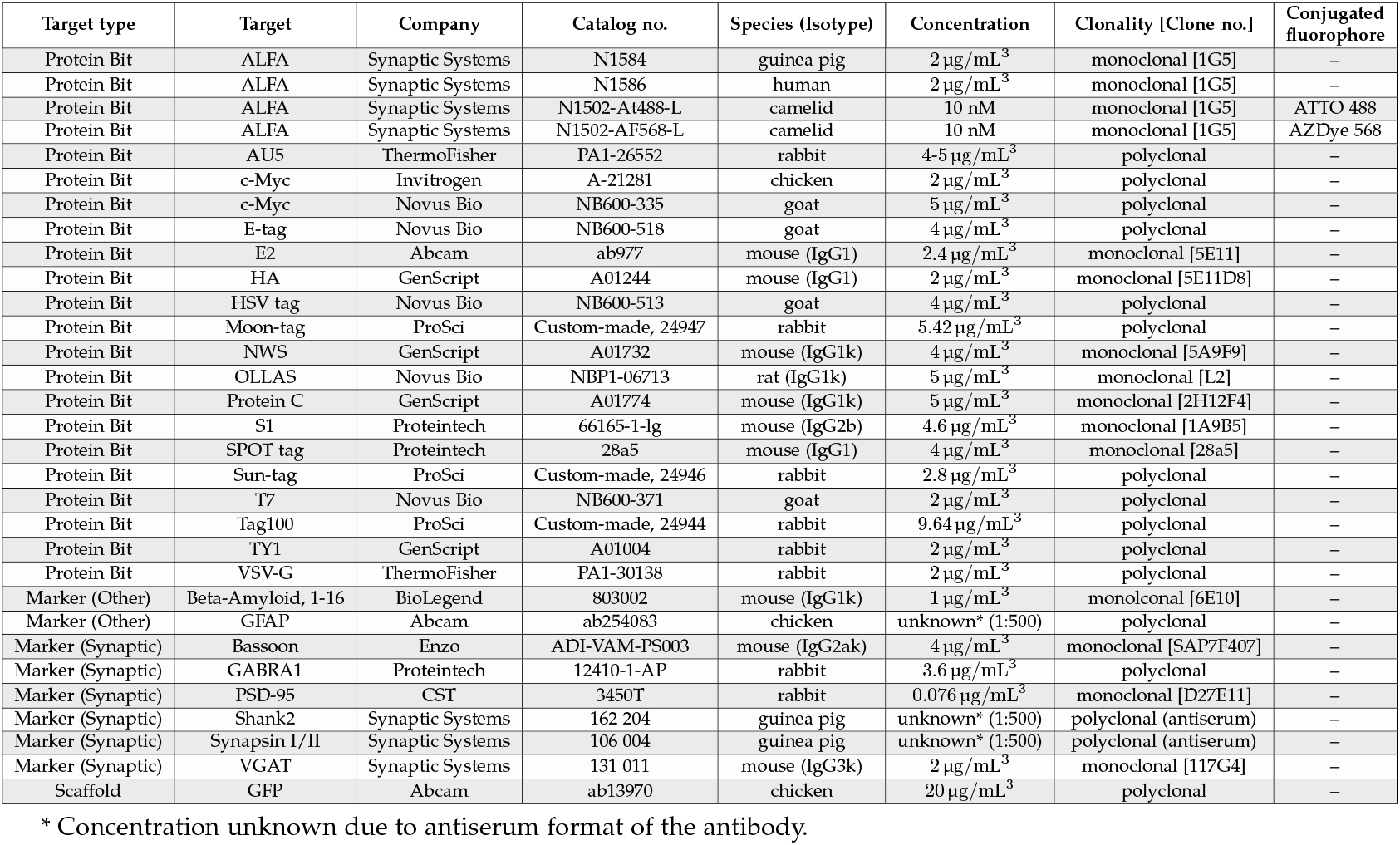
Validated primary antibodies for PRISM multiplexing against targets in mouse CA3 hippocampal dataset. All primary antibodies are unconjugated and stained with a fluorophore-conjugated secondary antibody (see Supplementary Table 4), except for those indicated in “Conjugated Fluorophore” column.

**Supplementary Table 3:** Imaging and staining round-up of all imaged targets in mouse CA3 hippocampal dataset. Round of staining specified by stain-strip cycle. Per stain-strip cycle, one or more imaging cycles were performed, imaging either the entirety or subsets of stained targets. Stain-strip round-up of targets were optimized based on antigen sensitivity to photo-induced epitope damage and heat-strip cycles. Imaging rounds, including order of channels per round, were optimized for imaging time and fluorophore bleaching.

| Stain-strip cycle | Imaging Round | Target type | Registration? | Target | Fluorophores | Channel order †† | Imaging settings † | In dataset? |
| --- | --- | --- | --- | --- | --- | --- | --- | --- |
| 1 | 1 | Protein Bit |  | E2 | ATTO488, FITC | 0 | 488 | Y |
| 1 | 1 | Protein Bit |  | S1 | Alexa Fluor 555, AZDye 568 | 1 | 555 | Y |
| 1 | 1 | Protein Bit | [R] | ALFA | Alexa Fluor 647 | 2 | 647 | Y |
| 1 | 2 | Protein Bit |  | TY1 | Alexa Fluor Plus 405 | 0 | 405 | Y |
| 1 | 2 | Protein Bit | [R] | ALFA | Alexa Fluor 647 | 1 | 647 | – |
| 1 | 2 | Marker (Other) |  | GFAP | ATTO 490LS | 2 | 490LS | N |
| 2 | 3 | Marker (Synaptic) |  | VGAT | FITC | 0 | 488 | Y |
| 2 | 3 | Synaptic |  | GABRA1 | Alexa Fluor Plus 405 | 1 | 405 | Y |
| 2 | 3 | Protein Bit |  | HA | Alexa Fluor 555, AZDye 568 | 2 | 555 | Y |
| 2 | 3 | Protein Bit | [R] | ALFA | Alexa Fluor 647 | 3 | 647 | – |
| 3 | 4 | Marker (Synaptic) |  | PSD-95 | FITC | 0 | 488 | Y |
| 3 | 4 | Marker (Synaptic) |  | Bassoon | Alexa Fluor 555 | 1 | 555 | Y |
| 3 | 4 | Protein Bit | [R] | ALFA | Alexa Fluor 647 | 2 | 647 | – |
| 3 | 5 | Marker (Other) |  | Beta-Amyloid | ATTO 490LS | 0 | 490LS | N |
| 3 | 5 | Protein Bit |  | c-Myc | Alexa Fluor Plus 405 | 1 | 405 | N |
| 3 | 5 | Protein Bit | [R] | ALFA | Alexa Fluor 647 | 2 | 647 | – |
| 4 | 6 | Protein Bit |  | T7 | FITC | 0 | 488 | Y |
| 4 | 6 | Protein Bit | [R] | ALFA | Alexa Fluor 647 | 1 | 647 | – |
| 4 | 6 | Marker (Synaptic) |  | Synapsin I/II | Cy3 | 2 | 555 | Y |
| 4 | 6 | Protein Bit |  | VSV-G | Alexa Fluor Plus 405 | 3 | 405 | Y |
| 5 | 7 | Marker (Synaptic) |  | Shank2 | FITC | 0 | 488 | N |
| 5 | 7 | Marker (Synaptic) |  | Bassoon | Alexa Fluor 555 | 1 | 647 | N |
| 5 | 7 | Protein Bit | [R] | ALFA | Cy3 | 2 | 555 | – |
| 5 | 8 | Protein Bit | [R] | ALFA | Cy3 | 0 | 555 | – |
| 5 | 8 | Protein Bit |  | AU5 | Alexa Fluor Plus 405 | 1 | 405 | Y |
| 6 | 9 | Protein Bit | [R] | ALFA | ATTO488, FITC | 0 | 488 | – |
| 6 | 9 | Protein Bit |  | NWS | Alexa Fluor 647 | 1 | 647 | Y |
| 6 | 9 | Protein Bit |  | Sun-tag | Alexa Fluor Plus 555, Cy3 | 2 | 555 | Y |
| 6 | 9 | Protein Bit |  | E-tag | Alexa Fluor Plus 405 | 3 | 405 | Y |
| 7 | 10 | Protein Bit |  | SPOT tag | Alexa Fluor 555 | 0 | 488 | Y |
| 7 | 10 | Protein Bit |  | Moon-tag | Alexa Fluor 647, Cy5 | 1 | 647 | Y |
| 7 | 10 | Protein Bit | [R] | ALFA | AZDye 568, Cy3 | 2 | 555 | – |
| 7 | 10 | Protein Bit |  | HSV tag | Alexa Fluor Plus 405 | 3 | 405 | Y |
| 8 | 11 | Protein Bit |  | Protein C | Alexa Fluor 488 | 0 | 488 | Y |
| 8 | 11 | Protein Bit |  | Tag100 | Alexa Fluor 647, Cy5 | 1 | 647 | Y |
| 8 | 11 | Protein Bit | [R] | ALFA | Cy3 | 2 | 555 | – |
| 8 | 11 | Protein Bit |  | c-Myc | Alexa Fluor Plus 405 | 3 | 405 | Y |
| 9 | 12 | Protein Bit |  | OLLAS | Alexa Fluor 488, FITC | 0 | 488 | Y |
| 9 | 12 | Scaffold | [R] | GFP | Alexa Fluor 647 | 1 | 647 | Y |
| 9 | 12 | Protein Bit |  | AU5 | Alexa Fluor 555, AZDye 568, Cy3 | 2 | 555 | N |
| 9 | 12 | Protein Bit |  | c-Myc | Alexa Fluor Plus 405 | 3 | 405 | – |

[R] Registration channel
Y Yes
N No
– Not in dataset but used for registration
† Imaging settings (See Supplementary Table 4)
†† Channels in zero order

**Supplementary Table 4:**
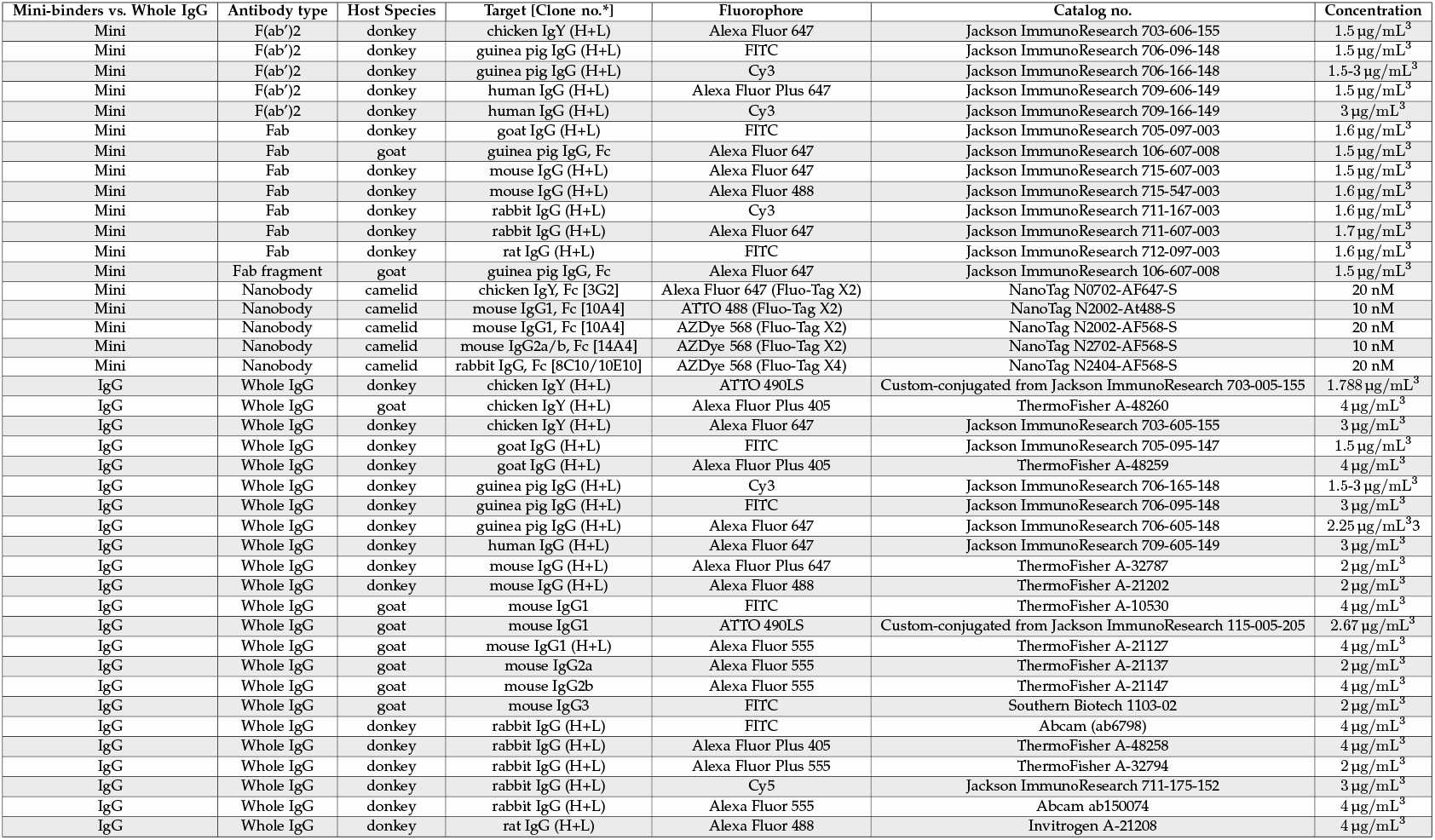
List of all validated secondary antibodies for PRISM multiplexing against targets in mouse CA3 hippocampal dataset. Minibinders were used for more efficient penetration or whole IgGs when minibinders were not available.

**Supplementary Table 5:** Imaging settings on spinning disk confocal for mouse CA3 hippocampal dataset. Details of laser excitation and fluorophore emission capture by channels (listed in “Imaging Settings” column in Supplementary Table 2). Light engine for all channels was the Omicron LightHUB Ultra, and all emission filters had a 25 nm diameter. Note that Long Stokes shift dye ATTO490LS (“490LS” channel) is excited by the 488 nm laser but captured by a Far Red emission filter.

| Channel Name | Laser Excitation |  | Emission |  |  |
| --- | --- | --- | --- | --- | --- |
|  | λ (nm) | Max Power (mW) | λ (nm) | Bandpass filter type | Manufacturer / Catalog no. |
| 405 | 405 | 120 | 447/60 | BrightLine single-pass | Semrock / FF02-447/60-25 |
| 488 | 488 | 200 | 525/50 | BrightLine single-pass | Semrock / FF01-525/50-25 |
| 490LS | 488 | 200 | 708/75 | BrightLine single-pass | Semrock / FF01-708/75-25 |
| 555 | 561 | 150 | 600/52 | BrightLine single-pass | Semrock / FF01-600/52-25 |
| 647 | 638 | 200 | 708/75 | BrightLine single-pass | Semrock / FF01-708/75-25 |

**Supplementary Table 6:** Training crops for instance segmentation.

| Crop | Dataset | Channels | Raw Shape | Labels Shape | #Labels |
| --- | --- | --- | --- | --- | --- |
| 0 | 231220 | 18 | (300, 650, 650) | (200, 500, 500) | 23 |
| 1 | 231220 | 18 | (300, 650, 650) | (200, 500, 500) | 13 |
| 2 | 231220 | 18 | (300, 650, 650) | (200, 500, 500) | 225 |
| 3 | 231220 | 18 | (300, 650, 650) | (200, 500, 500) | 44 |
| 4 | 231220 | 18 | (300, 650, 650) | (200, 500, 500) | 27 |
| 5 | 230701 | 12 | (300, 300, 300) | (300, 300, 300) | 233 |
| 6 | 230701 | 12 | (450, 750, 1600) | (450, 750, 1600) | 112 |
| 7 | 230701 | 12 | (500, 1000, 1000) | (468, 908, 908) | 5 |
| 8 | 230701 | 12 | (500, 1000, 1000) | (468, 908, 908) | 20 |
| 9 | 230701 | 12 | (468, 908, 908) | (468, 908, 908) | 26 |
| 10 | 230701 | 12 | (500, 1000, 1000) | (468, 908, 908) | 27 |
| 11 | 230701 | 12 | (500, 1000, 1000) | (468, 908, 908) | 42 |
| 12 | 230703 | 11 | (468, 908, 908) | (468, 908, 908) | 16 |
| 13 | 230703 | 11 | (468, 908, 908) | (468, 908, 908) | 16 |

**Supplementary Table 7:** Training crops for semantic segmentation (and binary expression).

| Crop | Dataset | Channels | Raw Shape | Labels Shape | Classes |
| --- | --- | --- | --- | --- | --- |
| 0 | 231220 | 18 | (50, 300, 300) | (50, 300, 300) | 4 |
| 1 | 231220 | 18 | (50, 300, 300) | (50, 300, 300) | 7 |
| 2 | 230701 | 12 | (65, 300, 300) | (65, 300, 300) | 6 |
| 3 | 230701 | 12 | (50, 300, 300) | (50, 300, 300) | 6 |
| 4 | 230701 | 12 | (50, 300, 300) | (50, 300, 300) | 6 |
| 5 | 230701 | 12 | (50, 300, 300) | (50, 300, 300) | 5 |
| 6 | 230701 | 12 | (50, 300, 300) | (50, 300, 300) | 5 |
| 7 | 230701 | 12 | (50, 300, 300) | (50, 300, 300) | 4 |
| 8 | 230701 | 12 | (50, 300, 300) | (50, 300, 300) | 4 |
| 9 | 230701 | 12 | (50, 300, 300) | (50, 300, 300) | 4 |
| 10 | 230701 | 12 | (50, 600, 600) | (50, 600, 600) | 4 |
| 11 | 230701 | 12 | (50, 300, 300) | (50, 300, 300) | 3 |
| 12 | 230701 | 12 | (50, 600, 600) | (50, 600, 600) | 7 |
| 13 | 230701 | 12 | (50, 600, 600) | (50, 600, 600) | 5 |
| 14 | 230701 | 12 | (50, 300, 300) | (50, 300, 300) | 5 |
| 15 | 230701 | 12 | (50, 300, 300) | (50, 300, 300) | 5 |
| 16 | 230701 | 12 | (50, 300, 300) | (50, 300, 300) | 2 |

**Supplementary Table 8:** Enhancement network parameters.

| Parameter | Value |
| --- | --- |
| Model type | UNet |
| Input feature maps | 12 |
| Layer fmap scale | 5 |
| Downsampling factors | [(1, 2, 2), (1, 2, 2), (2, 2, 2)] |
| Upsample Mode | Trilinear |
| Residual blocks | Yes |
| Input shape | [52, 148, 148] |
| Output shape | [20, 56, 56] |
| Loss | MSE |
| Optimizer | Adam |
| Learning Rate | 0.5e-4 |
| Betas | (0.9, 0.999) |
| Epsilon | 1x10-8 |
| Iterations | 150000 |
| Checkpoint | 60000 |

**Supplementary Table 9:**
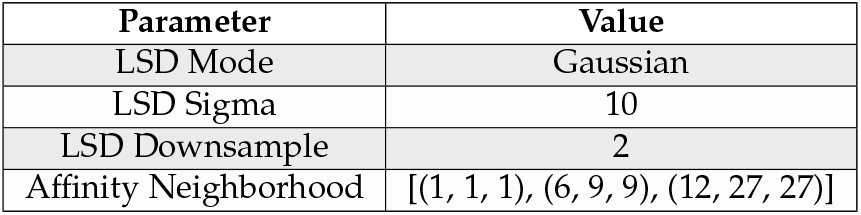
Affinity and LSD parameters.

**Supplementary Table 10:**
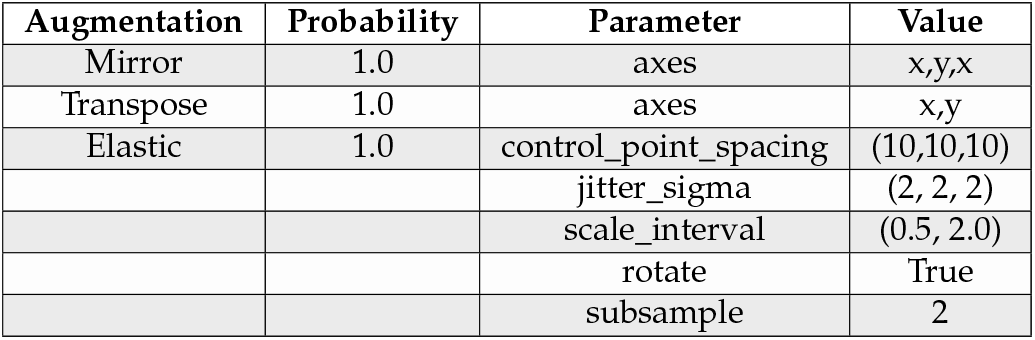
Enhancement network training augmentations.

**Supplementary Table 11:** Affinity network training parameters.

| Parameter | Value |
| --- | --- |
| Model type | UNet |
| Input feature maps | 12 |
| Layer fmap scale | 5 |
| Downsampling factors | [(1, 2, 2), (1, 2, 2), (2, 2, 2)] |
| Upsample Mode | Nearest |
| Residual blocks | No |
| Input shape | [76, 196, 196] |
| Output shape | [44, 104, 104] |
| Loss | MSE |
| Optimizer | Adam |
| Learning Rate | 0.5e-4 |
| Betas | (0.9, 0.999) |
| Epsilon | 1x10-8 |
| Iterations | 150000 |
| Checkpoint | 140000 |

**Supplementary Table 12:** Affinity network training augmentations.

| Augmentation | Probability | Parameter | Value |
| --- | --- | --- | --- |
| Shuffle | 1.0 |  |  |
| Mirror | 1.0 | axes | x,y,x |
| Transpose | 1.0 | axes | x,y |
| Elastic | 0.7 | control_point_spacing | (10,10,10) |
|  |  | jitter_sigma | (2, 2, 2) |
|  |  | scale_interval | (0.5, 2.0) |
|  |  | rotate | True |
|  |  | subsample | 4 |
| Gaussian Blur | 0.3 | range | (0.3, 1.2) |
| Intensity | 0.8 | scale | in [0.9, 1.2] |
|  |  | shift | in [-0.1, 0.2] |
|  |  | clip | True |
|  |  | channel_wise | True |
| Section Defects | 0.2 | prob_missing | 0.1 |
|  |  | prob_low_contrast | 0.1 |
|  |  | contrast_scale | 0.5 |
| Shape Defects | 0.3 | num_channels | 6 |

**Supplementary Table 13:** Contrastive embedding network training parameters.

| Parameter | Value |
| --- | --- |
| Model type | MLP |
| Hidden features | 1024 |
| Layers | 6 |
| Input shape | [1, 212, 212] |
| Output shape | [1, 212, 212] |
| Loss | Contrastive |
| Optimizer | Adam |
| Learning Rate | 0.001 |
| Betas | (0.9, 0.999) |
| Epsilon | 1x10-8 |
| Weight Decay | 0.0001 |
| Batch Size | 10 |
| Iterations | 10000 |
| Checkpoint | 10000 |

**Supplementary Table 14:** Contrastive embedding network training augmentations.

| Augmentation | Probability | Parameter | Value |
| --- | --- | --- | --- |
| Mirror | 1.0 | axes | x,y |
| Transpose | 1.0 | axes | x,y |
| Elastic | 0.2 | control_point_spacing | (10,10,10) |
|  |  | jitter_sigma | (2, 2, 2) |
|  |  | scale_interval | (0.8, 1.2) |
|  |  | rotate | True |
|  |  | subsample | 2 |
| Intensity | 0.8 | scale | in [0.9, 1.1] |
|  |  | shift | in [-0.2, 0.2] |
|  |  | clip | True |

**Supplementary Table 15:**
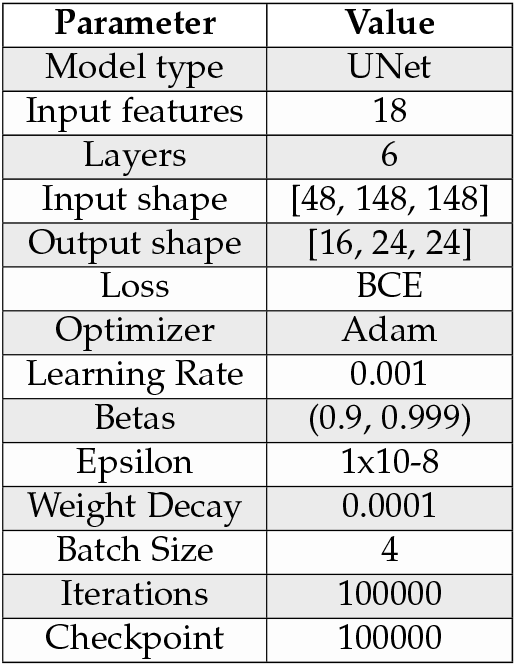
Binary expression network training parameters.

**Supplementary Table 16:**
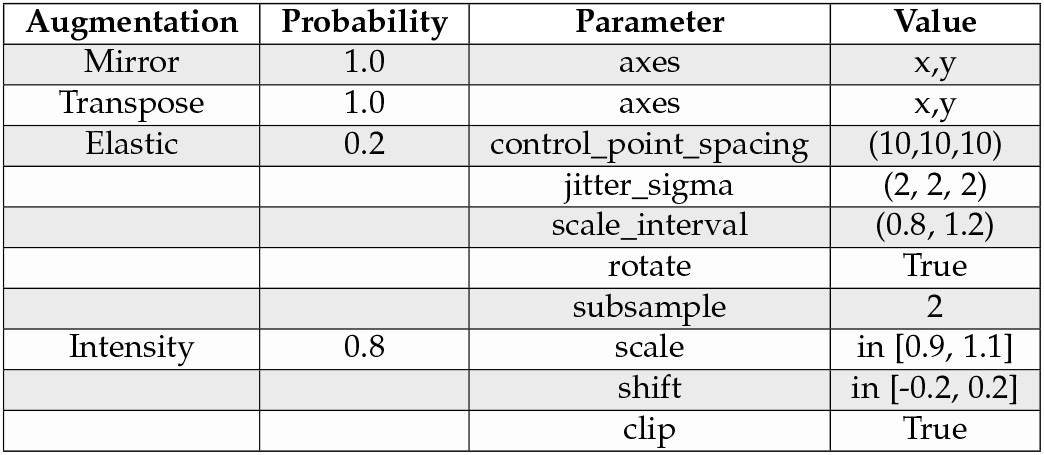
Binary expression network training augmentations.

**Supplementary Table 17:** Mutex watershed parameters.

| Parameter | Value |
| --- | --- |
| bias | [-0.4, -0.7] |
| strides | [(1, 1, 1), (6, 9, 9), (12, 27, 27)] |
| randomized strides | True |
| remove debris | 64 |

**Supplementary Table 18:** Segmentation evaluation results on largest RoI.

| Method | NVoI Split (↓) | NVoI Merge (↓) | NVoI Sum (↓) | ERL ($\mu\text{m}$, ↑) | NERL (↑) |
| --- | --- | --- | --- | --- | --- |
| R-GFP | 0.165 | 0.171 | 0.336 | 5.384 | 0.046 |
| R-M | 0.135 | 0.111 | 0.246 | 7.577 | 0.064 |
| R-S | 0.140 | 0.111 | 0.251 | 10.4 | 0.069 |
| E-M | 0.102 | 0.090 | 0.192 | 13.19 | 0.112 |
| E-S | **0.094** | 0.082 | 0.176 | 14.67 | 0.124 |
| E-S+U | 0.108 | **0.055** | **0.163** | **21.28** | **0.180** |

**Supplementary Table 19:** Segmentation evaluation results on second largest RoI.

| Method | Merges needed (↓) | Splits needed (↓) | Merges and splits needed (↓) | Unsplittable fragments (↓) |
| --- | --- | --- | --- | --- |
| R-GFP | 905 | 561 | 1466 | 250 |
| R-M | 724 | 478 | 1202 | 203 |
| R-S | 661 | 456 | 1117 | 209 |
| E-M | 477 | 362 | 839 | 152 |
| E-S | **325** | 311 | **636** | 142 |
| E-S+U | 451 | **239** | 690 | **95** |

**Supplementary Table 20:** Gap crossing evaluation results at ∼75 microns.

| Channels | FP (↓) | FN (↓) | TP (↑) | F1 (↑) |
| --- | --- | --- | --- | --- |
| 3 | 285 | 80 | 30 | 0.225 |
| 6 | 157 | 34 | 76 | 0.443 |
| 9 | 85 | 18 | 92 | 0.612 |
| 12 | 51 | 23 | 87 | 0.694 |
| 15 | 30 | 8 | 102 | 0.801 |
| 18 | **19** | **8** | **102** | **0.841** |

**Supplementary Table 21:** Gap crossing evaluation results at ∼200 microns.

| Channels | FP (↓) | FN (↓) | TP (↑) | F1 (↑) |
| --- | --- | --- | --- | --- |
| 3 | 7292 | 2779 | 297 | 0.045 |
| 6 | 5088 | 1794 | 1282 | 0.174 |
| 9 | 3304 | 1126 | 1950 | 0.293 |
| 12 | 2375 | 892 | 2184 | 0.355 |
| 15 | 1720 | 614 | 2462 | 0.432 |
| 18 | **1336** | **604** | **2472** | **0.464** |

## Footnotes

* https://github.com/JaneliaSciComp/bigstream

† https://github.com/e11bio/voxelflow

‡ https://pytorch.org/

§ U-Net encoder with kernel sizes of 1 that is densely connected from in channels to out channels

¶ https://funkelab.github.io/gunpowder/

|| https://github.com/e11bio/prism_training

** https://github.com/e11bio/volara

†† https://github.com/funkelab/daisy

‡‡ https://ariadne.ai/

## References

Aitken, C. E., Marshall, R. A., & Puglisi, J. D. (2008). An Oxygen Scavenging System for Improvement of Dye Stability in Single-Molecule Fluorescence Experiments [Publisher: Elsevier]. Biophysical Journal, 94(5), 1826–1835. 10.1529/biophysj.107.117689

Bae, J. A., Baptiste, M., Baptiste, M. R., Bishop, C. A., Bodor, A. L., Brittain, D., Brooks, V., Buchanan, J., Bumbarger, D. J., Castro, M. A., Celii, B., Cobos, E., Collman, F., da Costa, N. M., Danskin, B., Dorkenwald, S., Elabbady, L., Fahey, P. G., Fliss, T., … The MICrONS Consortium. (2025). Functional connectomics spanning multiple areas of mouse visual cortex [Publisher: Nature Publishing Group]. Nature, 640(8058), 435–447. 10.1038/s41586-025-08790-w

Begemann, I., & Galic, M. (2016). Correlative Light Electron Microscopy: Connecting Synaptic Structure and Function [Publisher: Frontiers]. Frontiers in Synaptic Neuroscience, 8. 10.3389/fnsyn.2016.00028

Beier, T., Pape, C., Rahaman, N., Prange, T., Berg, S., Bock, D. D., Cardona, A., Knott, G. W., Plaza, S. M., Scheffer, L. K., Koethe, U., Kreshuk, A., & Hamprecht, F. A. (2017). Multicut brings automated neurite segmentation closer to human performance [Publisher: Nature Publishing Group]. Nature Methods, 14(2), 101–102. 10.1038/nmeth.4151

Bogovic, J. A., Hanslovsky, P., Wong, A., & Saalfeld, S. (2016). Robust registration of calcium images by learned contrast synthesis [ISSN: 1945-8452]. 2016 IEEE 13th International Symposium on Biomedical Imaging (ISBI), 1123–1126. 10.1109/ISBI.2016.7493463

Borch Jensen, M., & Marblestone, A. (2021). In vivo Pooled Screening: A Scalable Tool to Study the Complexity of Aging and Age-Related Disease. Frontiers in Aging, 2, 714926. 10.3389/fragi.2021.714926

Cai, D., Cohen, K. B., Luo, T., Lichtman, J. W., & Sanes, J. R. (2013). NEW TOOLS FOR THE BRAINBOW TOOLBOX. Nature methods, 10(6), 540–547. 10.1038/nmeth.2450

Cao, L., Coventry, B., Goreshnik, I., Huang, B., Sheffler, W., Park, J. S., Jude, K. M., Marković, I., Kadam, R. U., Verschueren, K. H. G., Verstraete, K., Walsh, S. T. R., Bennett, N., Phal, A., Yang, A., Kozodoy, L., DeWitt, M., Picton, L., Miller, L., … Baker, D. (2022). Design of protein-binding proteins from the target structure alone [Publisher: Nature Publishing Group]. Nature, 605(7910), 551–560. 10.1038/s41586-022-04654-9

Chen, F., Tillberg, P. W., & Boyden, E. S. (2015). Expansion microscopy [Publisher: American Association for the Advancement of Science]. Science, 347(6221), 543–548. 10.1126/science.1260088

Chen, X., Sun, Y.-C., Zhan, H., Kebschull, J. M., Fischer, S., Matho, K., Huang, Z. J., Gillis, J., & Zador, A. M. (2019). High-Throughput Mapping of Long-Range Neuronal Projection Using In Situ Sequencing [Publisher: Elsevier]. Cell, 179(3), 772–786.e19. 10.1016/j.cell.2019.09.023

Cherubini, E., & Miles, R. (2015). The CA3 region of the hippocampus: How is it? what is it for? how does it do it? [Publisher: Frontiers]. Frontiers in Cellular Neuroscience, 9. 10.3389/fncel.2015.00019

Cordes, T., Vogelsang, J., & Tinnefeld, P. (2009). On the Mechanism of Trolox as Antiblinking and Antibleaching Reagent [Publisher: American Chemical Society]. Journal of the American Chemical Society, 131(14), 5018–5019. 10.1021/ja809117z

Damstra, H. G. J., Passmore, J. B., Serweta, A. K., Koutlas, I., Burute, M., Meye, F. J., Akhmanova, A., & Kapitein, L. C. (2023). GelMap: Intrinsic calibration and deformation mapping for expansion microscopy [Publisher: Nature Publishing Group]. Nature Methods, 20(10), 1573–1580. 10.1038/s41592-023-02001-y

Di Tommaso, P., Chatzou, M., Floden, E. W., Barja, P. P., Palumbo, E., & Notredame, C. (2017). Nextflow enables reproducible computational workflows [Publisher: Nature Publishing Group]. Nature Biotechnology, 35(4), 316–319. 10.1038/nbt.3820

Dorkenwald, S., Matsliah, A., Sterling, A. R., Schlegel, P., Yu, S.-c., McKellar, C. E., Lin, A., Costa, M., Eichler, K., Yin, Y., Silversmith, W., Schneider-Mizell, C., Jordan, C. S., Brittain, D., Halageri, A., Kuehner, K., Ogedengbe, O., Morey, R., Gager, J., … Murthy, M. (2024). Neuronal wiring diagram of an adult brain [Publisher: Nature Publishing Group]. Nature, 634(8032), 124–138. 10.1038/s41586-024-07558-y

Funke, J., Tschopp, F., Grisaitis, W., Sheridan, A., Singh, C., Saalfeld, S., & Turaga, S. C. (2019). Large Scale Image Segmentation with Structured Loss Based Deep Learning for Connectome Reconstruction. IEEE Transactions on Pattern Analysis and Machine Intelligence, 41(7), 1669–1680. 10.1109/TPAMI.2018.2835450

Gao, R., Asano, S. M., Upadhyayula, S., Pisarev, I., Milkie, D. E., Liu, T.-L., Singh, V., Graves, A., Huynh, G. H., Zhao, Y., Bogovic, J., Colonell, J., Ott, C. M., Zugates, C., Tappan, S., Rodriguez, A., Mosaliganti, K. R., Sheu, S.-H., Pasolli, H. A., … Betzig, E. (2019). Cortical column and whole-brain imaging with molecular contrast and nanoscale resolution [Publisher: American Association for the Advancement of Science]. Science, 363(6424), eaau8302. 10.1126/science.aau8302

Glaser, A., Chandrashekar, J., Vasquez, S., Arshadi, C., Javeri, R., Ouellette, N., Jiang, X., Baka, J., Kovacs, G., Woodard, M., Seshamani, S., Cao, K., Clack, N., Recknagel, A., Grim, A., Balaram, P., Turschak, E., Hooper, M., Liddell, A., … Svoboda, K. (2025). Expansion-assisted selective plane illumination microscopy for nanoscale imaging of centimeter-scale tissues [Publisher: eLife Sciences Publications Limited]. eLife, 12. 10.7554/eLife.91979.3

Gonzales, R. B., DeLeon Galvan, C. J., Rangel, Y. M., & Claiborne, B. J. (2001). Distribution of thorny excrescences on ca3 pyramidal neurons in the rat hippocampus. Journal of Comparative Neurology, 430(3), 357–368.

Goodwin, D. R., Vaughan, A., Leible, D., Alon, S., Henry, G. L., Cheng, A., Chen, X., Zhang, R., Xue, A. G., Wassie, A. T., Sinha, A., Bando, Y., Kajita, A., Marblestone, A. H., Zador, A. M., Boyden, E. S., Church, G. M., & Kohman, R. E. (2022, July). Expansion Sequencing of RNA Barcoded Neurons in the Mammalian Brain: Progress and Implications for Molecularly Annotated Connectomics [Pages: 2022.07.31.502046 Section: New Results]. 10.1101/2022.07.31.502046

Gut, G., Herrmann, M. D., & Pelkmans, L. (2018). Multiplexed protein maps link subcellular organization to cellular states [Publisher: American Association for the Advancement of Science]. Science, 361(6401), eaar7042. 10.1126/science.aar7042

Hardingham, G. E., & Bading, H. (2010). Synaptic versus extrasynaptic NMDA receptor signalling: Implications for neurodegenerative disorders [Publisher: Nature Publishing Group]. Nature Reviews Neuroscience, 11(10), 682–696. 10.1038/nrn2911

Herdly, L., Tinning, P. W., Geiser, A., Taylor, H., Gould, G. W., & van de Linde, S. (2023). Benchmarking Thiolate-Driven Photoswitching of Cyanine Dyes. The Journal of Physical Chemistry. B, 127(3), 732–741. 10.1021/acs.jpcb.2c06872

Hörl, D., Rojas Rusak, F., Preusser, F., Tillberg, P., Randel, N., Chhetri, R. K., Cardona, A., Keller, P. J., Harz, H., Leonhardt, H., Treier, M., & Preibisch, S. (2019). BigStitcher: Reconstructing high-resolution image datasets of cleared and expanded samples [Publisher: Nature Publishing Group]. Nature Methods, 16(9), 870–874. 10.1038/s41592-019-0501-0

Hunker, A. C., Wirthlin, M. E., Gill, G., Johansen, N. J., Hooper, M., Omstead, V., Vargas, S., Lerma, M. N., Taskin, N., Weed, N., Laird, W. D., Bishaw, Y. M., Bendrick, J. L., Gore, B. B., Ben-Simon, Y., Opitz-Araya, X., Martinez, R. A., Way, S. W., Thyagarajan, B., … Ting, J. T. (2025). Enhancer AAV toolbox for accessing and perturbing striatal cell types and circuits [Publisher: Elsevier]. Neuron, 113(10), 1507–1524.e17. 10.1016/j.neuron.2025.04.035

Januszewski, M., Kornfeld, J., Li, P. H., Pope, A., Blakely, T., Lindsey, L., Maitin-Shepard, J., Tyka, M., Denk, W., & Jain, V. (2018). High-precision automated reconstruction of neurons with flood-filling networks [Publisher: Nature Publishing Group]. Nature Methods, 15(8), 605–610. 10.1038/s41592-018-0049-4

Januszewski, M., Templier, T., Hayworth, K., Peale, D., & Hess, H. (2025, May). Accelerating Neuron Reconstruction with PATHFINDER [Pages: 2025.05.16.654254 Section: New Results]. 10.1101/2025.05.16.654254

Kebschull, J. M., Silva, P. G. d., Reid, A. P., Peikon, I. D., Albeanu, D. F., & Zador, A. M. (2016). High-Throughput Mapping of Single-Neuron Projections by Sequencing of Barcoded RNA [Publisher: Elsevier]. Neuron, 91(5), 975–987. 10.1016/j.neuron.2016.07.036

Kingsley, K., Carroll, K., Huff, J. L., & Plopper, G. E. (2001). Photobleaching of Arterial Autofluorescence for Immunofluorescence Applications [Publisher: Taylor & Francis]. BioTechniques, 30(4), 794–797. 10.2144/01304st05

Klimas, A., Gallagher, B. R., Wijesekara, P., Fekir, S., DiBernardo, E. F., Cheng, Z., Stolz, D. B., Cambi, F., Watkins, S. C., Brody, S. L., Horani, A., Barth, A. L., Moore, C. I., Ren, X., & Zhao, Y. (2023). Magnify is a universal molecular anchoring strategy for expansion microscopy [Number: 6 Publisher: Nature Publishing Group]. Nature Biotechnology, 41(6), 858–869. 10.1038/s41587-022-01546-1

Konno, K., & Watanabe, M. (2021). Immunohistochemistry for Ion Channels and Their Interacting Molecules: Tips for Improving Antibody Accessibility. In R. Lujan & F. Ciruela (Eds.), Receptor and Ion Channel Detection in the Brain (pp. 191–199). Springer US. 10.1007/978-1-0716-1522-514

Konno, K., Yamasaki, M., Miyazaki, T., & Watanabe, M. (2023). Glyoxal fixation: An approach to solve immunohistochemical problem in neuroscience research [Publisher: American Association for the Advancement of Science]. Science Advances, 9(28), eadf7084. 10.1126/sciadv.adf7084

Kornfeld, J. M. R. (2025). Brain tissue artificially expanded to show how neurons wire together [Bandiera abtest: a Cg type: News And Views Publisher: Nature Publishing Group Subject term: Brain, Neuroscience, Microscopy, Technology]. Nature, 642(8067), 306–307. 10.1038/d41586-025-01338-y

Krull, A., Buchholz, T.-O., & Jug, F. (2019). Noise2Void - Learning Denoising From Single Noisy Images [ISSN: 2575-7075]. 2019 IEEE/CVF Conference onComputer Vision and Pattern Recognition (CVPR), 2124–2132. 10.1109/CVPR.2019.00223

Kudo, T., Lane, K., & Covert, M. W. (2022). A multiplexed epitope barcoding strategy that enables dynamic cellular phenotypic screens. Cell Systems, 13(5), 376–387.e8. 10.1016/j.cels.2022.02.006

Lam, S. S., Martell, J. D., Kamer, K. J., Deerinck, T. J., Ellisman, M. H., Mootha, V. K., & Ting, A. Y. (2015). Directed evolution of APEX2 for electron microscopy and proximity labeling. Nature Methods, 12(1), 51–54. 10.1038/nmeth.3179

Lee, K., Lu, R., Luther, K., & Seung, H. S. (2021). Learning and Segmenting Dense Voxel Embeddings for 3D Neuron Reconstruction. IEEE Transactions on Medical Imaging, 40(12), 3801–3811. 10.1109/TMI.2021.3097826

Lee, K., Zung, J., Li, P., Jain, V., & Seung, H. S. (2017, May). Superhuman Accuracy on the SNEMI3D Connectomics Challenge [arXiv:1706.00120 [cs]]. 10.48550/arXiv.1706.00120

Leiwe, M. N., Fujimoto, S., Baba, T., Moriyasu, D., Saha, B., Sakaguchi, R., Inagaki, S., & Imai, T. (2024). Automated neuronal reconstruction with supermulticolour Tetbow labelling and threshold-based clustering of colour hues [Publisher: Nature Publishing Group]. Nature Communications, 15(1), 5279. 10.1038/s41467-024-49455-y

Lepeta, K., Lourenco, M. V., Schweitzer, B. C., Martino Adami, P. V., Banerjee, P., Catuara-Solarz, S., de La Fuente Revenga, M., Guillem, A. M., Haidar, M., Ijomone, O. M., Nadorp, B., Qi, L., Perera, N. D., Refsgaard, L. K., Reid, K. M., Sabbar, M., Sahoo, A., Schaefer, N., Sheean, R. K., … Seidenbecher, C. (2016). Synaptopathies: Synaptic dysfunction in neurological disorders - A review from students to students. Journal of Neurochemistry, 138(6), 785–805. 10.1111/jnc.13713

Li, Y., Walker, L. A., Zhao, Y., Edwards, E. M., Michki, N. S., Cheng, H. P. J., Ghazzi, M., Chen, T. Y., Chen, M., Roossien, D. H., & Cai, D. (2021). Bitbow Enables Highly Efficient Neuronal Lineage Tracing and Morphology Reconstruction in Single Drosophila Brains [Publisher: Frontiers]. Frontiers in Neural Circuits, 15. 10.3389/fncir.2021.732183

Livet, J., Weissman, T. A., Kang, H., Draft, R. W., Lu, J., Bennis, R. A., Sanes, J. R., & Lichtman, J. W. (2007). Transgenic strategies for combinatorial expression of fluorescent proteins in the nervous system [Publisher: Nature Publishing Group]. Nature, 450(7166), 56–62. 10.1038/nature06293

Lorincz, A., & Nusser, Z. (2008). Cell-Type-Dependent Molecular Composition of the Axon Initial Segment. The Journal of Neuroscience, 28(53), 14329–14340. 10.1523/JNEUROSCI.4833-08.2008

Loulier, K., Barry, R., Mahou, P., Le Franc, Y., Supatto, W., Matho, K. S., Ieng, S., Fouquet, S., Dupin, E., Benosman, R., Chèdotal, A., Beaurepaire, E., Morin, X., & Livet, J. (2014). Multiplex Cell and Lineage Tracking with Combinatorial Labels. Neuron, 81(3), 505–520. 10.1016/j.neuron.2013.12.016

McAuliffe, J. J., Bronson, S. L., Hester, M. S., Murphy, B. L., Dahlquist-Topalá, R., Richards, D. A., & Danzer, S. C. (2011). Altered patterning of dentate granule cell mossy fiber inputs onto ca3 pyramidal cells in limbic epilepsy. Hippocampus, 21(1), 93–107.

Megías, M., Emri, Z., Freund, T. F., & Gulyás, A. I. (2001). Total number and distribution of inhibitory and excitatory synapses on hippocampal CA1 pyramidal cells. Neuroscience, 102(3), 527–540. 10.1016/s0306-4522(00)00496-6

Meilă, M. (2007). Comparing clusterings—an information based distance. Journal of Multivariate Analysis, 98(5), 873–895. 10.1016/j.jmva.2006.11.013

Melone, M., Burette, A., & Weinberg, R. J. (2005). Light microscopic identification and immunocytochemical characterization of glutamatergic synapses in brain sections. The Journal of Comparative Neurology, 492(4), 495–509. 10.1002/cne.20743

Michalska, J. M., Lyudchik, J., Velicky, P., Štefaničková, H., Watson, J. F., Cenameri, A., Sommer, C., Amberg, N., Venturino, A., Roessler, K., Czech, T., Höftberger, R., Siegert, S., Novarino, G., Jonas, P., & Danzl, J. G. (2024). Imaging brain tissue architecture across millimeter to nanometer scales [Publisher: Nature Publishing Group]. Nature Biotechnology, 42(7), 1051–1064. 10.1038/s41587-023-01911-8

M’Saad, O., Kasula, R., Kondratiuk, I., Kidd, P., Falahati, H., Gentile, J. E., Niescier, R. F., Watters, K., Sterner, R. C., Lee, S., Liu, X., Camilli, P. D., Rothman, J. E., Koleske, A. J., Biederer, T., & Bewersdorf, J. (2022, April). All-optical visualization of specific molecules in the ultrastructural context of brain tissue [Pages: 2022.04.04.486901 Section: New Results]. 10.1101/2022.04.04.486901

Nguyen, T., Malin-Mayor, C., Patton, W., & Funke, J. (2022). Daisy: Block-wise task dependencies for luigi. (Version 1.0). https://github.com/funkelab/daisy

O’Rourke, N. A., Weiler, N. C., Micheva, K. D., & Smith, S. J. (2012). Deep molecular diversity of mammalian synapses: Why it matters and how to measure it [Publisher: Nature Publishing Group]. Nature Reviews Neuroscience, 13(6), 365–379. 10.1038/nrn3170

Patton, W., & Sheridan, A. (2025). Volara: Block-wise operations for large volumetric datasets (Version 1.0). https://github.com/e11bio/volara

Preibisch, S., Saalfeld, S., Rohlfing, T., & Tomancak, P. (2009). Bead-based mosaicing of single plane illumination microscopy images using geometric local descriptor matching. Medical Imaging 2009: Image Processing, 7259, 926–935.

Rebola, N., Carta, M., & Mulle, C. (2017). Operation and plasticity of hippocampal CA3 circuits: Implications for memory encoding [Publisher: Nature Publishing Group]. Nature Reviews Neuroscience, 18(4), 208–220. 10.1038/nrn.2017.10

Ronneberger, O., Fischer, P., & Brox, T. (2015, May). U-Net: Convolutional Networks for Biomedical Image Segmentation [arXiv:1505.04597 [cs]]. 10.48550/arXiv.1505.04597

Rovira-Clavè, X., Jiang, S., Bai, Y., Zhu, B., Barlow, G., Bhate, S., Coskun, A. F., Han, G., Ho, C.-M. K., Hitzman, C., Chen, S.-Y., Bava, F.-A., & Nolan, G. P. (2021). Subcellular localization of biomolecules and drug distribution by high-definition ion beam imaging [Publisher: Nature Publishing Group]. Nature Communications, 12(1), 4628. 10.1038/s41467-021-24822-1

Saka, S. K., Wang, Y., Kishi, J. Y., Zhu, A., Zeng, Y., Xie, W., Kirli, K., Yapp, C., Cicconet, M., Beliveau, B. J., Lapan, S. W., Yin, S., Lin, M., Boyden, E. S., Kaeser, P. S., Pihan, G., Church, G. M., & Yin, P. (2019). Immuno-SABER enables highly multiplexed and amplified protein imaging in tissues [Publisher: Nature Publishing Group]. Nature Biotechnology, 37(9), 1080–1090. 10.1038/s41587-019-0207-y

Sakaguchi, R., Leiwe, M. N., & Imai, T. (2018). Bright multicolor labeling of neuronal circuits with fluorescent proteins and chemical tags. Elife, 7. 10.7554/eLife.40350

Sammons, R. P., Vezir, M., Moreno-Velasquez, L., Cano, G., Orlando, M., Sievers, M., Grasso, E., Metodieva, V. D., Kempter, R., Schmidt, H., & Schmitz, D. (2024). Structure and function of the hippocampal CA3 module [Publisher: Proceedings of the National Academy of Sciences]. Proceedings of the National Academy of Sciences, 121(6), e2312281120. 10.1073/pnas.2312281120

Santuy, A., Tomás-Roca, L., Rodríguez, J.-R., González-Soriano, J., Zhu, F., Qiu, Z., Grant, S. G. N., De-Felipe, J., & Merchan-Perez, A. (2020). Estimation of the number of synapses in the hippocampus and brain-wide by volume electron microscopy and genetic labeling [Publisher: Nature Publishing Group]. Scientific Reports, 10(1), 14014. 10.1038/s41598-020-70859-5

Sato, M., Bitter, I., Bender, M., Kaufman, A., & Nakajima, M. (2000). TEASAR: Tree-structure extraction algorithm for accurate and robust skeletons. Proceedings the Eighth Pacific Conference on Computer Graphics and Applications, 281–449. 10.1109/PCCGA.2000.883951

Scaling up connectomics — Reports. (2023, June). Retrieved August 5, 2025, from https://wellcome.org/reports/scaling-connectomics

Schindelin, J., Arganda-Carreras, I., Frise, E., Kaynig, V., Longair, M., Pietzsch, T., Preibisch, S., Rueden, C., Saalfeld, S., Schmid, B., Tinevez, J.-Y., White, D. J., Hartenstein, V., Eliceiri, K., Tomancak, P., & Cardona, A. (2012). Fiji: An open-source platform for biological-image analysis [Publisher: Nature Publishing Group]. Nature Methods, 9(7), 676–682. 10.1038/nmeth.2019

Schneider Gasser, E. M., Straub, C. J., Panzanelli, P., Weinmann, O., Sassoe-Pognetto, M., & Fritschy, J.-M. (2006). Immunofluorescence in brain sections: Simultaneous detection of presynaptic and postsynaptic proteins in identified neurons [Publisher: Nature Publishing Group]. Nature Protocols, 1(4), 1887–1897. 10.1038/nprot.2006.265

Schneider-Mizell, C. M., Bodor, A. L., Brittain, D., Buchanan, J., Bumbarger, D. J., Elabbady, L., Gamlin, C., Kapner, D., Kinn, S., Mahalingam, G., Seshamani, S., Suckow, S., Takeno, M., Torres, R., Yin, W., Dorkenwald, S., Bae, J. A., Castro, M. A., Halageri, A., … da Costa, N. M. (2025). Inhibitory specificity from a connectomic census of mouse visual cortex [Publisher: Nature Publishing Group]. Nature, 640(8058), 448–458. 10.1038/s41586-024-07780-8

Schwarzkopf, M., Liu, M. C., Schulte, S. J., Ives, R., Husain, N., Choi, H. M. T., & Pierce, N. A. (2021). Hybridization chain reaction enables a unified approach to multiplexed, quantitative, high-resolution immunohistochemistry and in situ hybridization. Development (Cambridge, England), 148(22), dev199847. 10.1242/dev.199847

Shapson-Coe, A., Januszewski, M., Berger, D. R., Pope, A., Wu, Y., Blakely, T., Schalek, R. L., Li, P. H., Wang, S., Maitin-Shepard, J., Karlupia, N., Dorkenwald, S., Sjostedt, E., Leavitt, L., Lee, D., Troidl, J., Collman, F., Bailey, L., Fitzmaurice, A., … Lichtman, J. W. (2024). A petavoxel fragment of human cerebral cortex reconstructed at nanoscale resolution [Publisher: American Association for the Advancement of Science]. Science, 384(6696), eadk4858. 10.1126/science.adk4858

Shen, F. Y., Harrington, M. M., Walker, L. A., Cheng, H. P. J., Boyden, E. S., & Cai, D. (2020). Light microscopy based approach for mapping connectivity with molecular specificity [Publisher: Nature Publishing Group]. Nature Communications, 11(1), 4632. 10.1038/s41467-020-18422-8

Sheridan, A., Nguyen, T. M., Deb, D., Lee, W.-C. A., Saalfeld, S., Turaga, S. C., Manor, U., & Funke, J. (2022). Local shape descriptors for neuron segmentation [Publisher: Nature Publishing Group]. Nature Methods, 20(2), 295–303. 10.1038/s41592-022-01711-z

Silversmith, W., Bae, J. A., Li, P. H., & Wilson, A. (2021, September). Kimimaro: Skeletonize densely labeled 3D image segmentations. 10.5281/zenodo.5539913

Tavakoli, M. R., Lyudchik, J., Januszewski, M., Vistunou, V., Agudelo Dueñas, N., Vorlaufer, J., Sommer, C., Kreuzinger, C., Oliveira, B., Cenameri, A., Novarino, G., Jain, V., & Danzl, J. G. (2025). Lightmicroscopy-based connectomic reconstruction of mammalian brain tissue [Publisher: Nature Publishing Group]. Nature, 642(8067), 398–410. 10.1038/s41586-025-08985-1

Troidl, J., Knittel, J., Li, W., Zhan, F., Pfister, H., & Turaga, S. (2025, June). Global Neuron Shape Reasoning with Point Affinity Transformers [Pages: 2024.11.24.625067 Section: New Results]. 10.1101/2024.11.24.625067

Turaga, S. C., Murray, J. F., Jain, V., Roth, F., Helmstaedter, M., Briggman, K., Denk, W., & Seung, H. S. (2010). Convolutional Networks Can Learn to Generate Affinity Graphs for Image Segmentation. Neural Computation, 22(2), 511–538. 10.1162/neco.2009.10-08-881

Viswanathan, S., Williams, M. E., Bloss, E. B., Stasevich, T. J., Speer, C. M., Nern, A., Pfeiffer, B. D., Hooks, B. M., Li, W.-P., English, B. P., Tian, T., Henry, G. L., Macklin, J. J., Patel, R., Gerfen, C. R., Zhuang, X., Wang, Y., Rubin, G. M., & Looger, L. L. (2015). High-performance probes for light and electron microscopy [Publisher: Nature Publishing Group]. Nature Methods, 12(6), 568–576. 10.1038/nmeth.3365

Vosbein, P., Paredes Vergara, P., Huang, D. T., & Thomson, A. R. (2024). AlphaFold Ensemble Competition Screens Enable Peptide Binder Design with Single-Residue Sensitivity. ACS Chemical Biology, 19(10), 2198–2205. 10.1021/acschembio.4c00418

Wang, T., & Isola, P. (2020). Understanding Contrastive Representation Learning through Alignment and Uniformity on the Hypersphere [ISSN: 2640-3498]. Proceedings of the 37th International Conference on Machine Learning, 9929–9939. Retrieved August 5, 2025, from https://proceedings.mlr.press/v119/wang20k.html

Weigert, M., Schmidt, U., Boothe, T., Müller, A., Dibrov, A., Jain, A., Wilhelm, B., Schmidt, D., Broaddus, C., Culley, S., Rocha-Martins, M., Segovia-Miranda, F., Norden, C., Henriques, R., Zerial, M., Solimena, M., Rink, J., Tomancak, P., Royer, L., … Myers, E. W. (2018). Content-aware image restoration: Pushing the limits of fluorescence microscopy [Publisher: Nature Publishing Group]. Nature Methods, 15(12), 1090–1097. 10.1038/s41592-018-0216-7

Winnubst, J., Bas, E., Ferreira, T. A., Wu, Z., Economo, M. N., Edson, P., Arthur, B. J., Bruns, C., Rokicki, K., Schauder, D., Olbris, D. J., Murphy, S. D., Ackerman, D. G., Arshadi, C., Baldwin, P., Blake, R., Elsayed, A., Hasan, M., Ramirez, D., … Chandrashekar, J. (2019). Reconstruction of 1,000 Projection Neurons Reveals New Cell Types and Organization of Long-Range Connectivity in the Mouse Brain [Publisher: Elsevier]. Cell, 179(1), 268–281.e13. 10.1016/j.cell.2019.07.042

Wolf, S., Pape, C., Bailoni, A., Rahaman, N., Kreshuk, A., Kothe, U., & Hamprecht, F. (2018). The Mutex Watershed: Efficient, Parameter-Free Image Partitioning, 546–562. Retrieved August 5, 2025, from https://openaccess.thecvf.com/content_ECCV2018/html/Steffen_Wolf_The_Mutex_Watershed_ECCV_2018_paper.html

Wroblewska, A., Dhainaut, M., Ben-Zvi, B., Rose, S. A., Park, E. S., Amir, E.-A. D., Bektesevic, A., Baccarini, A., Merad, M., Rahman, A. H., & Brown, B. D. (2018). Protein Barcodes Enable High-Dimensional Single-Cell CRISPR Screens. Cell, 175(4), 1141–1155.e16. 10.1016/j.cell.2018.09.022

Yuan, L., Chen, X., Zhan, H., Henry, G. L., & Zador, A. M. (2024). Massive multiplexing of spatially resolved single neuron projections with axonal BARseq [Publisher: Nature Publishing Group]. Nature Communications, 15(1), 8371. 10.1038/s41467-024-52756-x

Zheng, Z., Park, C., Hammerschmith, E. W., Lu, R., Yu, S.-C., Sorek, M., Silverman, B., Jordan, C. S., Sterling, A. R., Silversmith, W. M., Collman, F., Seung, H. S., & Tank, D. W. (2025, July). Connectomic reconstruction from hippocampal CA3 reveals spatially graded mossy fiber inputs and selective feedforward inhibition to pyramidal cells [ISSN: 2692-8205 Pages: 2025.07.09.663979 Section: New Results]. 10.1101/2025.07.09.663979

## Supplementary References

Macrina, T., Lee, K., Lu, R., Turner, N. L., Wu, J., Popovych, S., Silversmith, W., Kemnitz, N., Bae, J. A., Castro, M. A., Dorkenwald, S., Halageri, A., Jia, Z., Jordan, C., Li, K., Mitchell, E., Mondal, S. S., Mu, S., Nehoran, B., … Seung, H. S. (2021, August). Petascale neural circuit reconstruction: Automated methods [Pages: 2021.08.04.455162 Section: New Results]. 10.1101/2021.08.04.455162

van der Walt, S., Schönberger, J. L., Nunez-Iglesias, J., Boulogne, F., Warner, J. D., Yager, N., Gouillart, E., Yu, T., & the scikit-image contributors. (2014). Scikit-image: Image processing in Python. PeerJ, 2, e453. 10.7717/peerj.453

Virtanen, P., Gommers, R., Oliphant, T. E., Haberland, M., Reddy, T., Cournapeau, D., Burovski, E., Peterson, P., Weckesser, W., Bright, J., van der Walt, S. J., Brett, M., Wilson, J., Millman, K. J., Mayorov, N., Nelson, A. R. J., Jones, E., Kern, R., Larson, E., … SciPy 1.0 Contributors. (2020). SciPy 1.0: Fundamental Algorithms for Scientific Computing in Python. Nature Methods, 17, 261–272. 10.1038/s41592-019-0686-2

